# Navigating the conjugated metabolome

**DOI:** 10.64898/2026.02.06.704496

**Authors:** Shipei Xing, Abubaker Patan, Julius Agongo, Harsha Gouda, Vincent Charron-Lamoureux, Yasin El Abiead, Zhewen Hu, Haoqi Nina Zhao, Ipsita Mohanty, Jasmine Zemlin, Wilhan Donizete Gonçalves Nunes, Lindsey A. Burnett, Mingxun Wang, Dionicio Siegel, Pieter C. Dorrestein

## Abstract

Life’s chemical diversity far exceeds current biochemical maps. While metabolomics has catalogued tens of thousands of small molecules, conjugated metabolites, formed when two or more molecular entities are covalently fused through amidation, esterification, or related chemistries, remain underexplored. These molecules can act as microbial signals, detoxification intermediates, or endogenous regulators. Here, we mined 1.32 billion MS/MS spectra across public metabolomics repositories using reverse spectral searching coupled with delta-mass inference to map conjugation events. We generated structural hypotheses for 24,227,439 MS/MS clusters. From these, we inferred 217,291 substructure pairs with dual spectral support and 3,412,720 candidate conjugates with single-match support. Predictions span host–microbe co-metabolites, diet-derived conjugates, and drug-derived species, including drug-ethanolamine and creatinine conjugates with altered bioactivities. We also uncover a family of steroid-phosphoethanolamine conjugates. Fifty-five conjugates were matched by MS/MS of synthetic standards for this work, with 27 additionally supported by retention time matching in biological samples. Guidance on how to leverage this resource is also provided. Together, these results deliver a pan-repository map of potential conjugation chemistry, establish a resource for structural discovery and MS/MS annotation, and offer a scalable framework to explore the scope and diversity of the conjugated metabolome.

## Main

Over the past decade, a new mode of biological discovery has taken hold. Insight increasingly comes not only from experimental advances but from systematic reanalysis of publicly available omics data. This shift is most evident in sequencing-based fields^1,2^ and, more recently, in proteomics, where computational reanalysis of existing datasets has revealed new regulatory circuits, splice isoforms, and post-translational modifications that were not recognized at the time of data deposition^3–5^. These efforts have substantially expanded our view of gene regulation and cellular networks.

Yet in metabolism, particularly in small molecule and natural product structure discovery, much of this latent richness remains untapped. In LC-MS/MS untargeted metabolomics, the average annotation rate based on MS/MS matching to reference libraries is only 8.1%, and 7.0% for human samples. Closing this gap will require approaches that extend beyond compounds that have already been synthesized or are commercially available. Data-driven strategies that generate structural hypotheses, followed by targeted chemical synthesis for validation^6^, offer a promising path forward.

One relatively accessible but insufficiently explored area is the conjugated metabolome - compounds formed when two or more molecular entities are covalently fused through reactions such as amidation, esterification, ether formation, alkylation, or thioester linkage. Many biologically important molecules arise from such conjugations (e.g., glucuronidation, glutathionylation, and sulfation). Emerging examples, such as the expanded repertoire of bile acid amidates^7^, acyl carnitines, N-acyl lipids^8^, together with compounds readily generated by multiplexed synthesis, point to a much broader chemical landscape. This landscape spans diverse biological and exposure-linked chemistry that may have long escaped annotation.

Here, we systematically mined 1.32 billion public MS/MS spectra deposited in metabolomics repositories of MetaboLights^9^, Metabolomics Workbench^10^ and GNPS/MassIVE^11^ (as of October 2024) to identify MS/MS signatures of metabolite conjugation, focusing on conjugates formed by covalent fusion of two component metabolites. By combining reverse spectral search with mass difference filtering to infer conjugating linkages, we generated 3,630,011 data-driven candidate substructure pairings spanning diverse biochemical classes. These candidates encompass host–microbe co-metabolites, food- and drug-derived conjugates, xenobiotic intermediates, and previously unrecognized endogenous molecules such as steroid-phosphoethanolamine conjugates. From this compendium, we established structure-based hypotheses that were further supported through chemical synthesis, with 28 conjugates validated by MS/MS matching and an additional 27 confirmed by combined MS/MS and retention time agreement.

### Repository-scale exploration of metabolite conjugates

Many metabolites arise through the conjugation of two smaller metabolites. Examples include bile acid conjugates, N-acyl lipids, and glycosylated compounds (**Fig. 1a**). The MS/MS fragmentation of such conjugates is often dominated by one of the two components, although in many cases both fragments are well represented - particularly when each moiety contains substituents that readily undergo fragmentation. These fragment ions can be traced to known substructures by comparing MS/MS spectra to reference libraries using reverse spectral search.

**Fig. 1.**
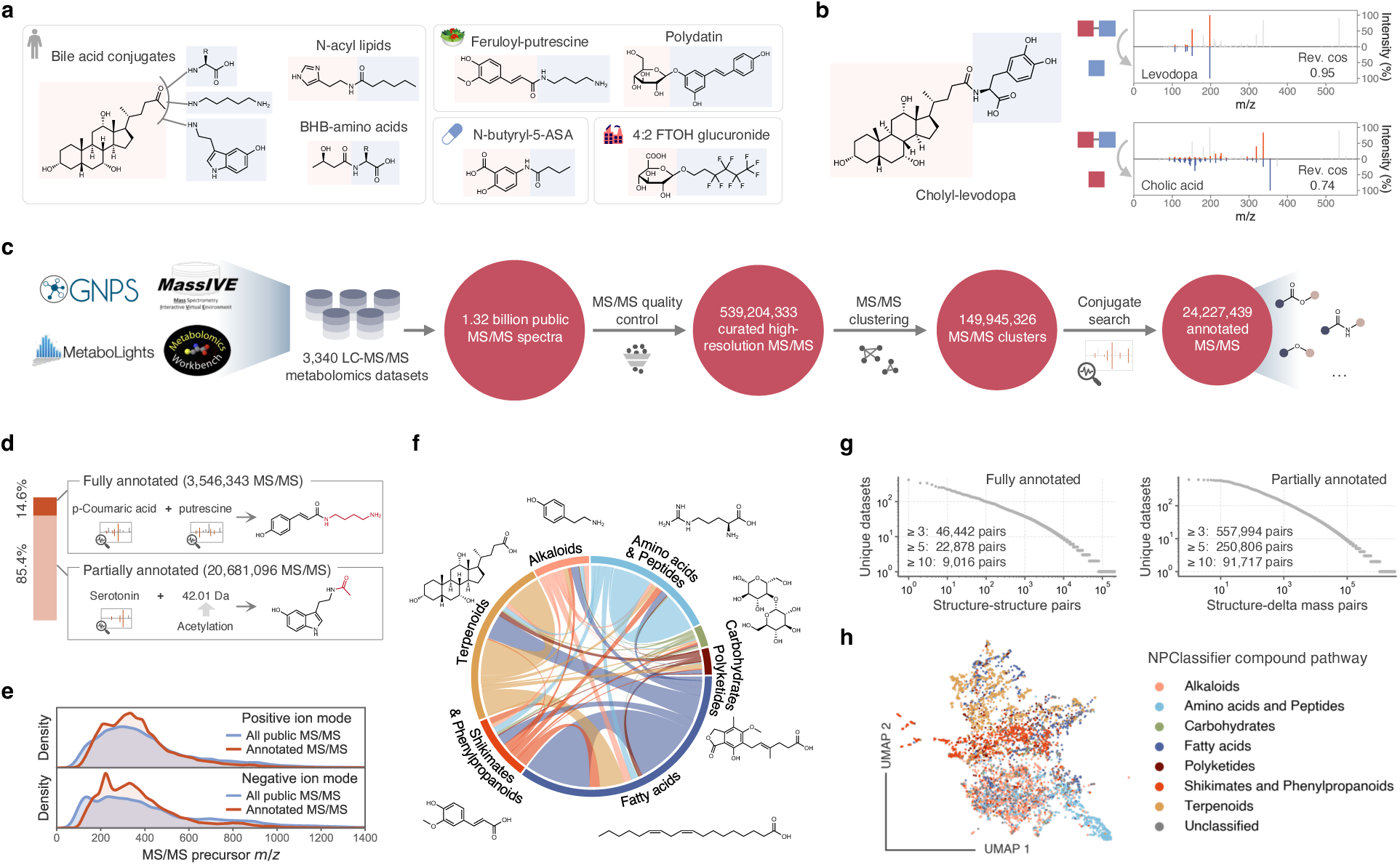
Pan-repository-scale exploration of metabolite conjugates in untargeted metabolomics. **a**) Examples of metabolite conjugates derived from diverse sources, including endogenous metabolites, dietary metabolites, drugs and industrial chemicals. **b**) Representative mirror plot showing both components of a metabolite conjugate annotated via reverse spectral matching. Reference spectra are available in the GNPS library: cholyl-levodopa (CCMSLIB00006581992), levodopa (CCMSLIB00006439802), cholic acid (CCMSLIB00006581904). Rev. cos, reverse cosine. **c**) Schematic workflow of exploration of metabolite conjugates at the repository scale. **d**) Reverse cosine searching generated candidate metabolite conjugate explanations for 24 million MS/MS spectra, of which 14.6% showed dual spectral matches. **e**) Mass distributions of annotated metabolite conjugates in both ionization modes, with most conjugates falling in the *m/z* range of 150 to 600. **f**) Conjugation patterns across compound classes based on NPClassifier pathways. Only metabolite conjugates supported by dual spectral matches are shown. **g**) Annotation frequencies of fully and partially annotated metabolite conjugates across public metabolomics repositories. **h**) UMAP visualization of metabolite conjugate diversity based on delta mass profiles, with features colored by NPClassifier pathway.

Originally introduced in 1975^12^, we have recently created an updated implementation of the reverse spectral search in Python^13^. Unlike the traditional MS/MS similarity metric, which compares all fragment ions in a query spectrum to all ions in a reference, the reverse search evaluates only the ions present in the reference spectrum. This enables more selective, substructure-level matching and avoids penalization from unmatched ions originating from unrelated parts of the conjugate. For example, the MS/MS spectrum of cholyl-levodopa, an amide-linked conjugate of levodopa and cholic acid, yields a reverse cosine score of 0.95 against the MS/MS spectrum of levodopa and 0.74 against that of cholic acid (**Fig. 1b**). This principle forms the foundation of the analysis presented in this work, enabling systematic data-driven hypothesis generation and discovery of the pan-repository conjugated metabolome.

To initiate this large-scale analysis while maintaining traceability to the original raw data through Universal Spectrum Identifiers (USIs) and the Metabolomics Spectrum Resolver^14^, we first filtered the pan-repository datasets for curated high-resolution MS/MS spectra collected on instruments such as Orbitrap, TOF, or FT-ICR (**Fig. 1c**, see Methods). Identical MS/MS spectra were subsequently clustered using Falcon^15^, after which a reverse cosine search was performed against 818,648 reference MS/MS spectra from GNPS, MoNA, MassBank and NIST20 libraries. Out of 149,945,326 MS/MS clusters, 24,227,439 (16.2%) yielded at least one reverse cosine match that met the following criteria: reverse cosine score ≥ 0.8, ≥ 3 matched fragment peaks and ≥ 30% spectral usage. Among these matched spectra, 20,681,096 spectra (85.4%) produced a single reverse cosine match (**Fig. 1d**). The unmatched portion of the spectrum could often be rationalized by a defined atomic mass difference corresponding to a plausible conjugating unit, such as a C_3_H_4_O_2_ addition (delta mass 72.02 Da) indicative of lactate conjugation (+C_3_H_6_O_3_−H_2_O). The remaining 14.6% of spectra showed two valid matches whose combined precursor masses - after accounting for the expected reaction (e.g., amide or ester linkage) - reconstructed the observed precursor mass of the conjugate.

Metabolite conjugation prediction performance varies across chemical classes. To benchmark reverse cosine for substructure annotation, we evaluated performance using a reference MS/MS library comprising 9,156 spectra from 7,575 unique chemical conjugates generated by multiplex synthesis of structurally diverse compounds^16^. False discovery rate (FDR) and recall were assessed across a range of score and peak-matching thresholds (**Extended Data Fig. 1**). As expected, increasing the score or the number of matched peaks improved precision at the cost of recall. To construct a routinely usable MS/MS reference library from reverse cosine results, we applied chemical class-specific thresholds based on observed performance, achieving FDR below 1% wherever possible (**Supplementary Table 1**). For the present study, which emphasizes large-scale molecule discovery, we adopted thresholds of score ≥ 0.8, ≥ 3 matched peaks, and ≥ 30% spectral usage to balance coverage and confidence. Under these criteria, FDRs were typically below 5% for phenylpropanoids and alkaloids, below 10% for amino acids and small peptides, and higher for terpenoids, fatty acids and related classes (**Extended Data Fig. 2**). Additional confidence can be derived from chemical or biological context, such as Mass Spectrometry Query Language (MassQL)^17^ constraints or co-occurrence of predicted conjugates with their corresponding parent molecules within the same data file (see drug examples discussed below).

### Global landscape of metabolite conjugates

Overall, compared with all public MS/MS data, the 24 million annotated conjugate spectra were predominantly distributed between precursor masses of 200 and 500 Da (**Fig. 1e**). Among the 3.5 million spectra with both components annotated, NPClassifier^18^ analysis revealed broad structural diversity (**Fig. 1f**, **Extended Data Fig. 3**). The most frequent pairings involved fatty acids, terpenoids, amino acids, and peptides, with additional occurrences across other structural classes, suggesting that conjugation reactions span both primary and specialized metabolism. To assess the prevalence of these findings across studies, we examined the frequency of recurrent structure pairs in independent datasets (**Fig. 1g**). For 217,291 annotated conjugates with two substructures assigned using reverse cosine, 46,442 unique pairings were detected in at least 3 datasets, and 9,016 pairings were observed in 10 or more, indicating that a core subset of conjugate chemistry recurs across laboratories, sample types, or analytical platforms. Inclusion of partially annotated spectra expanded these relationships by nearly an order of magnitude, highlighting the widespread yet underannotated nature of conjugate-like MS/MS features. Visualization of conjugate diversity using UMAP based on delta mass profiles revealed extensive chemical heterogeneity spanning multiple compound classes and metabolic pathways (**Fig. 1h**).

We next assessed how MS/MS spectra with predicted conjugates are distributed across biological contexts. Using Pan-ReDU^19^, a centralized framework integrating harmonized metadata from public metabolomics datasets, we linked metabolite conjugates to the biological sources in which they were detected. Metabolite conjugates were broadly distributed across biological systems (**Fig. 2a**). The largest intersections were observed in microbial monocultures, humans, plants, and rodents individually, indicating pervasive conjugate formation across distinct biological contexts. Shared conjugates among these systems further point to overlapping metabolic capabilities and cross-domain biochemical connectivity. Within human datasets, annotated conjugates were more prevalent in microbially complex body sites, although cross-study methodological heterogeneity may contribute (**Fig. 2b**). Fecal, urinary, and salivary samples showed higher frequencies of annotated conjugates, whereas low-microbial-burden matrices such as blood and brain contained substantially fewer (**Fig. 2c**). Intersections across human body sites revealed both niche-specific and widely distributed conjugates, indicating that host–microbe interactions, together with xenobiotic exposures, jointly shape the conjugated metabolome. Similar trends were observed in rodents (**Extended Data Fig. 4**).

**Fig. 2.**
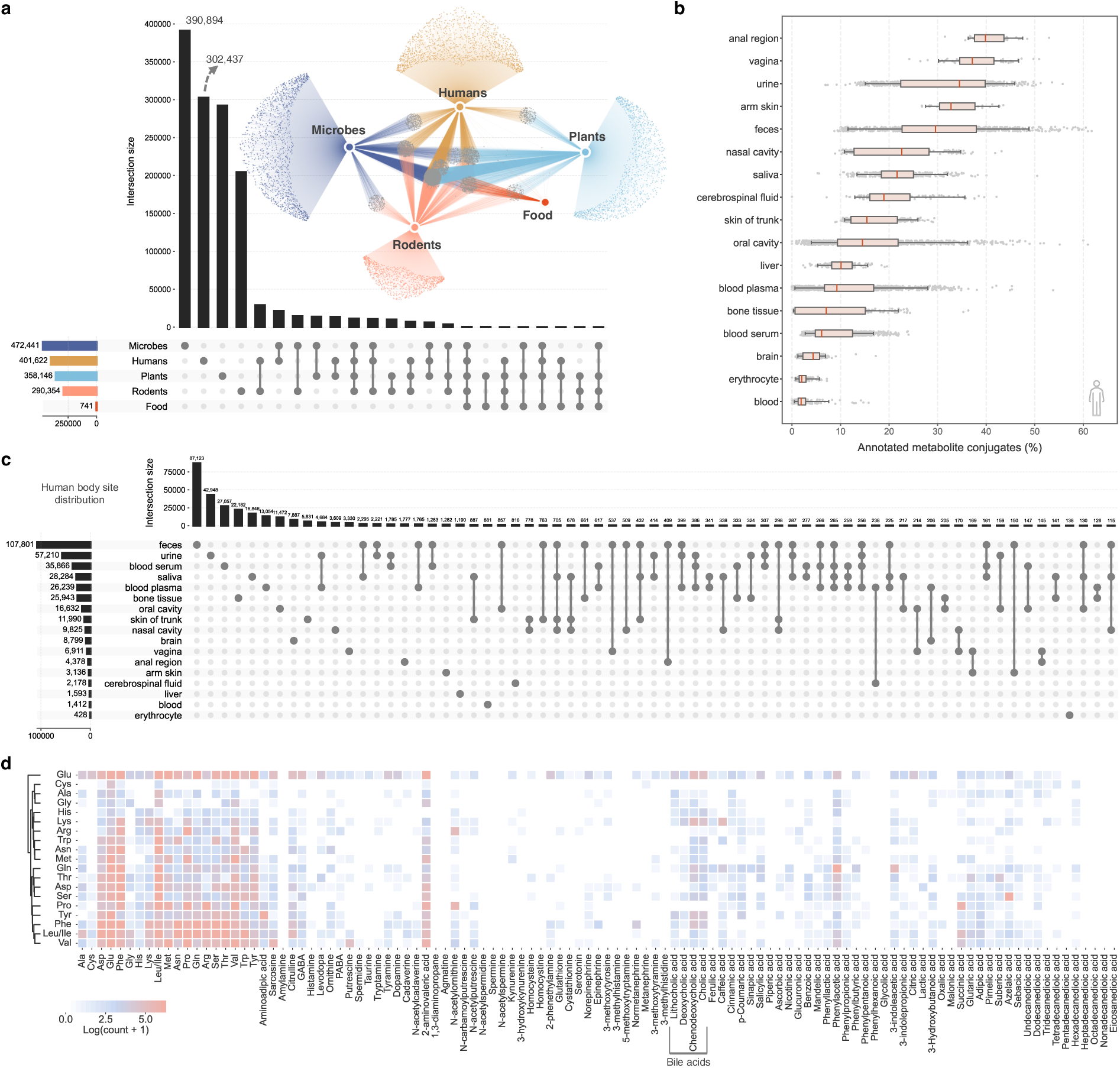
Diverse metabolite conjugate matches and their distributions in humans. **a**) MS/MS of predicted metabolite conjugates are widely distributed across microbial monocultures, humans, rodents, and plant- and food-related data. **b**) In human samples, more metabolite conjugates were annotated in microbially diverse and complex body sites. Interquartile ranges with median values are shown. Bars show the 5th percentile and the 95th percentile. **c**) Distribution of MS/MS of predicted metabolite conjugates across various human body parts. Intersections with sizes of more than 100 are shown. **d**) Heatmap showing representative metabolite names obtained from reverse cosine MS/MS matching that are predicted to conjugate with the 20 canonical proteinogenic amino acids.

### Amino acid conjugation chemistry

Next, we examined conjugation patterns involving the 20 canonical proteinogenic amino acids (**Fig. 2d**). Well-characterized amino acid conjugations, such as amino acid-amino acid linkages (dipeptides) and bile acid conjugates^7,20–22^ with lithocholic acid, deoxycholic acid, chenodeoxycholic acid, and cholic acid, served as internal benchmarks. Beyond these expected relationships, we annotated additional families of aromatic acid-amino acid conjugates, including phenylacetic acid and phenylpropionic acid derivatives, expanding the known chemical space of amino acid conjugation. At the same time, these analyses suggest that many plausible metabolite conjugates are not captured by standard MS/MS matching, particularly when one conjugated partner is small or weakly fragmenting. To rescue such conjugates, we performed delta mass search, which enables recovery of conjugates where one low-mass component - such as lactic acid and other short-chain hydroxy acids - cannot be independently matched during spectral search (**Fig. 3a**). Using this approach, we next focused on lactic acid-, phenylacetic acid-, and phenylpropionic acid-amino acid conjugates, representing chemical families whose diversity and distribution can now be systematically examined.

**Fig. 3.**
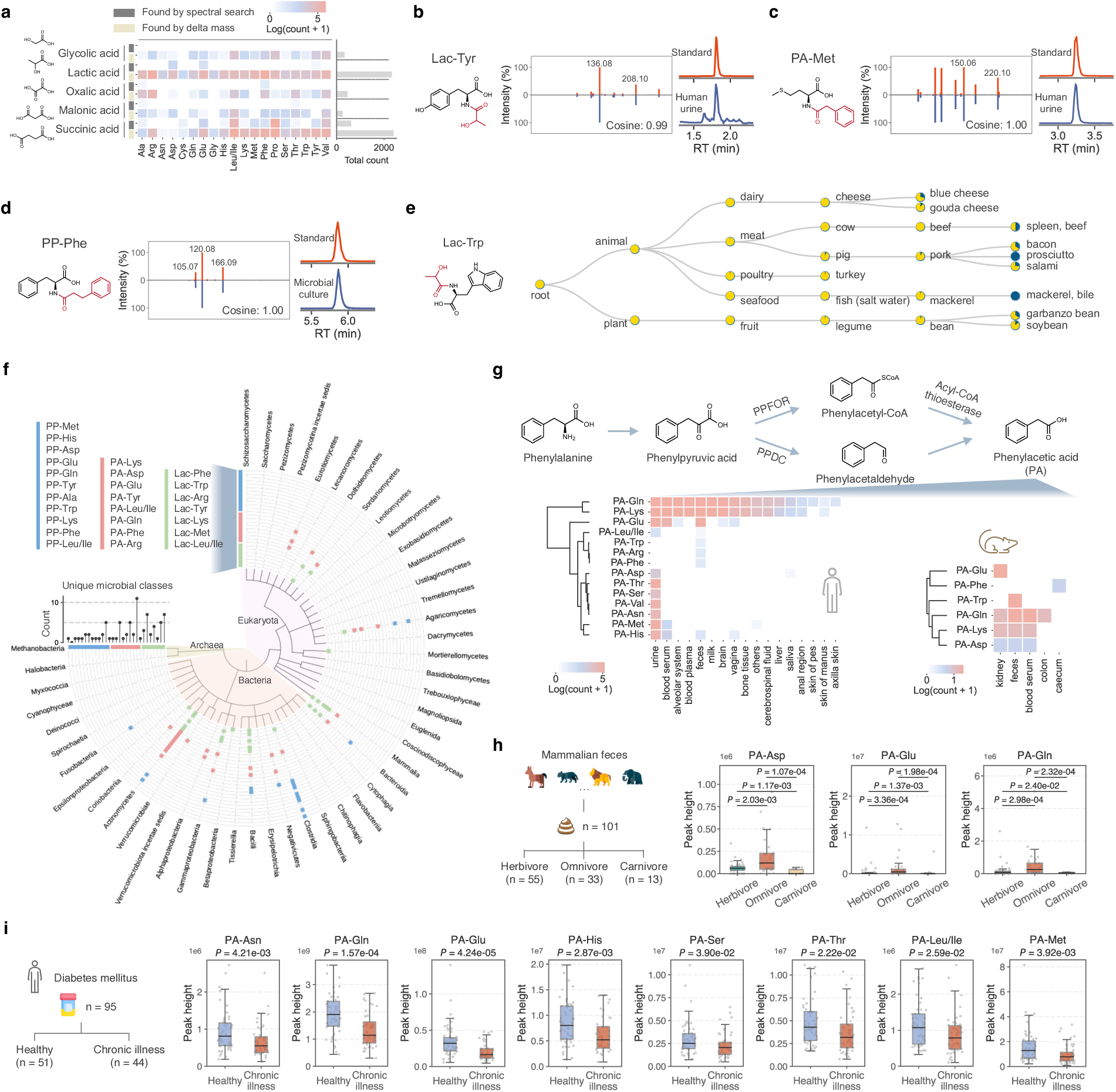
Metabolite conjugates derived from lactic acid (Lac), phenylacetic acid (PA) and phenylpropionic acid (PP). **a**) Small conjugate components, such as lactic acid, can be hard to recognize via spectral search, and can be rescued using delta mass search. **b-d**) Structure confirmation of lactic acid-tyrosine (Lac-Tyr), phenylacetic acid-methionine (PA-Met), and phenylpropionic acid-phenylalanine (PP-Phe) via MS/MS and retention time matching. **e**) Detection of lactic acid-tryptophan (Lac-Trp) across food samples in the public domain. **f**) Microbial class distributions of synthesized Lac-, PA- and PP-amino acid conjugates found by microbeMASST. **g**) Pathways of PA production and body site distributions of PA-amino acid conjugates across human and rodent samples. **h**) PA-amino acid conjugates detected in the mammalian fecal samples from 101 individuals representing 25 species (MSV000086131). Herbivore, n = 55; Omnivore, n = 33; Carnivore, n = 13. Two-sided Mann-Whitney *U* test applied. **i**) Significantly regulated PA-amino acid conjugates in the diabetes mellitus dataset (human urine, MSV000082261). Healthy, n = 51; Chronic illness, n = 44. Two-sided Mann-Whitney *U* test applied. For all boxplots, interquartile ranges (IQR) with median values are shown. Bars show the 75th percentile + 1.5 × IQR and the 25th percentile − 1.5 × IQR.

To support these assignments and verify their existence, we used multiplexed synthesis^6,16^ to generate a library of lactic acid-, phenylacetic acid-, and phenylpropionic acid-amino acid conjugates, yielding reference MS/MS spectra for 54 unique conjugates. Lactic acid-amino acid conjugates, such as lactic acid-phenylalanine, have been reported as endogenous metabolites formed through CNDP2-mediated reverse proteolysis in mammals^23–25^. Phenylacetic acid-glycine, phenylacetic acid-glutamine and phenylacetic acid-ornithine have been described as detoxification products in mammalian metabolism^26^. In contrast, phenylpropionic acid-amino acid conjugates have not undergone comparable systematic investigation beyond the reported glycine conjugate^27^. As these candidate conjugations could be molecules that exist in biological extracts pre-chromatography or form post chromatography within electrospray droplets^28^, retention time matching with standards will be able to distinguish these possibilities. To establish the structural validity of this broader class, we selected representative conjugates spanning multiple scaffolds, lactic acid-tyrosine, phenylacetic acid-methionine, and phenylpropionic acid-phenylalanine, and confirmed their identities by MS/MS and retention time matching between synthetic standards and biological samples (**Fig. 3b-d**, **Extended Data Figs. 5 and 6**). These validations leveraged the MAss Spectrometry Search Tool (MASST)^29,30^ based links to identify accessible or related biological samples, and the synthetic simplicity of the conjugates enabled rapid standard generation, allowing experimental confirmation despite initial reverse spectral matching.

Reference spectra of the synthetic conjugates were searched against the 1.7 billion indexed MS/MS spectra available as of October 2025 in public metabolomics repositories and subjected to reverse metabolomics, to map their prevalence and biological associations across species, diets, and health conditions^6,31^. Lactic acid-amino acid conjugates were broadly detected across humans, rodents, microbes, plants and food samples^6,31–36^ (**Extended Data Figs. 7 and 8a-b**). For example, lactic acid-tryptophan was observed in multiple high protein food sources, including cheese, beef, pork, fish, and beans (**Fig. 3e**). Using microbeMASST^32^, which aggregates more than 60,000 LC-MS/MS data files from microbial monocultures, we further mapped the taxonomic distribution of MS/MS spectra corresponding to the synthesized lactic acid-, phenylacetic acid-, and phenylpropionic acid-amino acid conjugates. These conjugates were detected across a broad range of bacteria and fungi (**Fig. 3f**), with Actinomycetes, Gammaproteobacteria, Bacilli, Clostridia, Bacteroidia and Agaricomycetes most frequently represented.

Phenylacetic acid itself is a well-characterized microbial metabolite derived from phenylalanine via two established gut microbial routes^37^: oxidative decarboxylation catalyzed by phenylpyruvate:ferredoxin oxidoreductase and non-oxidative decarboxylation catalyzed by phenylpyruvate decarboxylase (**Fig. 3g**). Phenylpropionic acid is likewise a common gut microbial product of phenylalanine metabolism^27^ and can also originate from dietary sources. Consistent with these origins, phenylacetic acid-and phenylpropionic acid-amino acid conjugates were detected in human and rodent fecal samples (**Fig. 3g**, **Extended Data Fig. 8c-d**), thus highlighting likely conserved host–microbe co-metabolic processes. Comparative analysis across 101 mammalian fecal samples revealed dietary associations^38^. Omnivores, in that study, exhibited higher levels of phenylacetic acid-aspartic acid, phenylacetic acid-glutamic acid and phenylacetic acid-glutamine conjugates than herbivores or carnivores (**Fig. 3h**). In a separate human urine cohort from individuals with diabetes mellitus, multiple phenylacetic acid-amino acid conjugates, including phenylacetic acid-asparagine, phenylacetic acid-glutamine and phenylacetic acid-glutamic acid, were downregulated in individuals with diabetes compared to controls (**Fig. 3i**, **Extended Data Fig. 8e-f**).

### Steroid-phosphoethanolamine conjugates

Next, we sought to demonstrate how targeted reverse cosine analysis can identify molecules potentially derived from a specific parent compound of interest, leading to the discovery of a new class of steroidal conjugates. As part of a larger effort in the lab to understand microbial processing of dietary molecules, we cultured tomatidine, a cytotoxic steroidal alkaloid aglycone produced by tomatoes, with defined human microbial communities of known composition^39^ (**Fig. 4a**, **Supplementary Table 2**). Using reverse cosine scoring, we identified all MS/MS spectra that were derived from the tomatidine steroidal core. Molecular networking of these spectra revealed a recurrent delta mass of 123.01 Da that, after accounting for water loss during conjugation, corresponds to an exact mass of 141.02 Da; spectra exhibiting this delta mass clustered with tomatidine sulfate in the molecular network (**Fig. 4b**). Formula annotation using BUDDY^40^ and subsequent molecular formula searches in PubChem^41^ identified a molecular formula of C_2_H_8_NO_4_P and that is likely corresponded to phosphoethanolamine (PEA) as the most plausible match, suggesting the presence of tomatidine-PEA. Consistent with this assignment, PEA-containing molecules typically exhibit either a diagnostic fragment ion at *m/z* 142.03 or a neutral loss of 141.02 Da^42^ (**Fig. 4c**). Inspection of its fragmentation pattern also showed a neutral loss corresponding to ethanolamine, indicating that conjugation is more likely to occur via the phosphate moiety to form a phosphate diester rather than conjugation via the amine group of PEA. Reverse metabolomics analysis revealed that the tomatidine-PEA MS/MS spectrum was observed in a tomato seedling dataset^43^ as well as in human fecal samples (**Extended Data Fig. 9a**). Importantly, in the microbial co-culturing experiments, tomatidine-PEA was detected only when tomatidine was supplied as a substrate, directly implicating microbiota-dependent formation (**Fig. 4d**). The structure was confirmed by chemical synthesis, followed by MS/MS and retention time matching, achieving level 1 identification^44^ (**Fig. 4e**).

**Fig. 4.**
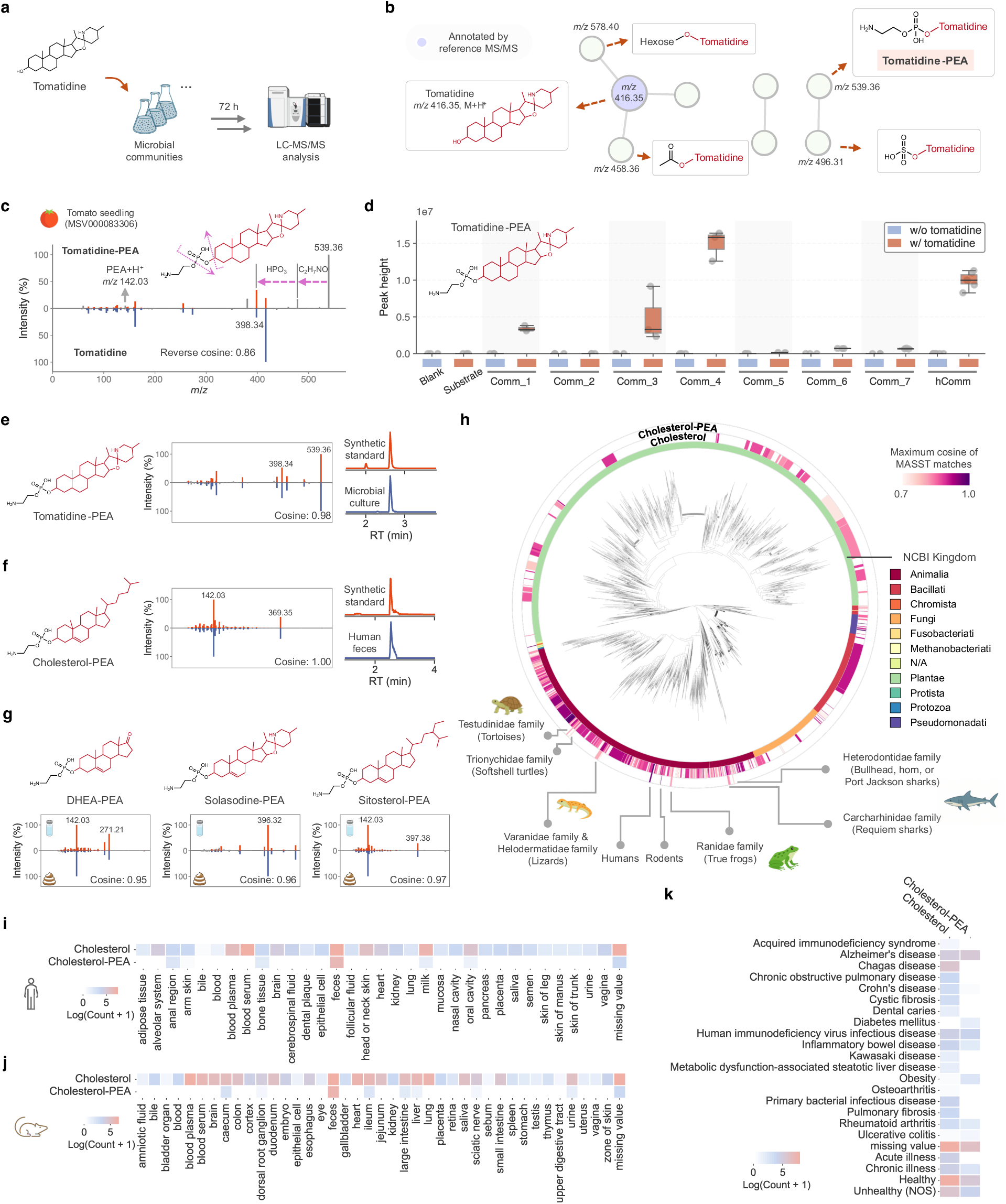
Discovery and verification of the steroid-phosphoethanolamine family. **a**) The microbial community culturing experiment showed that tomatidine-PEA can be made by human gut microbiota. **b**) Molecular networking visualization of predicted tomatidine conjugates. Eight singletons are not shown. **c**) MS/MS matching between tomatidine and tomatidine-PEA, which was observed in a tomato seedling dataset (MSV000083306). **d**) Tomatidine-PEA was formed by synthetic communities, only when tomatidine was added as a substrate. hComm, defined human synthetic microbial community created with 79 bacterial strains. Comm 1 to 7 represent subsets of hComm grouped by growth rates. **e-f**) MS/MS and retention time matching of tomatidine-PEA and cholesterol-PEA. **g**) MS/MS matching of dehydroepiandrosterone-PEA (DHEA-PEA), solasodine-PEA and sitosterol-PEA between synthetic standards and biological samples. **h**) Taxonomic distributions of cholesterol and cholesterol-PEA across public repositories. Family-level MS/MS matching was applied. **i**) Human body site distributions of cholesterol and cholesterol-PEA. The “missing value” indicates that MS/MS matches were obtained in human samples, but the specific sample types were not specified. **j**) Rodent body site distributions of cholesterol and cholesterol-PEA. **k**) Disease associations of cholesterol and cholesterol-PEA in the public domain.

The finding of tomatidine-PEA motivated a broader search for PEA-conjugated steroidal molecules across the pan-repository reverse metabolomics analysis. By filtering the reverse spectral resource for PEA-associated signatures and compounds containing a tetracyclic steroidal core (Methods), we identified 50 candidate steroidal compounds potentially modified with PEA. These candidates included both animal- and plant-derived sterol-like molecules, such as cholesterol, dehydroepiandrosterone (DHEA), sitosterol, and solasodine (**Supplementary Table 3**). To validate these annotations, we synthesized four representative steroid-PEA conjugates: cholesterol-PEA, DHEA-PEA, sitosterol-PEA, and solasodine-PEA. The MS/MS spectra of the synthetic standards matched the corresponding spectra annotated as steroid-PEAs by reverse cosine search, with cosine scores of 0.95 or higher, consistent with level 2 annotation^44^. In addition, owing to sample availability, cholesterol-PEA was further confirmed by retention time matching in human fecal extracts, achieving level 1 identification (**Fig. 4f,g**). Pan-repository searches showed that the MS/MS of DHEA-PEA was detected predominantly in human fecal samples (**Extended Data Fig. 9b**), whereas sitosterol-PEA occurred in both human and rodent feces (**Extended Data Fig. 9c**), consistent with sitosterol’s dietary origin as a major phytosterol abundant in plant-derived foods such as vegetable oil, nuts and grain^45^.

The cholesterol-PEA was particularly notable due to its prevalence, with 11,068 matched MS/MS spectra across 156 independent public datasets. This conjugate was detected in 16 species spanning vertebrates, reptiles, and marine organisms (**Fig. 4h**), indicating cross-species occurrence, while remaining undetected in plant-associated datasets. In human datasets, cholesterol-PEA was observed primarily in data from feces and anal swabs, with additional detections in breast milk, bone tissue, and the oral cavity (**Fig. 4i**). Its enrichment in microbiota-rich and mucosal-associated environments, such as the gut and oral cavity, suggests formation through microbial or host–microbe co-metabolic processes, consistent with the observations of tomatidine-PEA, whereas its presence in milk and bone tissue reflects systemic distribution and infant exposure. Consistent patterns were observed in rodent datasets, where cholesterol-PEA was most prevalent in feces, cecum, ileum, and large intestine (**Fig. 4j**).

Across human studies, cholesterol-PEA was most frequently detected in samples from healthy individuals (**Fig. 4k**). The metabolite was also observed across a broad range of health contexts, including cardiometabolic, neurodegenerative, and inflammatory conditions such as Alzheimer’s disease, type 2 diabetes, human immunodeficiency virus infection, obesity, and inflammatory bowel disease. Notably, cholesterol-PEA spectral matches were less frequent in gut-dysbiotic conditions, including Crohn’s disease and ulcerative colitis, which are characterized by decreased microbiome abundance and altered composition. While conducting this work, we found no prior reports describing direct PEA conjugation of cholesterol, tomatidine, sitosterol, solasodine, or related steroidal cores as a phosphodiester. A recent preprint on microbial metabolism of dietary solanums did independently propose potential PEA additions of tomatidine and solasodine; structural assignment in that study was supported by H/D exchange but lacked validation with authentic standards^46^. Collectively, these observations define a family of steroid-PEA conjugates spanning plant, animal (including human), and microbial systems.

### Drug-derived conjugates in humans

Having established the existence of these previously undescribed PEA conjugates, we next asked whether previously uncharacterized conjugations occur with xenobiotic compounds. Our reverse cosine analysis of MS/MS that originated from human metabolomics datasets revealed numerous predicted drug-derived conjugates, indicating that conjugation chemistry is not confined to endogenous metabolites but also derived from exogenous compounds. We examined the 200 most commonly used drugs in 2022^47^, 175 of which were small molecules. Endogenous molecules that are also administered as drugs (e.g., testosterone or estrogen) were excluded as endogenous production could not be distinguished from pharmaceutical intake. For the remaining compounds, we extracted predicted conjugates from the reverse cosine analysis and further required co-occurrence of each conjugate with its parent drug within the same LC-MS/MS files, acknowledging that this criterion may miss cases of complete drug metabolism. Using this approach, 74 drugs met these criteria, enabling plausible structural assignments. The resulting conjugates encompassed a broad spectrum of putative phase I- and phase II-like transformations, many of which have not been previously documented, including acylation with fatty acids or hydroxy fatty acids, esterification with fatty alcohols, glycosylation, sulfation, and conjugation with sugars or amino acids (**Fig. 5a**).

**Fig. 5.**
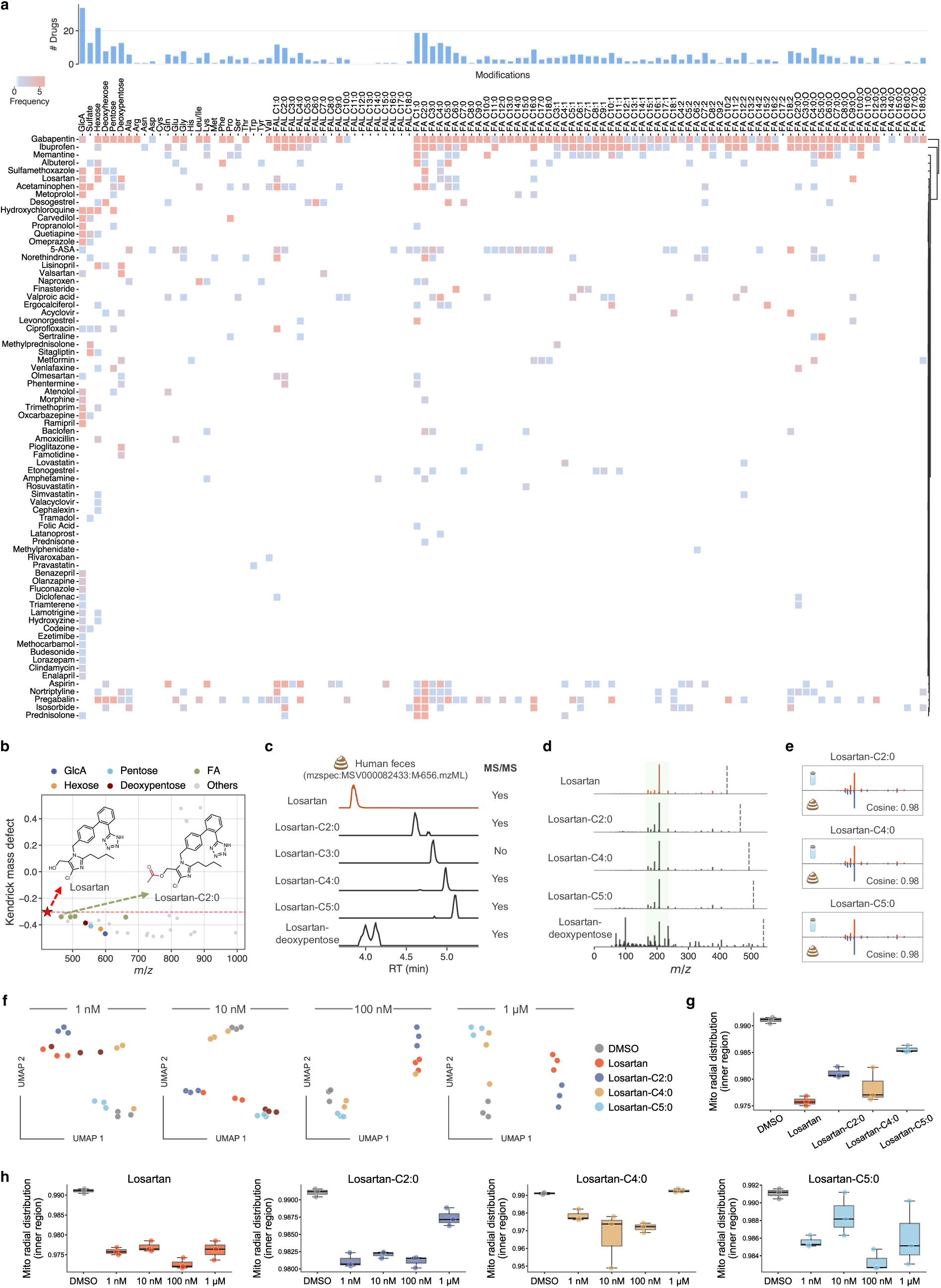
Metabolite conjugate predictions derived from drug molecules. **a**) Predicted drug conjugates that co-occurred with their parent drugs across public metabolomics datasets from the reverse cosine analysis. GlcA, glucuronic acid; FAL, fatty alcohol; FA, fatty acid; FA-O, hydroxy fatty acid. **b**) Kendrick mass defect analysis of predicted losartan conjugates. **c**) Extracted ion chromatograms of losartan and losartan conjugates in a human fecal sample from a public data set (MSV000082433). **d**) Aligned MS/MS spectra of losartan and its conjugates from this fecal sample. **e**) MS/MS matching between synthesized losartan derivatives and human feces, confirming losartan-C2:0, losartan-C4:0 and losartan-C5:0. **f**) UMAP embeddings of morphological cell painting profiles at 1 nM, 10 nM, 100 nM and 1 μM show that losartan acylates cluster distinctly from the parent drug and from DMSO controls. **g**) Mitochondrial radial distributions of DMSO, losartan and losartan acylates at 1 nM. Losartan acylates induce significant redistribution of mitochondrial signal toward the nuclear interior (inner radial shell). **h**) Dose-response effect on mitochondrial radial distributions of losartan and its acylates.

We highlight losartan, ibuprofen and naproxen as representative examples. Losartan is an angiotensin II receptor blocker widely prescribed for hypertension and cardiovascular disease. Beyond its established hepatic oxidation and glucuronidation pathways^48^, our data suggested the existence of additional losartan metabolites, substantially expanding its known metabolic complexity. Kendrick mass defect analysis revealed clusters of predicted losartan conjugates, including both acyl- and sugar-modified derivatives (**Fig. 5b**). In a representative human fecal sample, extracted ion chromatograms showed sequential peaks corresponding to losartan and its conjugated forms (**Fig. 5c**), and MS/MS spectral alignment confirmed their shared structural origin (**Fig. 5d**). To validate these predictions, we synthesized three losartan acylates, losartan-C2:0 (acetate), losartan-C4:0 (butyrate), and losartan-C5:0 (valerate), and confirmed their presence in human samples by MS/MS spectral matching, achieving level 2 annotation confidence^44^ (**Fig. 5e**). Repository-wide searches revealed that these losartan acylates were detected exclusively in human fecal samples, consistent with gut-associated metabolism.

We next examined losartan acylates to determine whether conjugation alters cellular responses. Losartan and three synthetic acylates were profiled using Cell Painting^49^, an image-based phenotypic cell profiling assay, across concentrations from 1 nM to 1 µM. The acylated derivatives produced distinct cellular signatures relative to the parent compound (**Fig. 5f**). Deeper analysis revealed altered mitochondrial radial distributions for losartan acylates relative to losartan, with reduced peripheral and increased perinuclear mitochondrial signal, while remaining intermediate between losartan and DMSO controls (**Fig. 5g**). Whereas parent losartan induced modest changes, losartan acylates produced stronger but acyl group-specific effects. In dose resolved analyses (**Fig. 5h**), losartan-C2:0 and losartan-C4:0 displayed biphasic behavior on mitochondrial radial distribution, with decreased perinuclear mitochondrial enrichment at low concentrations followed by increased perinuclear localization at 1 µM, consistent with stress-associated mitochondrial clustering observed at higher perturbation levels^50^. These results demonstrate that losartan acylates elicit cellular responses distinct from those of the parent drug, indicating that conjugation can meaningfully alter bioactivity.

We next extended this analysis to additional widely used therapeutic drugs by examining two widely used nonsteroidal anti-inflammatory drugs (NSAIDs), ibuprofen and naproxen. Reverse spectral searches consistently identified ethanolamine and creatinine conjugates of both compounds. Although ethanolamine-linked derivatives of ibuprofen and naproxen have been described previously as synthetic prodrug-like conjugates^51,52^, their occurrence as human derived co-metabolites has not been reported. Structural assignments were confirmed using synthesized reference standards by MS/MS spectral matching and retention time agreement with biological samples (**Fig. 6a,b**). Notably, these conjugates were observed in both human urine and serum samples.

**Fig. 6.**
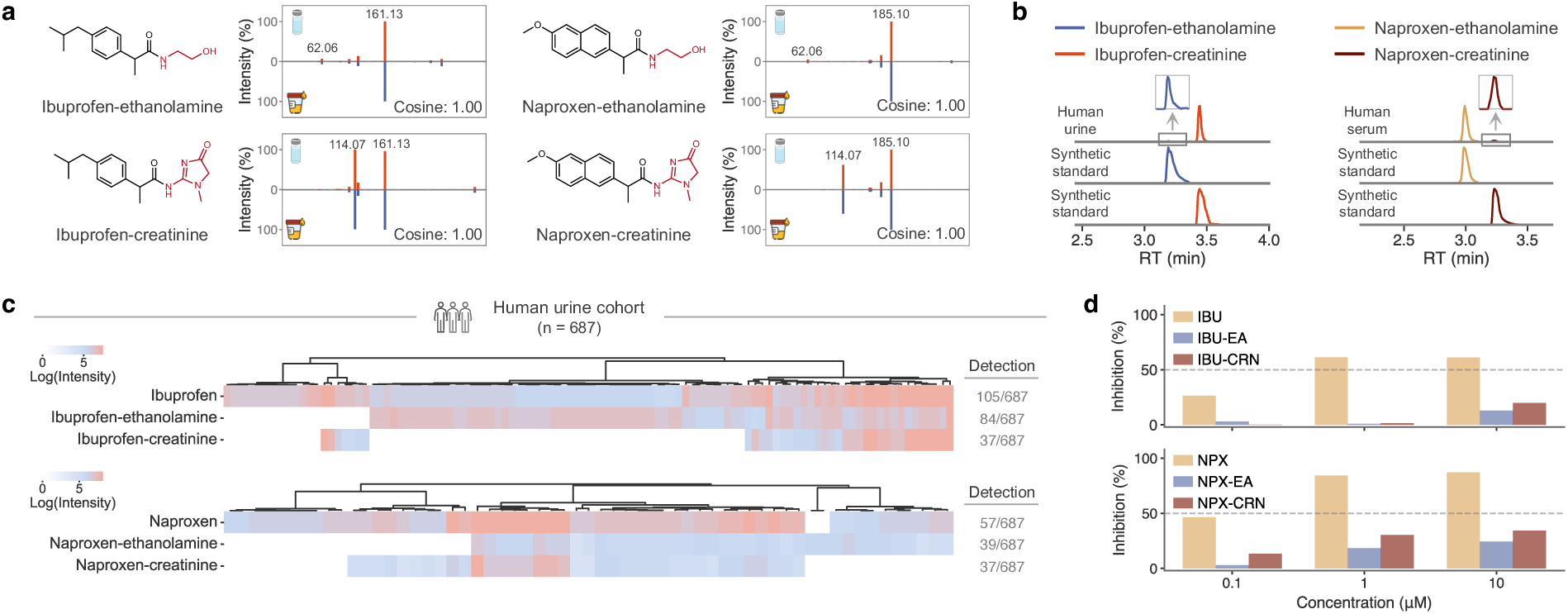
Newly discovered ethanolamine and creatinine conjugations of NSAIDs. **a**) MS/MS spectral matching of ibuprofen-ethanolamine, ibuprofen-creatinine, naproxen-ethanolamine and naproxen-creatinine between synthetic standards and biological samples. **b**) Retention time matching of synthetic drug conjugates against human urine samples. **c**) Detection of ibuprofen, naproxen and their ethanolamine and creatinine conjugates in a human urine cohort (MSV000096359, n = 687). **d**) *In vitro* COX-2 inhibition assay comparing parent drugs and ethanolamine and creatinine conjugates at 0.1 µM, 1 µM and 10 µM. IBU, ibuprofen; EA, ethanolamine; CRN, creatinine; NPX, naproxen.

The prevalence of these conjugates was then assessed in a large human urine cohort comprising 687 individuals (**Fig. 6c**). Ibuprofen-ethanolamine and ibuprofen-creatinine were detected in 84 and 37 individuals, respectively, while naproxen-ethanolamine and naproxen-creatinine were detected in 39 and 37 individuals. For all four drug metabolites, conjugate detection was strongly associated with detection of the corresponding parent drug at the individual level (Fisher’s exact test, *P* < 1.3 × 10^−33^ for all comparisons), whereas conjugates were rarely observed in samples lacking detectable parent drug. These patterns support *in vivo* formation of ethanolamine and creatinine conjugates following drug exposure. To assess whether conjugation alters target engagement, we evaluated the effects of these drug conjugates on cyclooxygenase-2 (COX-2) inhibition using an *in vitro* enzyme assay. Compared to their parent drugs, both ethanolamine and creatinine conjugates showed substantially reduced COX-2 inhibition across the tested concentration range (0.1, 1 and 10 µM) (**Fig. 6d**). At 1 µM, ibuprofen and naproxen inhibited COX-2 by 61.3% and 84.3%, respectively, whereas their ethanolamine conjugates achieved only 0.9% and 18.4% inhibition, indicating attenuated target engagement.

Together, these results demonstrate that ethanolamine and creatinine conjugation of NSAIDs occurs in humans, co-occurs with parent drug exposure, and substantially alters pharmacological activity. More broadly, they provide proof of principle that drug conjugation can generate metabolites with distinct biochemical and cellular properties, underscoring the need to systematically characterize conjugated drug metabolites beyond canonical metabolic pathways.

## Discussion

This work charts the conjugated metabolome across public LC-MS/MS repositories, revealing how covalent fusion between metabolites expands biochemical space. By combining reverse spectral matching with delta mass inference, we annotated millions of supported conjugates encompassing detoxification intermediates, host–microbe co-metabolites, diet-derived molecules, and drug-related products. These results position conjugation as a general mechanism through which living systems generate molecular diversity, link their metabolic networks and may influence biological/biochemical outcomes as shown with representative examples.

Metabolite conjugation has traditionally been examined within narrow biochemical contexts such as bile acid amidation, glutathione conjugation, or glucuronidation. The repository-scale view presented here shows that these reactions are not isolated but part of a broad network of molecular linkages extending across biological kingdoms. The recurrent detection of amino acid conjugates formed with lactic, phenylacetic, and phenylpropionic acids, together with steroid-phosphoethanolamine and drug-derived conjugates, illustrates how this chemistry connects endogenous, microbial, and xenobiotic metabolism. These processes are shaped by diet, microbiome activity, drug metabolism, physiology and disease^53–55^. For instance, phenylacetic acid-amino acid conjugates recur across mammals in a diet-dependent manner, reflecting the coupling of microbial aromatic metabolism with host conjugation pathways.

From a computational perspective, reverse spectral search provides a transparent and scalable framework for substructure-level annotation. It performs robustly across heterogeneous datasets and can be applied directly to the expanding landscape of public reference libraries without retraining or parameter tuning. Emerging artificial intelligence-based methods, such as DreaMS^56^, MIST^57^ and MS2DeepScore^58^, capture higher-order spectral relationships and, as they become more interpretable and transferable, are expected to offer powerful complements to reverse spectral search. In parallel, DeepMet^59^, a large language model trained on known metabolite structures, has been shown to anticipate plausible mammalian metabolites beyond existing databases, enabling generative exploration of chemical space and prioritization of candidate structures for experimental follow-up. Looking ahead, reverse spectral search could ground such generative hypotheses in repository-scale MS/MS evidence by highlighting conjugated metabolites that recur across diverse biological samples and independent studies. Together, these approaches point toward a convergent future in which interpretable algorithms and learning-based models jointly advance large-scale metabolite discovery.

Proper use of this resource starts with recognizing that it provides structure hypotheses rather than confirmed annotations. Reverse spectral matching gives directional evidence, which can be strengthened by considering extra chemical and biological expectations. Targeted prefilters help distinguish plausible conjugates from spectra with unrelated mass differences or artifacts. MassQL^17^ offers a direct route to query diagnostic fragmentation signatures for specific moieties, as we demonstrated for phosphoethanolamine conjugates. Similar logic can be applied to drug-related features. Co-occurrence of a parent drug and its predicted conjugates within the same LC-MS run provides orthogonal support for metabolic linkage and helps exclude spurious matches. Additional filtering based on precursor mass windows, known reactivity of functional groups, expected biosynthetic context or characteristic adduct patterns can further refine predictions. Together, these simple rules reduce false positives and enable users to convert large-scale hypotheses into focused candidate sets for experimental validation.

### Limitations

Several limitations should be acknowledged. The accuracy of substructure inference depends on fragmentation behavior and on the coverage and quality of reference spectra. False discovery rates vary among compound classes, isomeric conjugates or in-droplet reactions^28^ cannot yet be distinguished without orthogonal information such as chromatographic separation, ion mobility, deep expert knowledge about fragmentation information^60^, X-ray, or NMR, limiting structural resolution in cases where multiple linkage configurations or closely related isomers are possible. For example, a reverse cosine match to cis-3-hexenoic acid should be interpreted as a C6:1 fatty acid moiety, with the full set of plausible C6:1 isomers and reaction chemistries considered during follow-up.

Certain mass spectrometry phenomena can complicate the interpretation of conjugate-like MS/MS relationships. Some true conjugates may remain unannotated due to ionization-related effects, such as adduct formation, which are not yet explicitly modeled in the current framework and may therefore be missed. This limitation highlights an opportunity to expand the conjugation metabolome by systematically accounting for multiple adduct forms. In addition, although no such cases were observed in this study, in-source processes that alter precursor ion populations could also contribute to misannotation^28^. Finally, noncovalent associations, including heteromer formation or co-isolation and co-fragmentation of co-eluting species, can generate chimeric or composite spectra and could potentially result in mass relationships that mimic covalent conjugation^61^. Together, these considerations underscore the need for continued development of computational filtering strategies, contextual validation, and, where appropriate, manual curation supported by retention time matching to synthetic standards, as demonstrated here.

The reverse cosine approach relies on the availability of reference spectra; not all potential conjugation building blocks are represented, so expanded libraries will enable discovery of additional conjugates, including sequential or higher-order conjugations. Currently, the framework resolves two-component conjugates, whereas metabolites may undergo multi-step modifications in biological systems yielding more complex fragmentation patterns. Finally, although plausible conjugate pairs are identified, the precise site of conjugation is not determined - for instance, phenylpropionic acid-lysine can form amide linkages at either amino group, and both positional isomers were observed in microbial cultures (**Extended Data Fig. 10**).

The conjugate annotations generated here extend the reach of existing reference libraries and molecular networking-based annotation workflows. They can suggest structural hypotheses when no direct spectral match exists and can recover related chemistries that fail to propagate through molecular networks. But predictions require access to the synthetic standards and samples for validation. The approach also provides a practical bridge between data-centric discovery and synthesis-driven metabolomics. The predictions are being integrated into the ongoing multiplexed synthetic pipeline as part of our lab’s effort to annotate every detectable metabolite for future experimental validation. Indeed in a parallel large-scale synthesis effort, which was initiated prior to the reverse cosine based strategy, created over 170,000 compounds, we observed 1,061,694 MS/MS matches among the 24 million predicted spectra (cosine ≥ 0.8, level 2 annotation^44^), when compared to the reference library generated in our large-scale synthesis study^16^ (top 1,000 MS/MS matches are shown as **Supplementary File 1**) and many of the remaining predictions we found here will become part of future synthesis efforts.

Despite these caveats, this study demonstrates that existing public metabolomics data contain extensive, previously underutilized information on biochemical relationships that can be computationally reconstructed into testable hypotheses. By integrating large-scale data mining with chemical synthesis and biological validation, this work provides both a resource and a framework for systematically exploring conjugation chemistry and expanding the known chemical space of living systems.

## Methods

In addition to the molecule discovery workflows presented in this study, which show how the reverse cosine framework can be used to generate and test structure hypotheses across public metabolomics datasets, we also provide three complementary resources derived from the reverse cosine results of metabolite conjugations. The complete dataset, containing search results from 149.9 million clustered MS/MS spectra, is available on Zenodo (https://zenodo.org/records/17245769) for open exploration. A curated and refined subset is released as a reference MS/MS library within the GNPS propagated libraries (https://external.gnps2.org/gnpslibrary, “GNPS-CONJUGATED-METABOLOME”, 4 partitions in total), and can be used in routine library searching to supply structure hypotheses. An interactive web application is also available (https://conjugated-metabolome.gnps2.org), enabling users to query predicted conjugates for molecules of interest using compound names or SMILES strings.

### Reverse spectral search algorithm

Reverse spectral search is a template-based MS/MS similarity framework originally proposed for spectral identification and later extended to improve robustness to chimeric spectra^12,13^. Its asymmetric scoring design makes it particularly well suited for substructure annotation, where only a portion of the query molecule is expected to match a known structure. By treating the reference spectrum as a template and scoring only fragment ions characteristic of that structure, reverse spectral search captures partial structural overlap without penalization from additional fragments present in the query spectrum.

Formally, a reference MS/MS spectrum and a query MS/MS spectrum are compared after data centroiding, intensity scaling and normalization. Fragment ions in the reference spectrum are aligned to fragment ions in the query spectrum within a specified mass tolerance, optionally allowing a precursor mass offset to account for structural modifications. Only fragment ions present in the reference spectrum contribute to the similarity score. Reference fragment ions that cannot be aligned reduce the similarity score, whereas fragment ions present only in the query spectrum do not directly penalize the score. This scoring scheme suppresses interference from unrelated fragments and preserves sensitivity to shared substructures, enabling reliable substructure annotation even when the query spectrum corresponds to a chemically modified or conjugated molecule.

In this study, we used the modified (also referred to as hybrid search^62^) reverse cosine similarity, in which fragment ion alignment is performed after accounting for a potential precursor mass offset between query and reference spectra. This allows fragment ions derived from a shared substructure to be aligned even when the query precursor mass differs due to covalent modification. In addition to the similarity score and the number of matched fragment ions, spectral usage was used as an orthogonal quality metric and is defined as the fraction of the total reference fragment ion intensity explained by matched peaks in the query spectrum.

### Preparation of MS/MS reference libraries for reverse spectral search

A comprehensive MS/MS reference library was established by integrating four major spectral libraries: GNPS MS/MS library (downloaded on Nov 11, 2024), MoNA (MassBank of North America, downloaded on Oct 17, 2024), MassBank (2024.06 release) and NIST20 (commercially available). The database was refined by eliminating entries lacking valid SMILES or InChIKey identifiers. Reference spectra with the following precursor types were retained: M+H, M+H–H_2_O, M+NH_4_ and M–H. Reference MS/MS collected using low-resolution MS instruments (e.g., ion trap, QQQ) were excluded. Using RDKit, compounds were filtered to retain only those containing at least one of the following functional groups: carboxyl, hydroxyl, primary amine or secondary amine. Structure conversion (SMILES and InChIKey) and compound classification (NPClassifier^18^ and ClassyFire^63^) were completed via the GNPS API. The final curated library comprised 818,648 MS/MS reference spectra, representing 43,017 unique chemical compounds.

### Pan-repository MS/MS data retrieval and filtering

In this study, more than 1.32 billion MS/MS spectra from 3,340 datasets across GNPS/MassIVE, MetaboLights and Metabolomics Workbench were retrieved, filtered and clustered. Proteomics or data-independent acquisition (DIA) datasets were excluded through keyword searches in dataset titles or dataset descriptions (e.g., “protein”, “proteome”, “proteomics”, “data independent”, “DIA”). Metabolomics datasets with valid mzML or mzXML files were processed, and profile MS data files were discarded. The following analysis was restricted to MS files with known ion polarities and MS instrument types, specifically those collected using Orbitrap, TOF, or FT-ICR instruments, while excluding data from low-resolution MS instruments, GC-MS, or data-independent acquisition mode. At the MS/MS spectrum level, we applied the following criteria: (1) precursor *m/z* is in the range of 50–2000, (2) the spectrum contains at least 5 fragment peaks with >1% relative intensities, (3) the precursor is singly charged, and (4) top 50 peaks were retained. Above filterings led to a total of 539,204,333 curated high-resolution MS/MS spectra, including 429,964,017 positively ionized spectra and 109,240,316 negatively ionized spectra.

### Pan-repository MS/MS clustering

Falcon^15^ (version 0.1.3) was used to cluster pan-repository MS/MS spectra using the following parameters: precursor mass tolerance, 20 ppm; fragment peak mass tolerance, 0.05 Da; cosine distance for clustering, 0.05; minimum peaks, 4; minimum mass range, 0; minimum peak *m/z*, 0; maximum peak *m/z*, 2000. Other parameters were kept at default values. Both singleton spectra and cluster representative spectra were exported. To prevent excessive memory usage, MS/MS spectra were split into distinct MGF files based on their ion polarities and precursor masses prior to clustering, and MS/MS clustering was performed individually for each dataset. This yielded 118,764,042 positively ionized MS/MS clusters and 31,181,284 negatively ionized MS/MS clusters.

### Pan-repository reverse spectral search of clustered MS/MS spectra

Reverse spectral search was applied at scale to 149.9 million clustered MS/MS spectra derived from public metabolomics repositories and searched against a curated reference library comprising 818,648 MS/MS spectra. This search space corresponds to approximately 122.8 trillion potential spectrum-spectrum comparisons. To enable feasible computation at this scale, we implemented and adapted the flash entropy framework^62^ for efficient reverse cosine computation. All analyses were performed on a Linux workstation running Ubuntu 20.04.6 LTS with a 24-core Intel Xeon Gold 5220R processor (2 threads per core) and 512 GB of RAM. The complete pan-repository reverse spectral search was completed within approximately one month of wall-clock time.

For the primary reverse spectral search, matches were retained if they met the following criteria: reverse cosine score ≥ 0.8, at least three matched fragment ions, and spectral usage ≥ 30%. To support inference of conjugation events in cases where only one substructure could be confidently matched, a secondary reverse spectral search was performed using relaxed thresholds (reverse cosine score ≥ 0.7, at least three matched fragment ions, and spectral usage ≥ 15%). In these cases, the precursor mass difference between the query spectrum and the matched reference spectrum was interpreted as a candidate conjugating unit, subject to downstream chemical and biological filtering. In total, 22,431,333 positive-mode and 1,796,106 negative-mode MS/MS spectra were putatively annotated with at least one inferred substructure component.

### Construction of the conjugated metabolome MS/MS library

We constructed a curated reference MSMS library from a filtered subset of the reverse cosine predictions. The aim was to retain high confidence conjugate spectra that can serve as reference entries while excluding spectra with uncertain assignments or weak structural support. We first applied class specific FDR thresholds derived from the multiplex synthesis benchmark. Classes that reached a 1% FDR were filtered using the corresponding class level score and matched peak requirements. For compound classes that could not meet this threshold, a uniform stringent cutoff was applied: minimum reverse cosine score 0.9, at least 6 matched peaks, and at least 80% spectral usage. These rules removed low confidence predictions and restricted the library to spectra with strong evidence for the inferred substructures. To avoid introducing entries that corresponded to known compounds already represented in public libraries, we removed any predicted conjugate spectrum that matched to reference libraries at cosine 0.7 or higher with at least 6 shared peaks. For each unique inferred substructure pairing, we kept only the spectrum with the highest reverse cosine score. This step held the size of the library at a manageable scale. After filtering, 165,379 positive mode spectra and 32,688 negative mode spectra remained.

Delta mass values associated with the conjugating units were then annotated using *msbuddy* (https://github.com/Philipbear/msbuddy, version 0.3.12) to provide possible molecular formula assignments. Each curated spectrum was stored with its USI to maintain traceability to the underlying public dataset. The resulting collection was compiled into “GNPS-CONJUGATED-METABOLOME” reference libraries for routine MS/MS annotation workflows.

### Pan-repository analysis of predicted metabolite conjugates

To enable retrospective analysis of predicted conjugates across the public untargeted metabolomics data, we retained the Universal Spectrum Identifiers (USIs) for all inferred MS/MS spectra. These USIs were mapped to the harmonized metadata from Pan-ReDU^19^, which standardizes sample type, organism, and body site information across GNPS, MetaboLights, and Metabolomics Workbench. The NCBITaxonomy column was filtered for human data files (“9606|Homo sapiens”) and rodent data files (“10088|Mus”, “10090|Mus musculus”, “10105|Mus minutoides”, “10114|Rattus” and “10116|Rattus norvegicus”). The UBERONBodyPartName field was used to assign spectra to body sites. Files originating from foods, plants, or microbial cultures were labeled using the corresponding domain-specific MASST resources (foodMASST^33^, plantMASST^34^ and microbeMASST^32^). These tables provide controlled vocabulary labels. Only files with valid Pan-ReDU metadata were retained. Each USI was joined to its file level metadata to create a table linking every predicted conjugate to its sample source.

### Evaluation of reverse cosine search on substructure annotation

To evaluate the performance of reverse cosine scoring on substructure annotation, we applied it on a reference MS/MS library created via multiplex synthesis from a diverse range of compounds, including conjugates of endogenous compounds such as bile acids, neurotransmitters and vitamins, and conjugates of xenobiotic compounds. The library can be downloaded on GNPS2 (https://external.gnps2.org/gnpslibrary, “MULTIPLEX-SYNTHESIS-LIBRARY-FILTERED”, 4 partitions in total). We further filtered out the library by selecting the protonated reference spectra for chemical conjugates without ambiguous isomeric structures (0 isomers and 0 isobar peaks), resulting in a total of 9,156 spectra for 7,575 unique chemical conjugates.

To establish ground-truth substructures, two complementary structure similarity metrics were utilized, the myopic Maximum Common Edge Subgraph (mMCES) distance^64^ and Maximum Common Substructure (MCS) analysis. Both metrics were implemented in Python using the *myopic_mces* and *rdkit* packages. We here illustrate the procedure using deoxycholyl-phenylalanine as an example. First, SMILES strings were collected for the conjugate and its two components (deoxycholic acid and phenylalanine). The conjugate MS/MS spectrum was then queried against the reference library using reverse cosine scoring, and candidate matches, along with their corresponding SMILES strings, were retrieved (for instance, chenodeoxycholic acid). We next calculated the mMCES distances between the matched structure (chenodeoxycholic acid) and each component (deoxycholic acid and phenylalanine) to assess whether the spectral match was structurally similar to either constituent. Matches with mMCES distance ≤ 4 were considered valid substructures, corresponding to tolerance for up to two bond differences from a component structure. In parallel, we computed the maximum common substructure (MCS) between the matched compound and the conjugate. If the atom count difference between the MCS and the matched compound was ≤ 1, the match was accepted as a valid substructure, accounting for the water loss typically accompanying conjugation.

### Pan-repository MS/MS search of synthesized compounds and phenotype association

Reference MS/MS spectra from the synthesized conjugates were searched against public metabolomics repositories using MASST^29^. Batch submission was performed with *microbe_masst* in Python (https://github.com/robinschmid/microbe_masst). Searches used a minimum cosine score of 0.7, at least 4 matched fragment ions, MS1 tolerance 0.05 Da and MS/MS tolerance 0.05 Da. MASST returns all matching MS/MS scans together with the dataset and file identifiers. These outputs also include domain MASST results that classify files as microbe, plant, or food related when available. For phenotype association, MASST results were merged with the Pan-ReDU metadata table. We treated each LC-MS file as one biological observation. Disease related information was taken from the DOIDCommonName and HealthStatus fields. Body site information was taken from UBERONBodyPartName, and taxonomic information from NCBITaxonomy.

### Analysis of tomatidine conjugates

MS/MS spectra were first extracted from mzML files generated in the microbial co-culturing experiments and compiled into an MGF file. This MGF file was searched against the combined MS/MS reference library comprising 818,648 spectra using reverse cosine scoring. Candidate tomatidine-derived spectra were then retrieved from the reverse cosine results by filtering for matches to the planar InChIKey of tomatidine (XYNPYHXGMWJBLV). In total, 190 MS/MS spectra corresponding to candidate tomatidine conjugates were retained and exported as a separate MGF file. This file was subsequently analyzed using a classical molecular networking workflow on GNPS2 with the following parameters: minimum cosine for library search, 0.7; minimum matched peaks for library search, 6; MS1 mass tolerance, 0.02 Da; MS/MS mass tolerance, 0.02 Da; minimum cosine for networking, 0.7; minimum matched peaks for networking, 6; MS/MS clustering, enabled; minimum cluster size, 1. The classical molecular networking job is available at https://gnps2.org/status?task=06f0b0e04b1e4fe6b05deeb0de73ea7b.

### Analysis of steroid-phosphoethanolamine conjugates

For candidate phosphoethanolamine conjugates, we applied the following MassQL^17^ filter to capture spectra containing either the diagnostic fragment at *m/z* 142.0264 or the neutral loss of 141.0191 Da: *QUERY scaninfo(MS2DATA) WHERE MS2PROD=142.0264:TOLERANCEMZ=0.02:INTENSITYPERCENT=5 OR MS2NL=141.0191:TOLERANCEMZ=0.02:INTENSITYPERCENT=5* From the set of predicted phosphoethanolamine conjugates, we then selected steroid related candidates using ClassyFire classification and keeping only those assigned to the class “Steroids and steroid derivatives”. The resulting list is provided in **Supplementary Table 3**.

### Analysis of drug conjugates

The list of the 200 most prescribed drugs in 2022 was obtained from the “Top 200 posters” compiled by the Njardarson Laboratory^47^, of which 175 were xenobiotic small-molecule drugs. To assess co-occurrence of parent drugs and their predicted conjugates in public LC-MS/MS datasets, we first retrieved reference MS/MS spectra for the 175 drugs from the combined GNPS, MoNA, MassBank and NIST20 libraries. These reference spectra were searched against all public metabolomics repositories using MASST, which returned MS/MS-level matches together with the corresponding dataset and LC-MS file identifiers in which each parent drug was detected. This established the set of LC-MS data files with MS/MS-supported evidence for the presence of each parent drug. In parallel, predicted conjugates were extracted from the reverse cosine search results, with their associated LC-MS data files identified via Universal Spectrum Identifiers (USIs). Co-occurrence was defined at the LC-MS data file level, requiring that each predicted conjugate be detected in a file that also contained MS/MS evidence for the corresponding parent drug. Predicted conjugates lacking parent-drug evidence in the same file were excluded, prioritizing metabolically plausible conjugates while potentially excluding cases of complete or near-complete drug metabolism.

### Chemical synthesis

A total of 55 metabolite conjugates were synthesized and validated in this study, including lactic acid-amino acid conjugates, phenylacetic acid-amino acid conjugates, phenylpropionic acid-amino acid conjugates, steroid-phosphoethanolamine conjugates, and conjugates of losartan, ibuprofen and naproxen.

#### Synthesis of lactic acid-amino acid conjugates

Methyl lactate (1.0 eq.) and a pooled amino acid mixture (each at 0.01 eq.; 20 proteinogenic amino acids) were dissolved in methanol (20 mL) and refluxed overnight. After cooling to room temperature, the reaction mixture was concentrated *in vacuo*, and the residue was directly reconstituted for LC-MS/MS analysis.

#### Synthesis of phenylacetic acid- and phenylpropionic acid-amino acid conjugates

Phenyacetic acid or phenylpropionic acid (1.0 eq.) was dissolved in DMF (2 mL) in a 20 mL scintillation vial equipped with a magnetic stir bar. Solid HATU (1.2 eq.) and neat DIPEA (1.5 eq.) were added sequentially, and the mixture was stirred at 23 °C for 15 min. Amino acids (each at 0.01 eq.; 20 proteinogenic amino acids) were then added, and the reaction was stirred overnight. The mixture was concentrated *in vacuo*, and the residue was reconstituted for LC-MS/MS analysis.

#### Synthesis of steroid-phosphoethanolamine conjugates

Boc-glycerinol (1.0 eq.) and tris(2,2,2-trifluoroethyl) phosphate (1.02 eq.) were dissolved in toluene (3 mL) under N_2_ and then cooled to 0 °C. DBU (1,8-Diazabicyclo[5.4.0]undec-7-ene, 1.0 eq.) was then added. The reaction mixture was stirred for 5 h at room temperature and quenched at 0 °C by addition of phosphate buffer (pH 7). The mixture was extracted twice with ethyl acetate. The combined organic layers were washed with aqueous brine, dried over Na_2_SO_4_, and concentrated *in vacuo*. The residue was purified by CombiFlash normal-phase chromatography (80 g column, 40 mL/min) using dichloromethane (solvent A) and methanol (solvent B) with the following gradient: 0–5 min, 5% B; 5–10 min, 10% B; 10–15 min, 30% B; 15–20 min, 80% B. tert-Butyl (2-((bis(2,2,2-trifluoroethoxy)phosphoryl)oxy)ethyl)carbamate eluted at 15 min and was obtained in 95% yield. A steroid (1.0 eq.) was dissolved in toluene (4 mL) and cooled to 0 °C. Lithium tert-butoxide (2.5 eq., 1.0 M in hexane) was added, and the mixture was stirred for 1 h at 0 °C. A toluene solution of tert-butyl (2-((bis(2,2,2-trifluoroethoxy)phosphoryl)oxy)ethyl)carbamate (1.5 eq.) was then added at −45 °C. The reaction was stirred for 3 h and quenched at −45 °C by addition of acetic acid (4.1 eq.) in toluene, followed by phosphate buffer (pH 7). The mixture was extracted twice with ethyl acetate. The combined organic layers were washed with aqueous brine, dried over Na_2_SO_4_, and concentrated *in vacuo*. The residue was used directly in the next step without further purification. tert-Butyl (2-((bis(2,2,2-trifluoroethoxy)phosphoryl)oxy)ethyl)carbamate-steroid (1.0 eq.) was dissolved in toluene (5 mL) and cooled to −45 °C. Lithium tert-butoxide (14.5 eq., 1.0 M in hexane) was added, and the reaction mixture was stirred for 3 h at −45 °C. The reaction was quenched at −45 °C by addition of acetic acid (1.2 eq.) in toluene, followed by phosphate buffer (pH 7) at 0 °C. The mixture was extracted twice with ethyl acetate. The combined organic layers were washed with aqueous brine, dried over Na_2_SO_4_, concentrated *in vacuo*, and used directly in the next step. tert-Butyl (2-(((tert-butylperoxy)(2,2,2-trifluoroethoxy)phosphoryl)oxy)ethyl)carbamate-steroid (1.0 eq.) was dissolved in dichloromethane (5 mL) and cooled to 0 °C. Hydrogen chloride in 1,4-dioxane (4 M, 6.0 eq.) was added, and the reaction mixture was stirred for 3 h at room temperature. The mixture was concentrated *in vacuo* to afford steroid-phosphoethanolamine, which was used directly for LC-MS/MS analysis.

#### Synthesis of losartan acylates

Losartan (1.0 eq.) and anhydrous DCM (2.5 mL) were added to a 20 mL scintillation vial equipped with a magnetic stir bar. The solution was cooled to 0 °C using an ice bath. Triethylamine (1.5 eq.) was then added. While stirring at 0 °C, a mixture containing each acyl chloride (0.01 eq.) was added dropwise. The reaction mixture was stirred for 3 h and allowed to gradually warm to 23 °C. The mixture was then concentrated *in vacuo*, and an LC-MS/MS sample was prepared.

#### Synthesis of ibuprofen-ethanolamine, ibuprofen-creatinine, naproxen-ethanolamine and naproxen-creatinine

Ibuprofen or naproxen (1.0 eq.) was dissolved in anhydrous DMF (2.5 mL) in a 20 mL scintillation vial equipped with a magnetic stir bar. HATU (1.2 eq.) and triethylamine (1.2 eq.) were added sequentially, and the mixture was stirred for 20 min at room temperature. Ethanolamine or creatinine (1.0 eq.) was then added, and the reaction was stirred overnight at room temperature. The reaction mixture was concentrated *in vacuo*, and the residue was reconstituted for LC-MS/MS analysis.

### LC-MS/MS data acquisition for synthetic standards and biological samples

LC-MS/MS analysis was performed using a Vanquish UHPLC system coupled to a Q Exactive Orbitrap mass spectrometer (Thermo Fisher Scientific). Chromatographic separation was achieved on a reverse-phase polar C18 column (Kinetex Polar C18, 100 mm × 2.1 mm, 2.6 µm particle size, 100 Å pore size). The column compartment was maintained at 40 °C using forced-air temperature control, and the autosampler was held at 10 °C. Injection volume was set to 5 µL. The mobile phase consisted of solvent A (water with 0.1% formic acid) and solvent B (acetonitrile with 0.1% formic acid). The flow rate was set to 0.50 mL/min. The gradient was as follows: 0 min, 5% B; 1 min, 5% B; 2 min, 40% B; 3 min, 98% B; 5 min, 98% B; 5.01 min, 5% B; 6 min, 5% B. Mass spectrometry data were acquired in positive ion mode using data-dependent acquisition. Full MS scans were acquired at a resolution of 35,000 over an *m/z* range of 100–1000, with an AGC target of 1 × 10^6^ and a maximum injection time of 100 ms. MS/MS scans were acquired at a resolution of 17,500 with an AGC target of 2 × 10^5^, a maximum injection time of 150 ms, an isolation window of 1.0 *m/z*, and TopN set to 5. Stepped normalized collision energies of 25%, 40%, and 60% were applied. Dynamic exclusion was set to 1.0 s, isotope exclusion was enabled, and lock mass correction was used during acquisition.

### LC-MS/MS data analysis

Raw LC-MS/MS data (Thermo .raw files) were converted to mzML files using ProteoWizard MSConvert (version 3.0). For targeted quantitative analysis, we manually created a list of target *m/z* values with associated retention times, then used a custom Python script to extract metabolite peak heights from the mzML files. For **Fig. 3h**, **Fig. 3i** and **Fig. 4d**, sample metadata were obtained from Pan-ReDU and merged with the peak height table for downstream visualization. For **Fig. 6c**, the limit of detection was set to 1 × 10^6^; features were considered detected only if their peak heights exceeded this threshold.

### Microbial culturing, extraction and LC-MS/MS analysis

Microbial culture data were generated as part of the microbiomeMASST effort in our lab^36^, and were used here to discover and validate the tomatidine-phosphoethanolamine conjugate. Bacterial community composition used in this study can be found in **Supplementary Table 2**. Cultures were started from glycerol stock preserved at −80 °C and were incubated in an anaerobic chamber (20% CO_2_, 5% H_2_, and 75% N_2_) at 37 °C in modified brain heart infusion (BHI) medium, with pH adjusted to 7.5 using 5 N NaOH. Modified BHI consisted of brain heart infusion broth (35 g/L) supplemented with yeast extract (10 g/L), proteose peptone (5 g/L), cellobiose (2 g/L), maltose (1 g/L), and sodium pyruvate (2 g/L). The medium was further supplemented with vitamin K/hemin solution (final concentrations 0.01 mg/L vitamin K and 0.1 mg/L hemin), a short-chain fatty acid mixture (3.1 mL/L; acetic, propionic, butyric, isovaleric, isobutyric, valeric, and 2-methylbutyric acids), trace mineral supplement (10 mL/L), vitamin supplement (10 mL/L), and L-cysteine HCl as a reducing agent. Each bacterial culture was normalized to an optical density (OD) at 600 nm of 0.02 prior to mixing at an equal proportion to create the defined groups. Communities were grown in 96-well plates (edge wells filled with sterile medium to minimize evaporation) for 72 h in the presence or absence of tomatidine (final concentration 10 µM; DMSO stock at 1 mM). After incubation, cultures were extracted with pre-chilled 80% MeOH/H_2_O at a 1:4 (v/v) sample-to-solvent ratio and incubated at 4 °C overnight. Samples were centrifuged at 2,500 rpm for 10 min, supernatants transferred to 96-well plates, dried overnight in a CentriVap, and stored at −80 °C until analysis.

Extracts were analyzed by LC-MS/MS using a Vanquish UHPLC system coupled to a Orbitrap Exploris 240 mass spectrometer (Thermo Fisher Scientific). Chromatographic separation was achieved on a reverse-phase polar C18 column (Kinetex Polar C18, 100 mm × 2.1 mm, 2.6 µm particle size, 100 Å pore size). The column compartment was maintained at 40 °C using still-air temperature control, and the autosampler was held at 4 °C. Injection volume was set to 5 µL. The mobile phase consisted of solvent A (H_2_O with 0.1% formic acid) and solvent B (ACN with 0.1% formic acid). The flow rate was set to 0.5 mL/min. The gradient was as follows: 0 min, 5% B; 1 min, 5% B; 7.5 min, 40% B; 8.5 min, 99% B; 9.5 min, 99% B; 10 min, 5% B; 10.5 min, 5% B; 10.75 min, 99% B; 11.25 min, 99% B; 11.5 min, 5% B; 12 min, 5% B. Mass spectrometry data were acquired in positive ion mode using data-dependent acquisition. Full MS scans were acquired at a resolution of 60,000 over an *m/z* range of 100–1000, RF Lens (%) set to 70, with an AGC target to Standard and a maximum injection time set to auto. MS/MS scans were acquired at a resolution of 22,500 with an AGC target set to “Custom” with a normalized AGC Target of 200%, a maximum injection time set to “auto”, an isolation window of 1.0 *m/z*, and TopN set to 10. Stepped normalized collision energies of 25%, 40%, and 60% were applied. Dynamic exclusion was set to “custom”, excluded after 2 times if it occurs within 3 s for a duration of 6 s, and isotope exclusion was enabled.

### Cell painting

U2-OS cells (ATCC, HTB-96) were cultured in DMEM (Gibco, 11965092) supplemented with 5% fetal bovine serum (FBS; Gibco, A5256801), 100 units/mL penicillin, and 100 µg/mL streptomycin at 37 °C in a humidified incubator (ambient O_2_, 5% CO_2_). For profiling, cells were seeded on Day 0 in 384-well plates at 2,000 cells per well in 50 µL DMEM supplemented with 2% FBS, 100 units/mL penicillin, and 100 µg/mL streptomycin, and incubated overnight. On Day 1, cells were treated with losartan or losartan acylates (losartan-C2:0, losartan-C4:0, losartan-C5:0) at 1 nM, 10 nM, 100 nM, and 1 µM final concentration for 24 h at 37 °C (ambient O_2_, 5% CO_2_). Compounds were prepared in DMSO; DMSO-only wells served as vehicle controls. To mitigate edge effects, perimeter wells were filled with DMEM without cells and excluded from downstream imaging analysis. Plates were placed in a humidified container (wet paper towels) during incubation.

#### Staining and imaging

Cell Painting staining was performed using the Invitrogen image-iT Cell Painting Kit (I65000) following the manufacturer’s user guide. Stock solutions of the six fluorescent dyes were prepared as recommended by the manufacturer by dissolving dyes in DMSO, sodium bicarbonate, and dH_2_O as required. Cells were stained, washed, and fixed according to the kit protocol^65^. Images were acquired on an ImageXpress Micro XLS wide-field epifluorescence microscope (Molecular Devices) using a 20× objective and exported as TIFF files for analysis.

#### Image processing and feature extraction

Images were processed in CellProfiler^66^ (version 4.2.8). Illumination correction was performed using the CellProfiler project file (Illum.cpproj) to generate multi-image illumination correction functions, which were saved in NumPy array format. Corrected images were then segmented and quantified using the CellProfiler project file (analysis.cpproj) to extract morphological profiling features, including intensity, texture, shape, and contextual measurements. CellProfiler project files were deposited on Zenodo (https://zenodo.org/records/18408023).

#### Data preprocessing, feature filtering, and normalization

Per-image features were merged with treatment metadata by ImageNumber. Wells were restricted to vehicle controls (DMSO) and the assayed compound dose conditions, and profiles with fewer than 300 detected objects were excluded. Non-morphological and identifier features were removed, including object count and parent–child relationship fields, object identifier columns, and nuclei location coordinate features. Site-level features were averaged within each well to generate one profile per well. Features were filtered to remove unstable and uninformative measurements by excluding those with high variance across DMSO wells (above the 95th percentile), low variance across all wells (below the 5th percentile), or extreme dispersion in DMSO as assessed by the MAD/SD ratio (retained range 0.1 to 10). Remaining features were normalized to DMSO using a robust z-score based on the median and median absolute deviation (scaled by 1.4826). The resulting normalized feature matrix was used for downstream UMAP analysis.

#### UMAP analysis

The cleaned, normalized feature matrix was embedded with UMAP (*umap-learn* in Python) using Euclidean distance. UMAP was run separately for each compound concentration (1 nM, 10 nM, 100 nM, and 1 µM), each time using DMSO wells together with wells at the queried concentration. The number of neighbors was set to 3 and min_dist to 0.1. The resulting two-dimensional UMAP embeddings were used to assess compound-level separation.

### COX-2 enzyme inhibition assay

COX-2 inhibition was assessed using the COX Colorimetric Inhibitor Screening Assay Kit (Cayman Chemical, Item No. 701050) according to the manufacturer’s instructions. Test compounds, including ibuprofen, ibuprofen-ethanolamine, ibuprofen-creatinine, naproxen, naproxen-ethanolamine and naproxen-creatinine, were dissolved in dimethyl sulfoxide (DMSO) and assayed at final concentrations of 0.1, 1, and 10 µM. For each condition, measurements were performed in duplicate, and mean absorbance values were used for analysis. Briefly, human recombinant COX-2 enzyme was preincubated with test compounds or vehicle control in an assay buffer containing hemin for 5 min at 25 °C. Reactions were initiated by addition of arachidonic acid substrate and allowed to proceed for exactly 2 min before measurement. COX-2 activity was quantified colorimetrically by monitoring oxidation of the chromogenic substrate (TMPD) at 590 nm using a TECAN 96-well plate reader. Background absorbance from non-enzymatic control wells was subtracted prior to calculation of percent inhibition, which was determined relative to vehicle-treated controls.

### Statistics

No statistical methods were used to predetermine sample size. Two-sided Mann-Whitney *U* tests were used for two-group comparisons of metabolite abundances (**Fig. 3h,i**). Associations between detection of parent drugs and their predicted conjugates across individuals were tested using Fisher’s exact test (**Fig. 6c**); the reported *P* values correspond to two-sided tests. No randomization or blinding was performed. Data processing and statistical analyses were carried out in Python 3.10.

## Data availability

All LC-MS/MS data involved in this study are publicly available on GNPS/MassIVE: mammalian feces (MSV000086131), diabetes study (MSV000082261), tomato seedling extracts (MSV000083306), microbial co-culturing (MSV000098638), human feces for losartan acylates (MSV000082433), human urine cohort for drug conjugates (MSV000096359), retention time matching between synthetic chemical standards and biological samples (MSV000099690).

Raw conjugate search results from 149.9 million clustered MS/MS spectra are available on Zenodo (https://zenodo.org/records/17245769). A rigorously curated subset was compiled into a reference MS/MS library and is accessible on GNPS2 (https://external.gnps2.org/gnpslibrary, “GNPS-CONJUGATED-METABOLOME”, four partitions). An interactive web application for exploring the conjugated metabolome is available at https://conjugated-metabolome.gnps2.org (also hosted at https://conjugated-metabolome.streamlit.app). The classical molecular networking job for tomatidine and its predicted conjugates is available at https://gnps2.org/status?task=06f0b0e04b1e4fe6b05deeb0de73ea7b.

## Code availability

Source codes for the reverse spectral search algorithm and associated data analysis are available on GitHub (https://github.com/Philipbear/conjugated_metabolome) and Zenodo (https://zenodo.org/records/18408023).

## Acknowledgement

This work is supported, in part, by National Institute of Health Sciences R01DK136117, U24DK133658, BBSRC/NSF award 2152526, and Chan Zuckerberg Initiative. VCL is supported by Fonds de recherche du Québec - Santé (FRQS) Postdoctoral fellowship (335368) and from Natural Sciences and Engineering Research Council of Canada (NSERC) postdoctoral fellowship (598938). LAB is supported by Reproductive Scientist Development Program Grant. HNZ is supported by the National Institute Of Environmental Health Sciences of the National Institutes of Health under Award Number K99ES037746. YEA acknowledges funding of ÖAW through APART-USA.

## Author contributions

Conceptualization, SX and PCD; Methodology, SX; LC-MS/MS data collection, AP, JA, VCL, HG and SX; Data analysis, SX; Chemical synthesis, AP, JA and ZH; Microbial culturing, VCL and HG; Cell painting, HG and SX; COX-2 enzyme inhibition assay, HG and SX; Resources, YEA, HNZ, IM, LAB and PCD; Library creation, SX, JZ and MW; Web app creation, SX, WDGN and MW; Writing – original draft, SX and PCD; Writing – review & editing, all authors; Supervision and funding acquisition, PCD.

## Extended Data Figures

**Extended Data Fig. 1.**
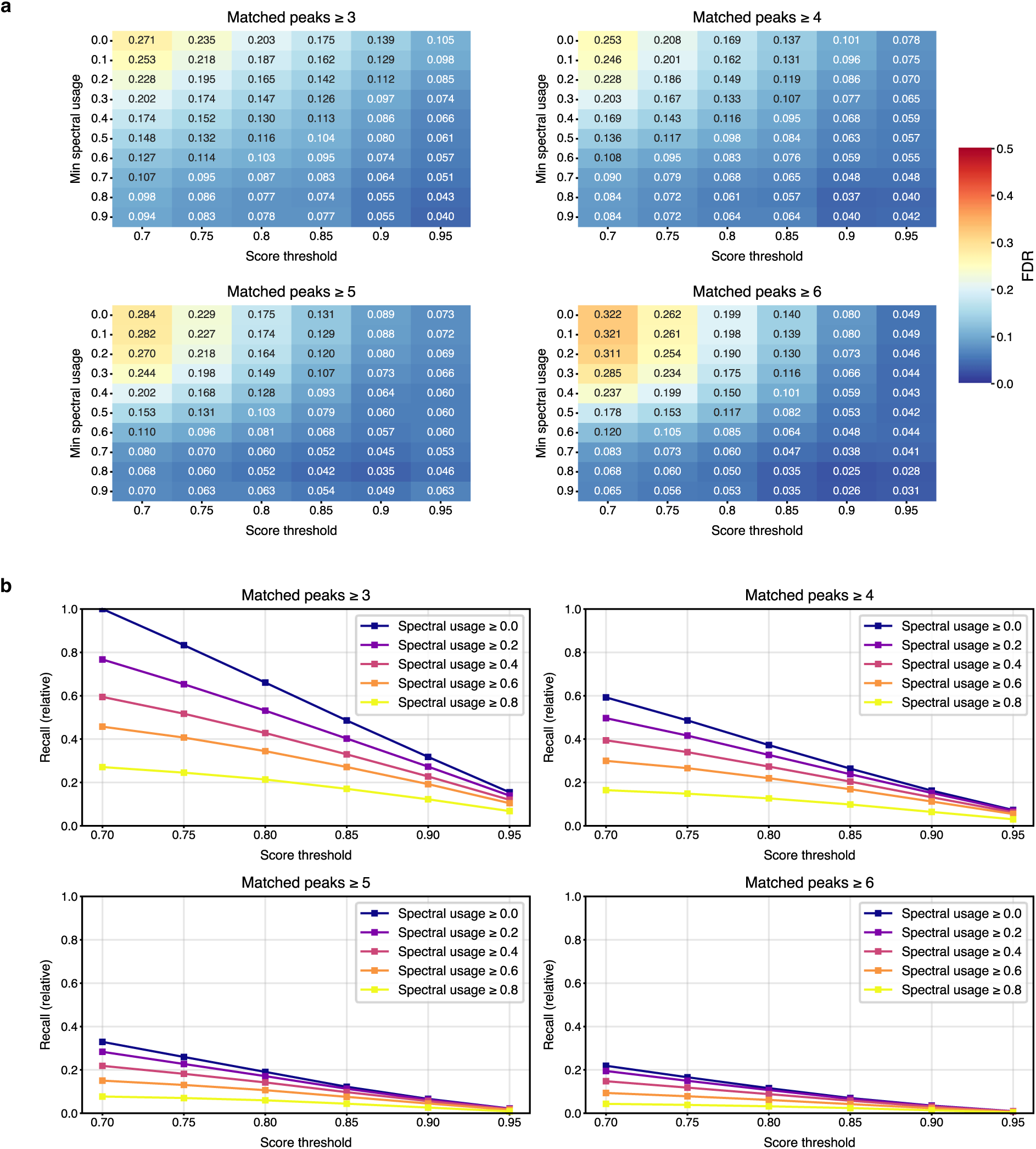
Evaluation of reverse cosine on substructure annotation. **a**) Overall false discovery rate (FDR) under different thresholds of score, matched peak and spectral usage. **b**) Recall performance. Recall rates were calculated relative to the number of positive matches obtained under thresholds of minimum score 0.7, minimum matched peaks 3 and minimum usage 0.0.

**Extended Data Fig. 2.**
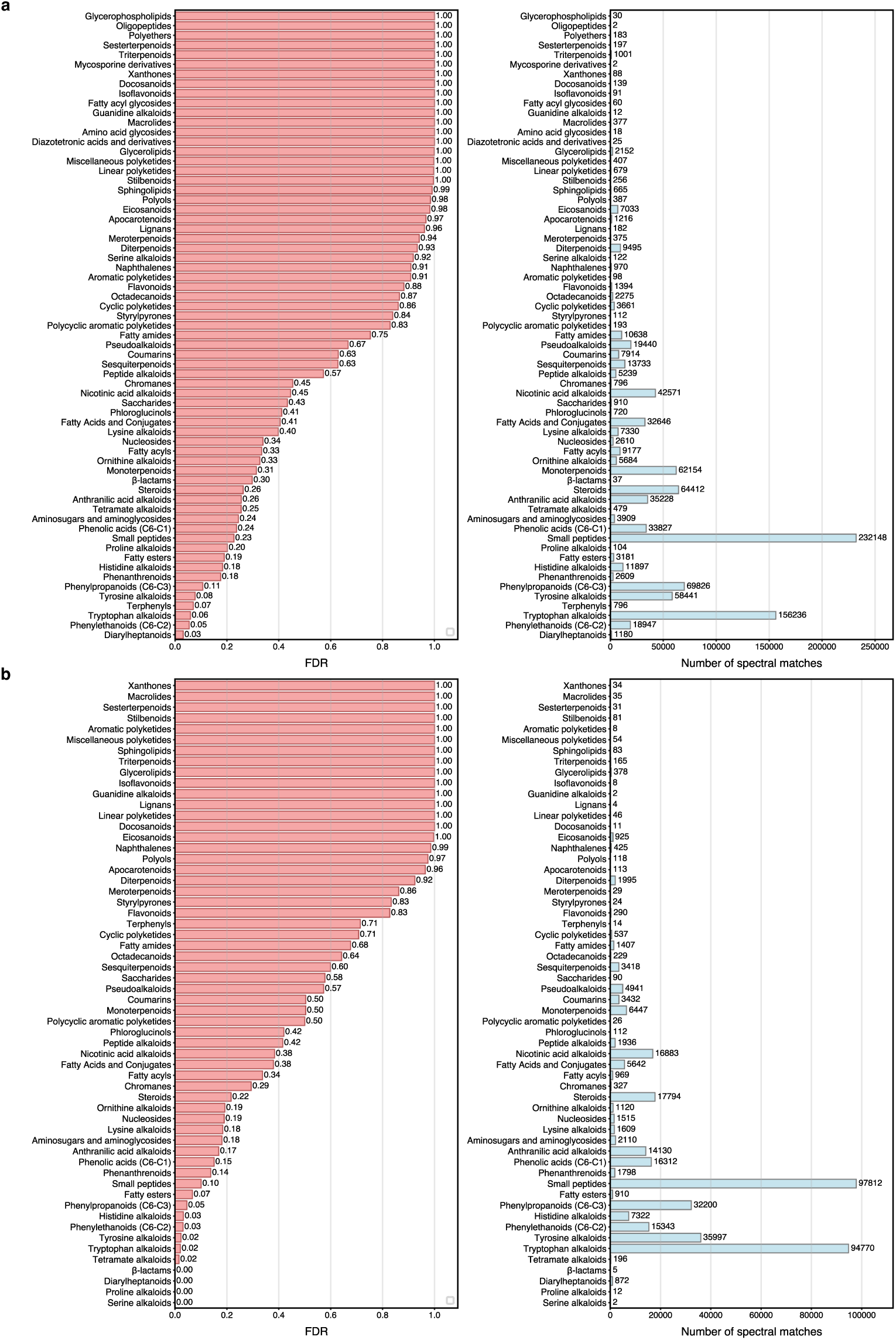
FDR evaluation of reverse cosine on different compound classes. **a**) The following matching thresholds were applied: minimum score of 0.7, minimum matched peak of 3 and spectral usage of 0%. **b**) The following matching thresholds were applied: minimum score of 0.8, minimum matched peak of 3 and spectral usage of 30%. NPClassifier superclass was used.

**Extended Data Fig. 3.**
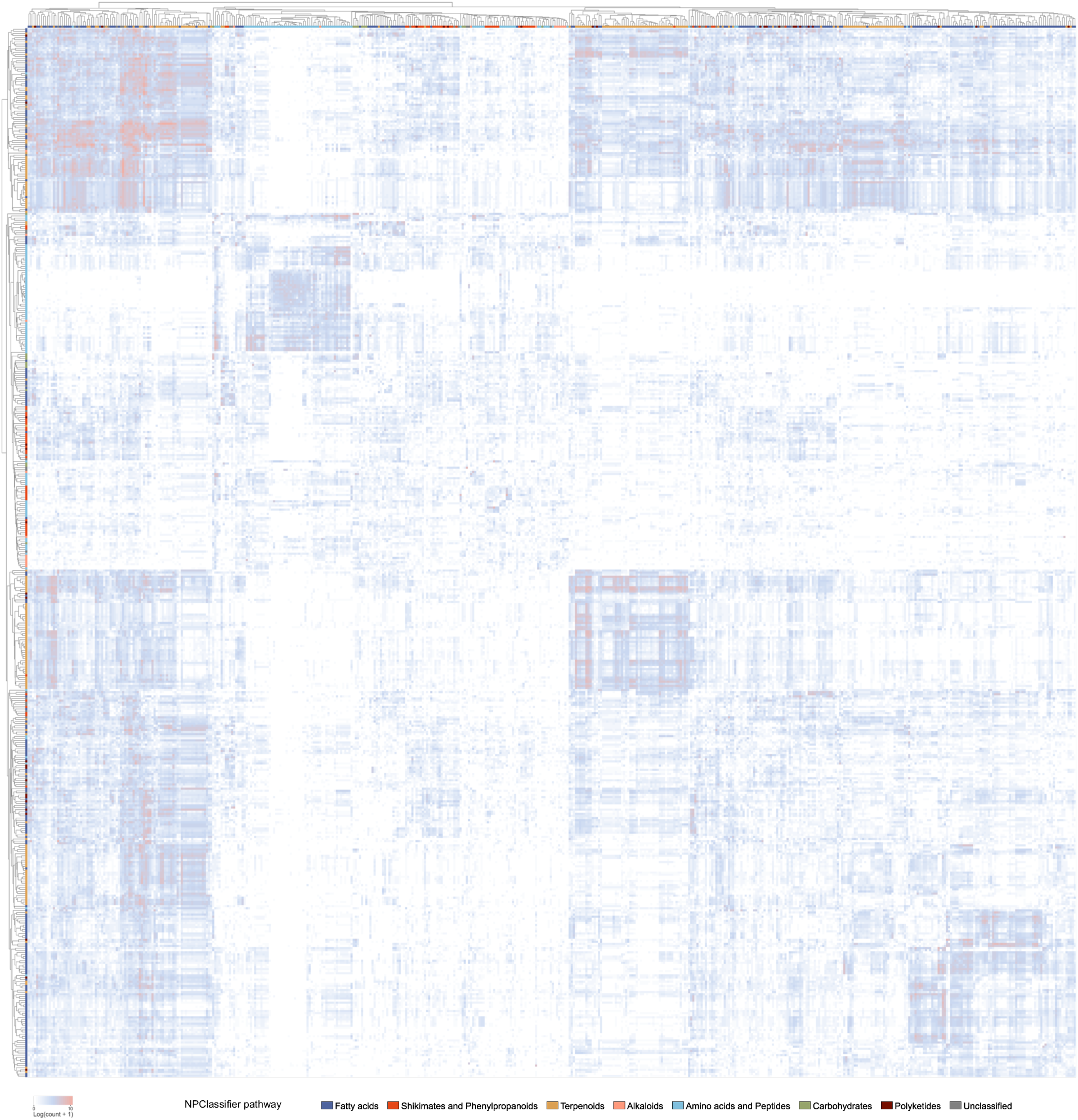
Metabolite conjugations among the 500 most frequently molecules matched by MS/MS. Each row and each column is color coded by compound class information (NPClassifier pathway).

**Extended Data Fig. 4.**
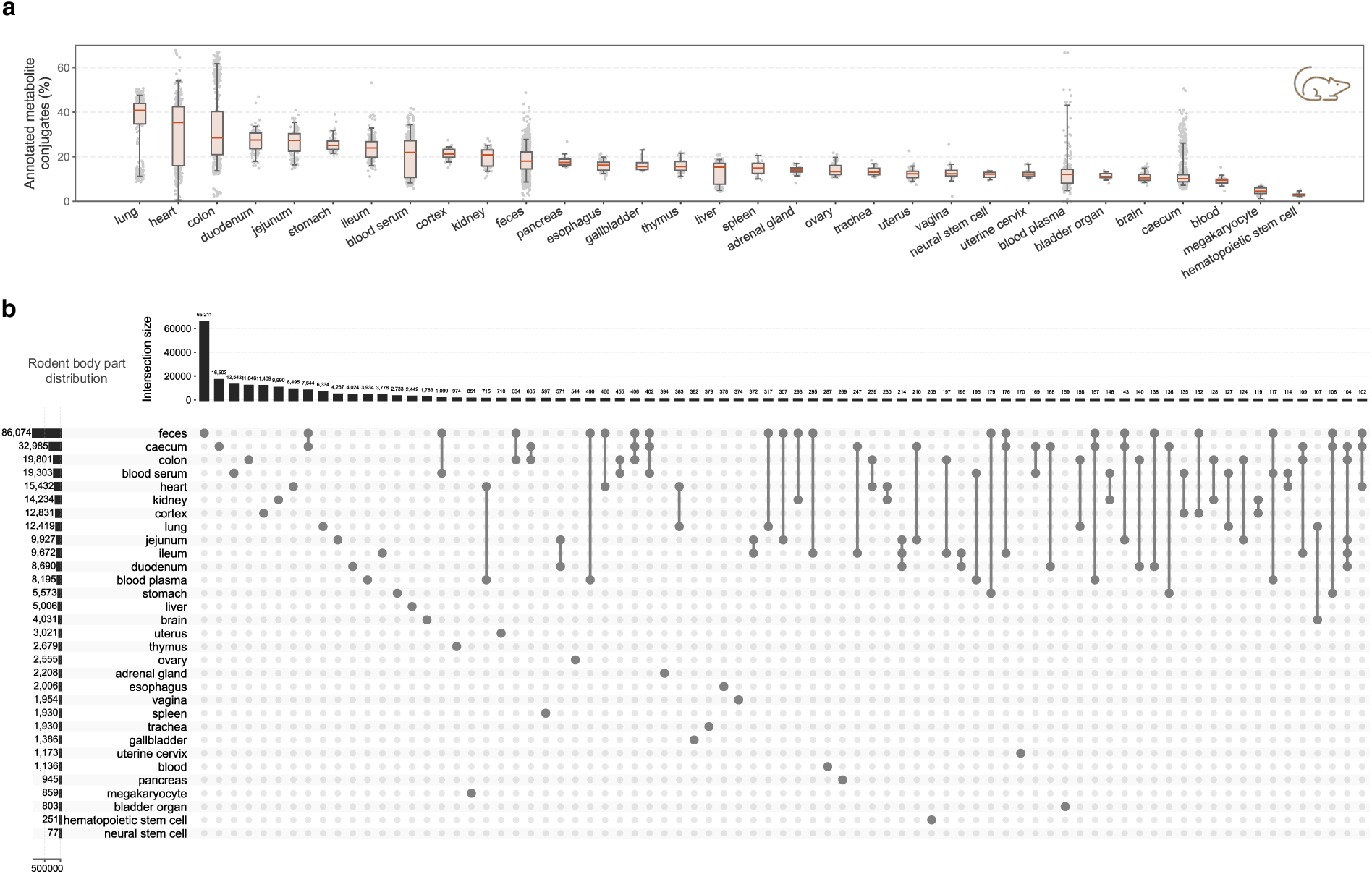
Metabolite conjugates in rodents. **a**) Percentages of annotated metabolite conjugates in mouse samples. Interquartile ranges with median values are shown. Bars show the 5th percentile and the 95th percentile. **b**) Distribution of metabolite conjugates in mouse body sites. Intersections with a size of more than 100 are shown.

**Extended Data Fig. 5.**
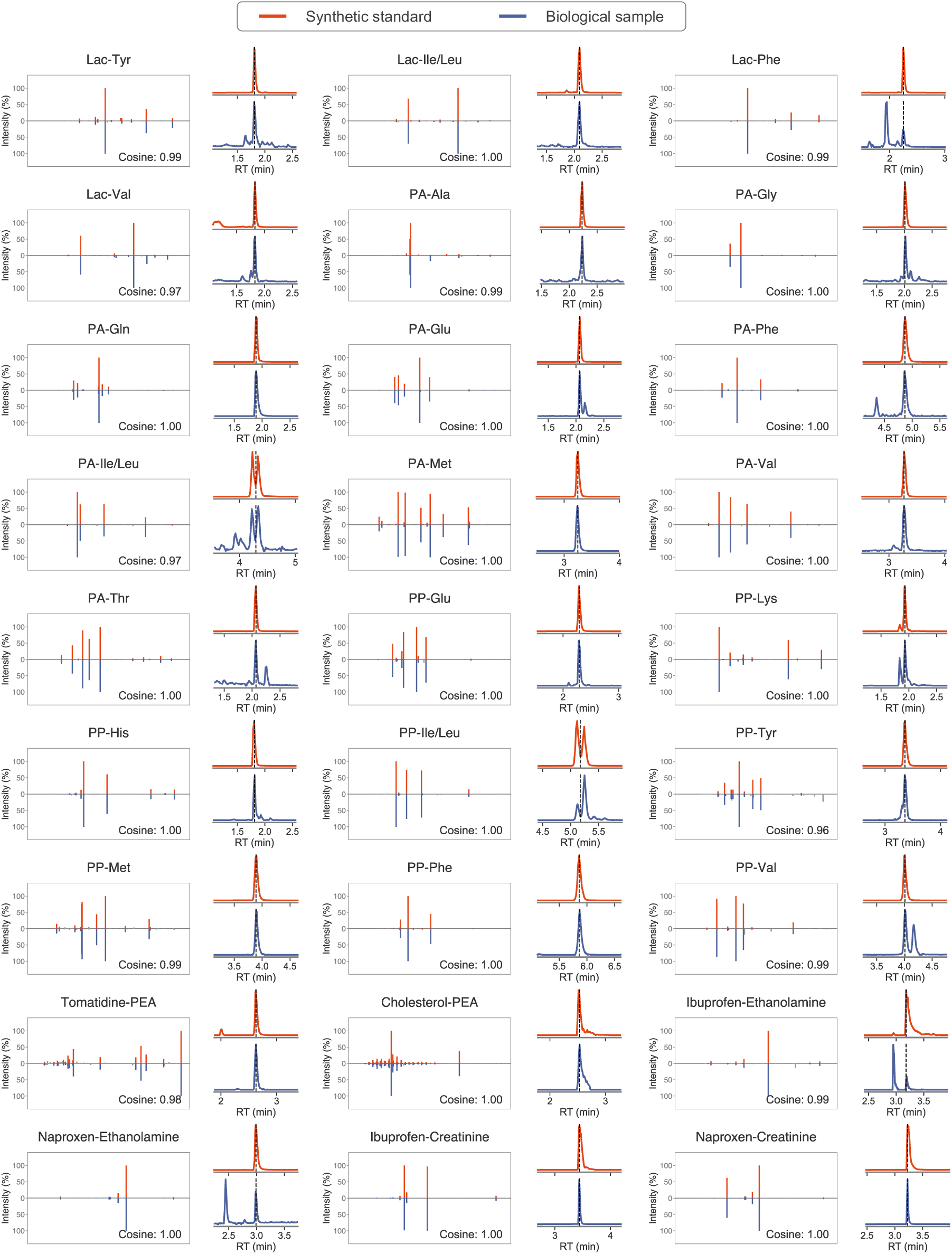
MS/MS and retention time matching of 27 metabolite conjugates (level 1 confidence) between synthetic standards and biological samples. Lac, lactic acid; PA, phenylacetic acid; PP, phenylpropionic acid; PEA, phosphoethanolamine.

**Extended Data Fig. 6.**
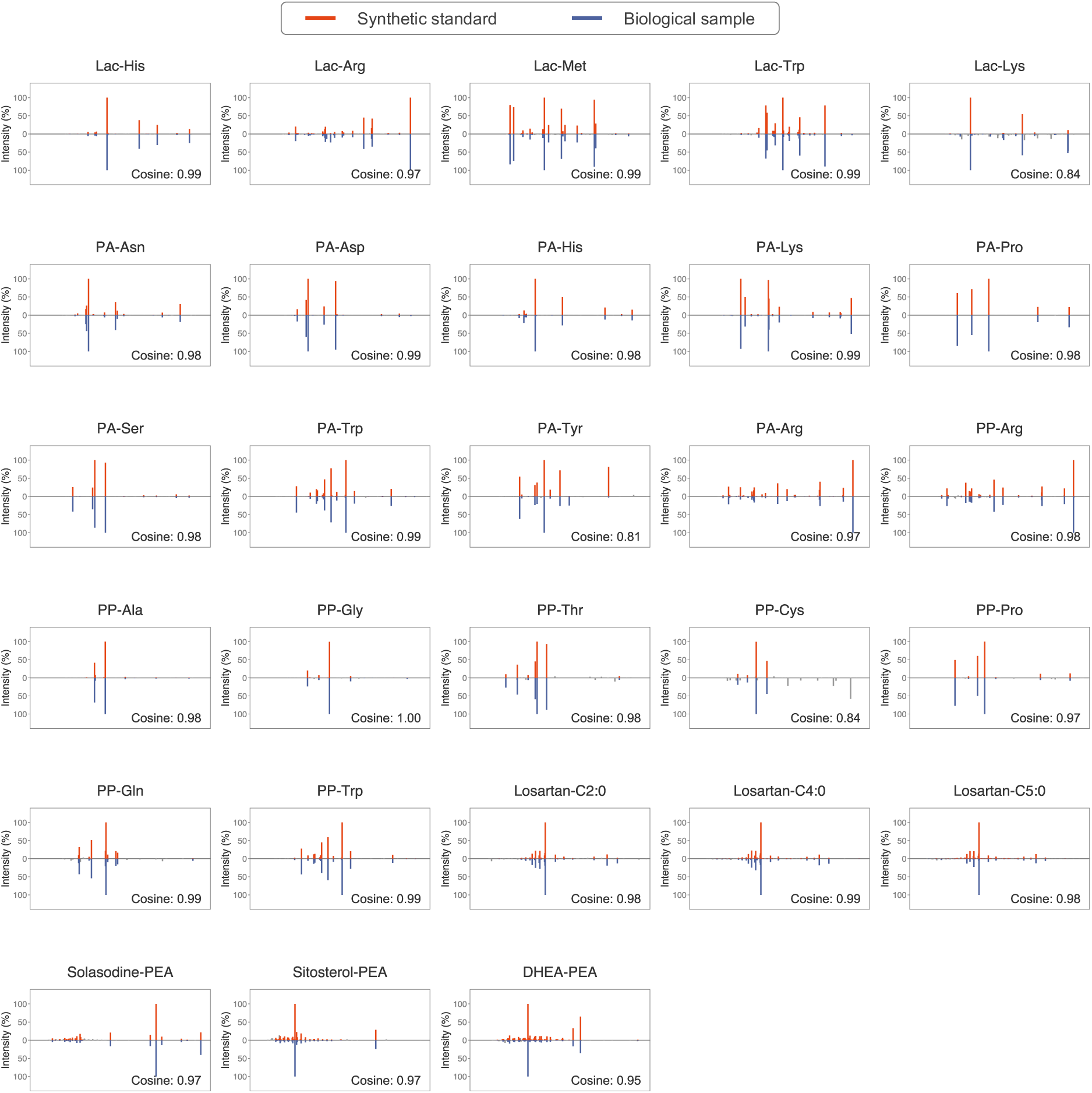
28 metabolite conjugates matched via MS/MS matching (level 2 confidence) between synthetic standards and biological samples. Lac, lactic acid; PA, phenylacetic acid; PP, phenylpropionic acid; PEA, phosphoethanolamine.

**Extended Data Fig. 7.**
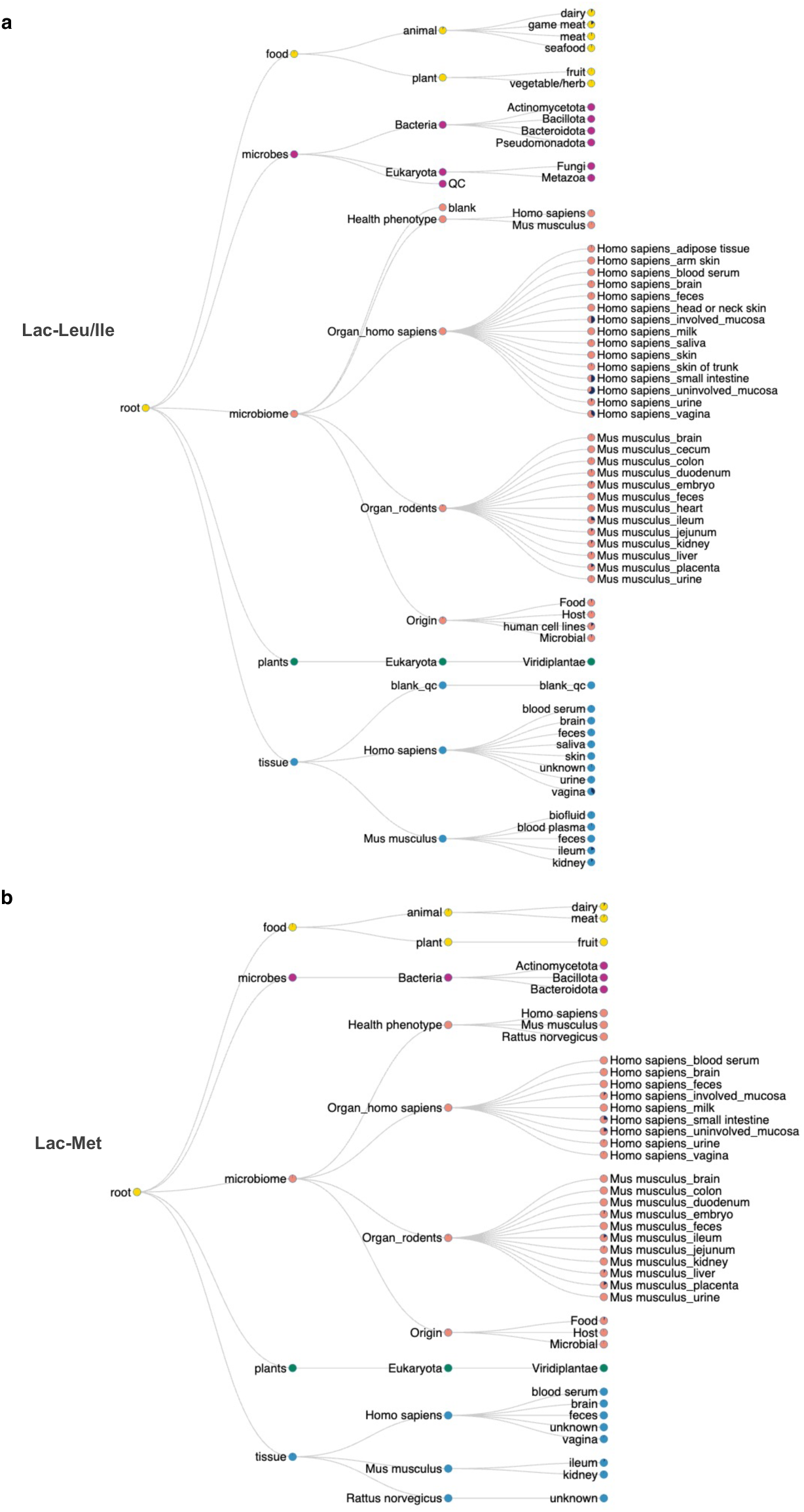
Widespread lactic acid-(iso)leucine and lactic acid-methionine in food, microbes, humans, rodents and plants.

**Extended Data Fig. 8.**
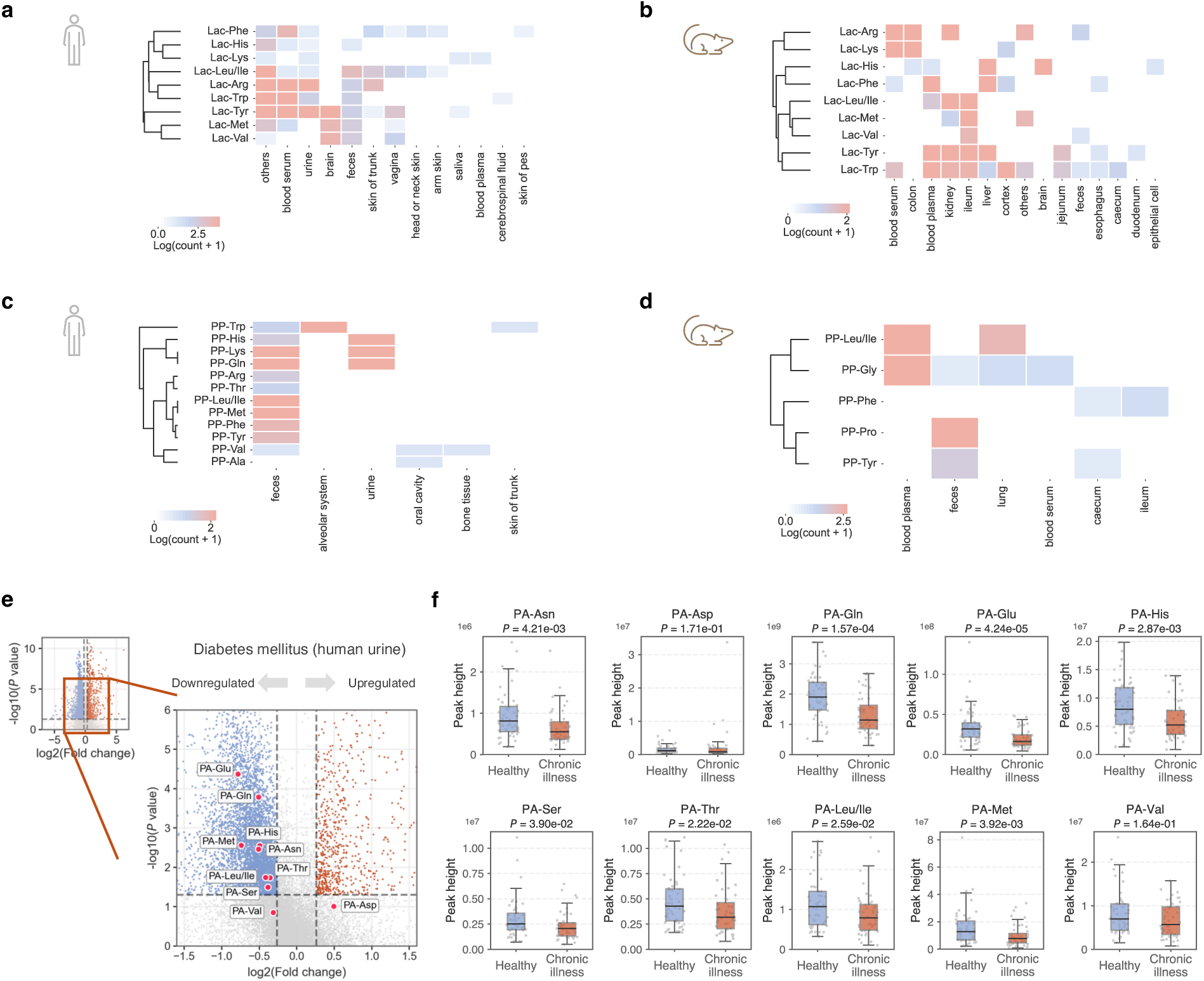
Lactic acid (Lac)-, phenylacetic acid (PA)- and phenylpropionic acid (PP)-amino acid conjugates detected in humans and rodents. **a**) Lac-amino acid conjugates in human body parts. **b**) Lac-amino acid conjugates in rodent body parts. **c**) PP-amino acid conjugates in human body parts. **d**) PP-amino acid conjugates in rodent body parts. **e**) Volcano plot of metabolite features showing dysregulated PA-amino acid conjugates in the diabetes mellitus dataset (human urine samples, MSV000082261). Dashed lines in the volcano plot represent thresholds of fold change 1.2 and *P* value 0.05. **f**) Boxplots of ten detected PA-amino acid conjugates in the diabetes mellitus dataset. Healthy, n = 51; Chronic illness, n = 44. Two-sided Mann-Whitney U test applied. Interquartile ranges (IQR) with median values are shown. Bars show the 75th percentile + 1.5 × IQR and the 25th percentile − 1.5 × IQR.

**Extended Data Fig. 9.**
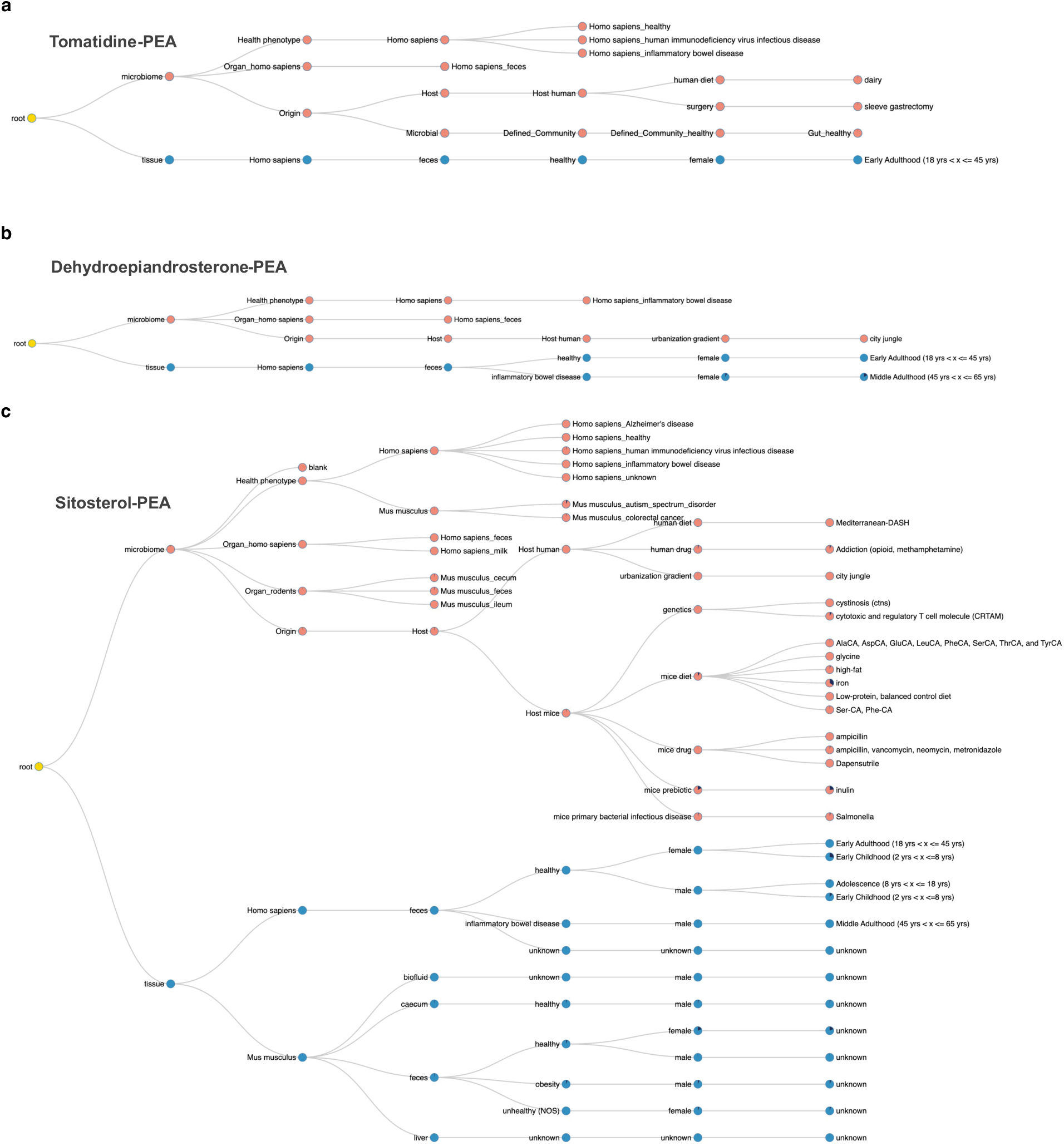
Detection of tomatidine-PEA, dehydroepiandrosterone-PEA and sitosterol-PEA across public metabolomics repositories. **a**) The MS/MS of Tomatidine-PEA was detected in human samples from and only in feces. **b**) The MS/MS of Dehydroepiandrosterone-PEA was also found only in human feces. **c**) The MS/MS of Sitosterol-PEA was primarily detected in human and rodent feces but also in the mouse caecum and liver.

**Extended Data Fig. 10.**
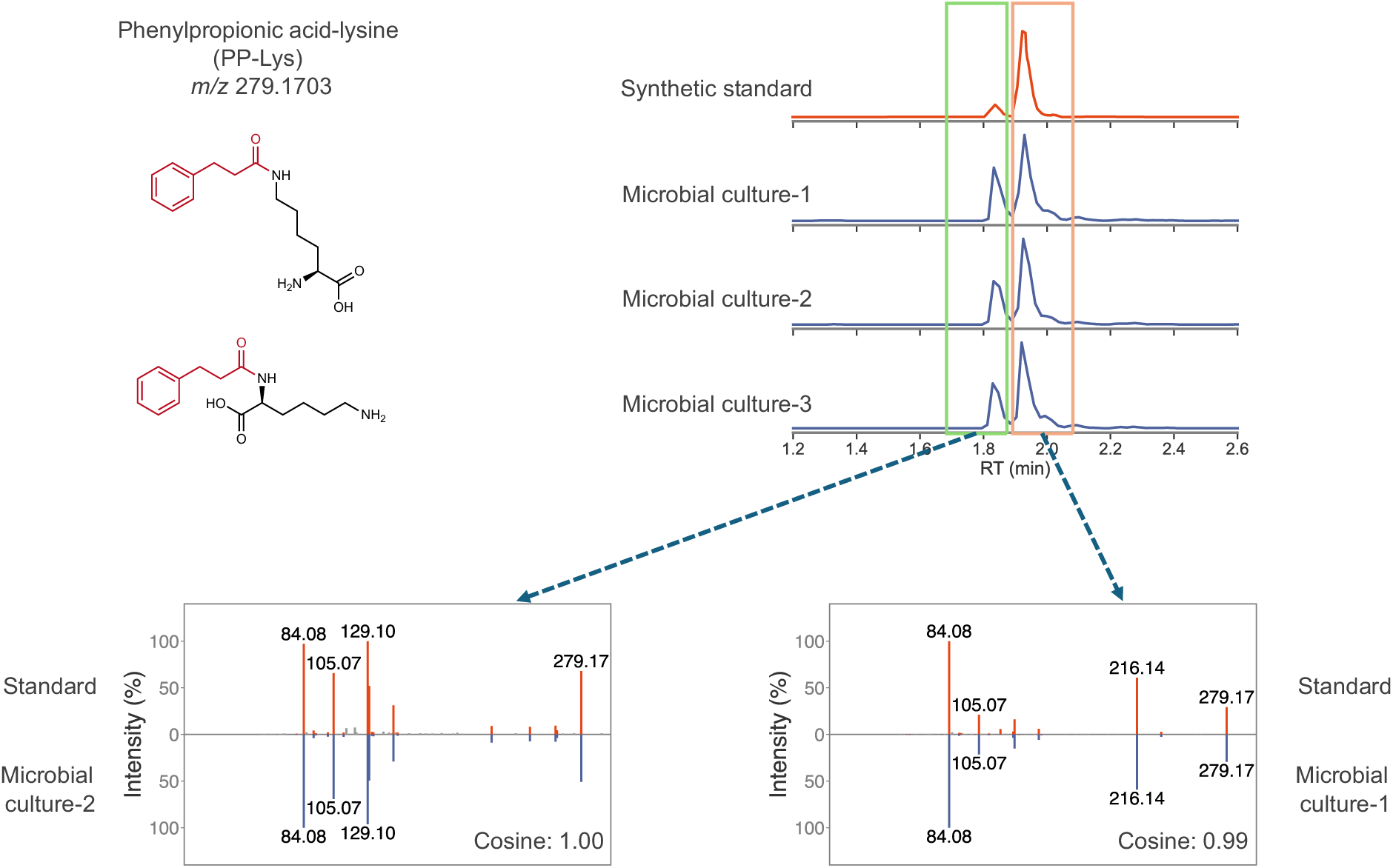
Phenylpropionic acid-lysine as an example of isomeric conjugates. Lysine contains two primary amine groups that can conjugate with phenylpropionic acid. In both synthetic standard mixture and microbial culture samples, two distinct chromatographic peaks were observed corresponding to the mass of phenylpropionic acid-lysine. MS/MS matching confirmed that they are isomers of phenylpropionic acid-lysine: although the MS/MS from the 2 peaks share common fragments of *m/z* 84.08, *m/z* 105.07 and *m/z* 129.10, they have markedly different intensity patterns. This indicates that there might be more unique structures than what reverse spectral search discovered due to multiple conjugation sites.

## References

1. Rodríguez del Río, Á., et al. Functional and evolutionary significance of unknown genes from uncultivated taxa. Nature 626, 377–384 (2024).

2. Pavlopoulos, G. A. et al. Unraveling the functional dark matter through global metagenomics. Nature 1–9 (2023) doi:10.1038/s41586-023-06583-7.

3. Durairaj, J. et al. Uncovering new families and folds in the natural protein universe. Nature 622, 646–653 (2023).

4. Wen, B. et al. DeepMVP: deep learning models trained on high-quality data accurately predict PTM sites and variant-induced alterations. Nat. Methods 22, 1857–1867 (2025).

5. Dai, C. et al. quantms: a cloud-based pipeline for quantitative proteomics enables the reanalysis of public proteomics data. Nat. Methods 21, 1603–1607 (2024).

6. Gentry, E. C. et al. Reverse metabolomics for the discovery of chemical structures from humans. Nature 1–8 (2023) doi:10.1038/s41586-023-06906-8.

7. Mohanty, I. et al. The underappreciated diversity of bile acid modifications. Cell 187, 1801–1818.e20 (2024).

8. Mannochio-Russo, H. et al. The microbiome diversifies long- to short-chain fatty acid-derived N-acyl lipids. Cell 188, 4154–4169.e19 (2025).

9. Yurekten, O. et al. MetaboLights: open data repository for metabolomics. Nucleic Acids Res. 52, D640–D646 (2024).

10. Sud, M. et al. Metabolomics Workbench: An international repository for metabolomics data and metadata, metabolite standards, protocols, tutorials and training, and analysis tools. Nucleic Acids Res. 44, D463–D470 (2016).

11. Wang, M. et al. Sharing and community curation of mass spectrometry data with Global Natural Products Social Molecular Networking. Nat. Biotechnol. 34, 828–837 (2016).

12. Abramson, F. P. Automated identification of mass spectra by the reverse search. Anal. Chem. 47, 45–49 (1975).

13. Xing, S. et al. Reverse Spectral Search Reimagined: A Simple but Overlooked Solution for Chimeric Spectral Annotation. Anal. Chem. 97, 17926–17930 (2025).

14. Bittremieux, W. et al. Universal MS/MS Visualization and Retrieval with the Metabolomics Spectrum Resolver Web Service. 2020.05.09.086066 Preprint at 10.1101/2020.05.09.086066 (2020).

15. Bittremieux, W., Laukens, K., Noble, W. S. & Dorrestein, P. C. Large-scale tandem mass spectrum clustering using fast nearest neighbor searching. Rapid Commun. Mass Spectrom. **n/a**, e9153.

16. Patan, A. et al. Charting the Undiscovered Metabolome with Synthetic Multiplexing. 2025.11.18.689170 Preprint at 10.1101/2025.11.18.689170 (2025).

17. Damiani, T. et al. A universal language for finding mass spectrometry data patterns. Nat. Methods 1–8 (2025) doi:10.1038/s41592-025-02660-z.

18. Kim, H. W. et al. NPClassifier: A Deep Neural Network-Based Structural Classification Tool for Natural Products. J. Nat. Prod. 84, 2795–2807 (2021).

19. El Abiead, Y., et al. Enabling pan-repository reanalysis for big data science of public metabolomics data. Nat. Commun. 16, 4838 (2025).

20. Quinn, R. A. et al. Global chemical effects of the microbiome include new bile-acid conjugations. Nature 579, 123–129 (2020).

21. Pristner, M. et al. Neuroactive metabolites and bile acids are altered in extremely premature infants with brain injury. Cell Rep. Med. 5, (2024).

22. Lucas, L. N., et al. Dominant Bacterial Phyla from the Human Gut Show Widespread Ability To Transform and Conjugate Bile Acids. mSystems 6, 10.1128/msystems.00805-21 (2021).

23. Jansen, R. S. et al. N-lactoyl-amino acids are ubiquitous metabolites that originate from CNDP2-mediated reverse proteolysis of lactate and amino acids. Proc. Natl. Acad. Sci. 112, 6601–6606 (2015).

24. Li, V. L. et al. An exercise-inducible metabolite that suppresses feeding and obesity. Nature 606, 785–790 (2022).

25. Cajka, T. et al. Untargeted Metabolomics Identifies N-Lactoyl-Amino Acids as Dose-Responsive Plasma Biomarkers of Metformin Adherence in Type 2 Diabetes. Clin. Pharmacol. Ther. **n/a**,.

26. James, M., Smith, R. L. & Williams, R. T. Conjugates of phenylacetic acid with taurine and other amino acids in various species. Biochem. J. 124, 15P–16P (1971).

27. Pruss, K. M. et al. Host-microbe co-metabolism via MCAD generates circulating metabolites including hippuric acid. Nat. Commun. 14, 512 (2023).

28. Song, X. et al. Dark Reactions in Microdroplets Explain Widespread Artifacts in Metabolomic Profiling. ACS Meas. Sci. Au https://doi.org/10.1021/acsmeasuresciau.5c00146 (2026) doi:10.1021/acsmeasuresciau.5c00146.

29. Wang, M. et al. Mass spectrometry searches using MASST. Nat. Biotechnol. 38, 23–26 (2020).

30. Mongia, M. et al. Fast mass spectrometry search and clustering of untargeted metabolomics data. Nat. Biotechnol. 42, 1672–1677 (2024).

31. Charron-Lamoureux, V. et al. A guide to reverse metabolomics—a framework for big data discovery strategy. Nat. Protoc. 1–34 (2025) doi:10.1038/s41596-024-01136-2.

32. Zuffa, S. et al. microbeMASST: a taxonomically informed mass spectrometry search tool for microbial metabolomics data. Nat. Microbiol. 1–10 (2024) doi:10.1038/s41564-023-01575-9.

33. West, K. A., Schmid, R., Gauglitz, J. M., Wang, M. & Dorrestein, P. C. foodMASST a mass spectrometry search tool for foods and beverages. Npj Sci. Food 6, 22 (2022).

34. Gomes, P. W. P. et al. plantMASST - Community-driven chemotaxonomic digitization of plants. 2024.05.13.593988 Preprint at 10.1101/2024.05.13.593988 (2024).

35. Zuffa, S. et al. A Multi-Organ Murine Metabolomics Atlas Reveals Molecular Dysregulations in Alzheimer’s Disease. 2025.04.28.651123 Preprint at 10.1101/2025.04.28.651123 (2025).

36. Charron-Lamoureux, V. et al. A searchable metadata network graph for microbiome metabolomics. 2026.02.04.703849 Preprint at 10.64898/2026.02.04.703849 (2026).

37. Zhu, Y. et al. Two distinct gut microbial pathways contribute to meta-organismal production of phenylacetylglutamine with links to cardiovascular disease. Cell Host Microbe 31, 18–32.e9 (2023).

38. Gregor, R. et al. Mammalian gut metabolomes mirror microbiome composition and host phylogeny. ISME J. 16, 1262–1274 (2022).

39. Cheng, A. G. et al. Design, construction, and in vivo augmentation of a complex gut microbiome. Cell 185, 3617–3636.e19 (2022).

40. Xing, S., Shen, S., Xu, B., Li, X. & Huan, T. BUDDY: molecular formula discovery via bottom-up MS/MS interrogation. Nat. Methods 1–10 (2023) doi:10.1038/s41592-023-01850-x.

41. Kim, S. et al. PubChem 2019 update: improved access to chemical data. Nucleic Acids Res. 47, D1102–D1109 (2019).

42. Tsugawa, H. et al. A lipidome atlas in MS-DIAL 4. Nat. Biotechnol. 38, 1159–1163 (2020).

43. Tripathi, A. et al. Chemically informed analyses of metabolomics mass spectrometry data with Qemistree. Nat. Chem. Biol. 17, 146–151 (2021).

44. Schymanski, E. L. et al. Identifying Small Molecules via High Resolution Mass Spectrometry: Communicating Confidence. Environ. Sci. Technol. 48, 2097–2098 (2014).

45. Poli, A. et al. Phytosterols, Cholesterol Control, and Cardiovascular Disease. Nutrients 13, 2810 (2021).

46. Liou, C. S. et al. Gut microbiota gate host exposure to metabolites from dietary Solanums. 2024.03.20.584512 Preprint at 10.1101/2024.03.20.584512 (2024).

47. McGrath, N. A., Brichacek, M. & Njardarson, J. T. A Graphical Journey of Innovative Organic Architectures That Have Improved Our Lives. J. Chem. Educ. 87, 1348–1349 (2010).

48. Alonen, A. et al. Enzyme-assisted synthesis and structure characterization of glucuronic acid conjugates of losartan, candesartan, and zolarsartan. Bioorganic Chem. 36, 148–155 (2008).

49. Seal, S. et al. Cell Painting: a decade of discovery and innovation in cellular imaging. Nat. Methods 22, 254–268 (2025).

50. Al-Mehdi, A.-B. et al. Perinuclear Mitochondrial Clustering Creates an Oxidant-Rich Nuclear Domain Required for Hypoxia-Induced Transcription. Sci. Signal. 5, ra47–ra47 (2012).

51. Zhang, X., Liu, X., Gong, T., Sun, X. & Zhang, Z. In vitro and in vivo investigation of dexibuprofen derivatives for CNS delivery. Acta Pharmacol. Sin. 33, 279–288 (2012).

52. Mahmoud, S. & Mohammad, A. Brain-Specific Delivery of Naproxen Using Different Carrier Systems. Arch. Pharm. (Weinheim) 343, 639–647 (2010).

53. Culp, E. J., Nelson, N. T., Verdegaal, A. A. & Goodman, A. L. Microbial transformation of dietary xenobiotics shapes gut microbiome composition. Cell 187, 6327–6345.e20 (2024).

54. Funabashi, M. et al. A metabolic pathway for bile acid dehydroxylation by the gut microbiome. Nature 582, 566–570 (2020).

55. Zimmermann, M., Zimmermann-Kogadeeva, M., Wegmann, R. & Goodman, A. L. Mapping human microbiome drug metabolism by gut bacteria and their genes. Nature 570, 462–467 (2019).

56. Bushuiev, R. et al. Self-supervised learning of molecular representations from millions of tandem mass spectra using DreaMS. Nat. Biotechnol. 1–11 (2025) doi:10.1038/s41587-025-02663-3.

57. Goldman, S. et al. Annotating metabolite mass spectra with domain-inspired chemical formula transformers. Nat. Mach. Intell. 1–15 (2023) doi:10.1038/s42256-023-00708-3.

58. Huber, F., van der Burg, S., van der Hooft, J. J. J. & Ridder, L. MS2DeepScore: a novel deep learning similarity measure to compare tandem mass spectra. J. Cheminformatics 13, 84 (2021).

59. Qiang, H. et al. Language model-guided anticipation and discovery of mammalian metabolites. Nature 1–10 (2026) doi:10.1038/s41586-025-09969-x.

60. Mohanty, I. et al. MS/MS Mass Spectrometry Filtering Tree for Bile Acid Isomer Annotation. 2025.03.04.641505 Preprint at 10.1101/2025.03.04.641505 (2025).

61. El Abiead, Y., et al. Heterogeneous multimeric metabolite ion species observed in LC-MS based metabolomics data sets. Anal. Chim. Acta 1229, 340352 (2022).

62. Li, Y. & Fiehn, O. Flash entropy search to query all mass spectral libraries in real time. Nat. Methods 20, 1475–1478 (2023).

63. Djoumbou Feunang, Y., et al. ClassyFire: automated chemical classification with a comprehensive, computable taxonomy. J. Cheminformatics 8, 61 (2016).

64. Kretschmer, F., Seipp, J., Ludwig, M., Klau, G. W. & Böcker, S. Coverage bias in small molecule machine learning. Nat. Commun. 16, 554 (2025).

65. Bray, M.-A. et al. Cell Painting, a high-content image-based assay for morphological profiling using multiplexed fluorescent dyes. Nat. Protoc. 11, 1757–1774 (2016).

66. Carpenter, A. E. et al. CellProfiler: image analysis software for identifying and quantifying cell phenotypes. Genome Biol. 7, R100 (2006).

