## Supplementary material for "Navigating the conjugated metabolome": SI_file

**Comparisons of predicted substructures obtained with reverse cosine and top 1000 unique annotations obtained with MS/MS matches to the recently generated multiplexed synthetic reference library.** For the structures in that library when two molecules are coupled only one representative structure is shown and other structural isomers may sometimes be possible. Dashed lines are the precursor ions; red spectra show synthetic standards and blue spectra show matched biological MS/MS spectra with reverse cosine predictions.

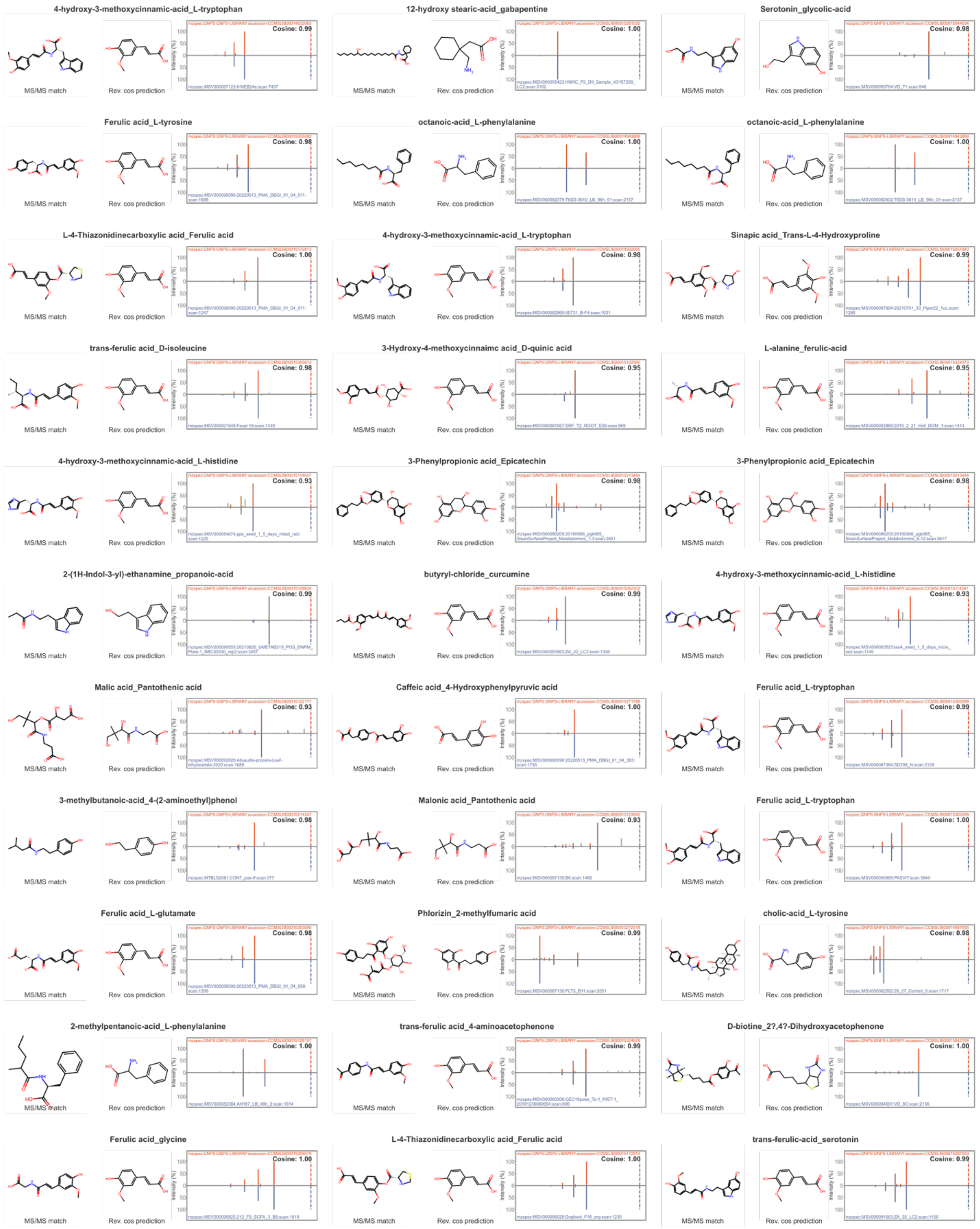
4-hydroxy-3-methoxycinnamic-acid_L-tryptophan
MS/MS match
Rev. cos prediction
Cosine: 0.99
12-hydroxy stearic-acid_gabapentine
MS/MS match
Rev. cos prediction
Cosine: 1.00
Serotonin_glycolic-acid
MS/MS match
Rev. cos prediction
Cosine: 0.98
Ferulic acid_L-tyrosine
MS/MS match
Rev. cos prediction
Cosine: 0.98
octanoic-acid_L-phenylalanine
MS/MS match
Rev. cos prediction
Cosine: 1.00
octanoic-acid_L-phenylalanine
MS/MS match
Rev. cos prediction
Cosine: 1.00
L-4-Thiazonidinecarboxylic acid_Ferulic acid
MS/MS match
Rev. cos prediction
Cosine: 1.00
4-hydroxy-3-methoxycinnamic-acid_L-tryptophan
MS/MS match
Rev. cos prediction
Cosine: 0.98
Sinapic acid_Trans-L-4-Hydroxyproline
MS/MS match
Rev. cos prediction
Cosine: 0.99
trans-ferulic acid_D-isoleucine
MS/MS match
Rev. cos prediction
Cosine: 0.98
3-Hydroxy-4-methoxycinnaimc acid_D-quinic acid
MS/MS match
Rev. cos prediction
Cosine: 0.95
L-alanine_ferulic-acid
MS/MS match
Rev. cos prediction
Cosine: 0.95
4-hydroxy-3-methoxycinnamic-acid_L-histidine
MS/MS match
Rev. cos prediction
Cosine: 0.93
3-Phenylpropionic acid_Epicatechin
MS/MS match
Rev. cos prediction
Cosine: 0.98
3-Phenylpropionic acid_Epicatechin
MS/MS match
Rev. cos prediction
Cosine: 0.98
2-(1H-Indol-3-yl)-ethanamine_propanoic-acid
MS/MS match
Rev. cos prediction
Cosine: 0.99
butyryl-chloride_curcumine
MS/MS match
Rev. cos prediction
Cosine: 0.99
4-hydroxy-3-methoxycinnamic-acid_L-histidine
MS/MS match
Rev. cos prediction
Cosine: 0.93
Malic acid_Pantothenic acid
MS/MS match
Rev. cos prediction
Cosine: 0.93
Caffeic acid_4-Hydroxyphenylpyruvic acid
MS/MS match
Rev. cos prediction
Cosine: 1.00
Ferulic acid_L-tryptophan
MS/MS match
Rev. cos prediction
Cosine: 0.99
3-methylbutanoic-acid_4-(2-aminoethyl)phenol
MS/MS match
Rev. cos prediction
Cosine: 0.98
Malonic acid_Pantothenic acid
MS/MS match
Rev. cos prediction
Cosine: 0.93
Ferulic acid_L-tryptophan
MS/MS match
Rev. cos prediction
Cosine: 1.00
Ferulic acid_L-glutamate
MS/MS match
Rev. cos prediction
Cosine: 0.98
Phlorizin_2-methylfumaric acid
MS/MS match
Rev. cos prediction
Cosine: 0.98
cholic-acid_L-tyrosine
MS/MS match
Rev. cos prediction
Cosine: 0.98
2-methylpentanoic-acid_L-phenylalanine
MS/MS match
Rev. cos prediction
Cosine: 1.00
trans-ferulic acid_4-aminoacetophenone
MS/MS match
Rev. cos prediction
Cosine: 0.99
D-biotine_2?,4?-Dihydroxyacetophenone
MS/MS match
Rev. cos prediction
Cosine: 1.00
Ferulic acid_glycine
MS/MS match
Rev. cos prediction
Cosine: 1.00
L-4-Thiazonidinecarboxylic acid_Ferulic acid
MS/MS match
Rev. cos prediction
Cosine: 1.00
trans-ferulic-acid_serotonin
MS/MS match
Rev. cos prediction
Cosine: 0.99

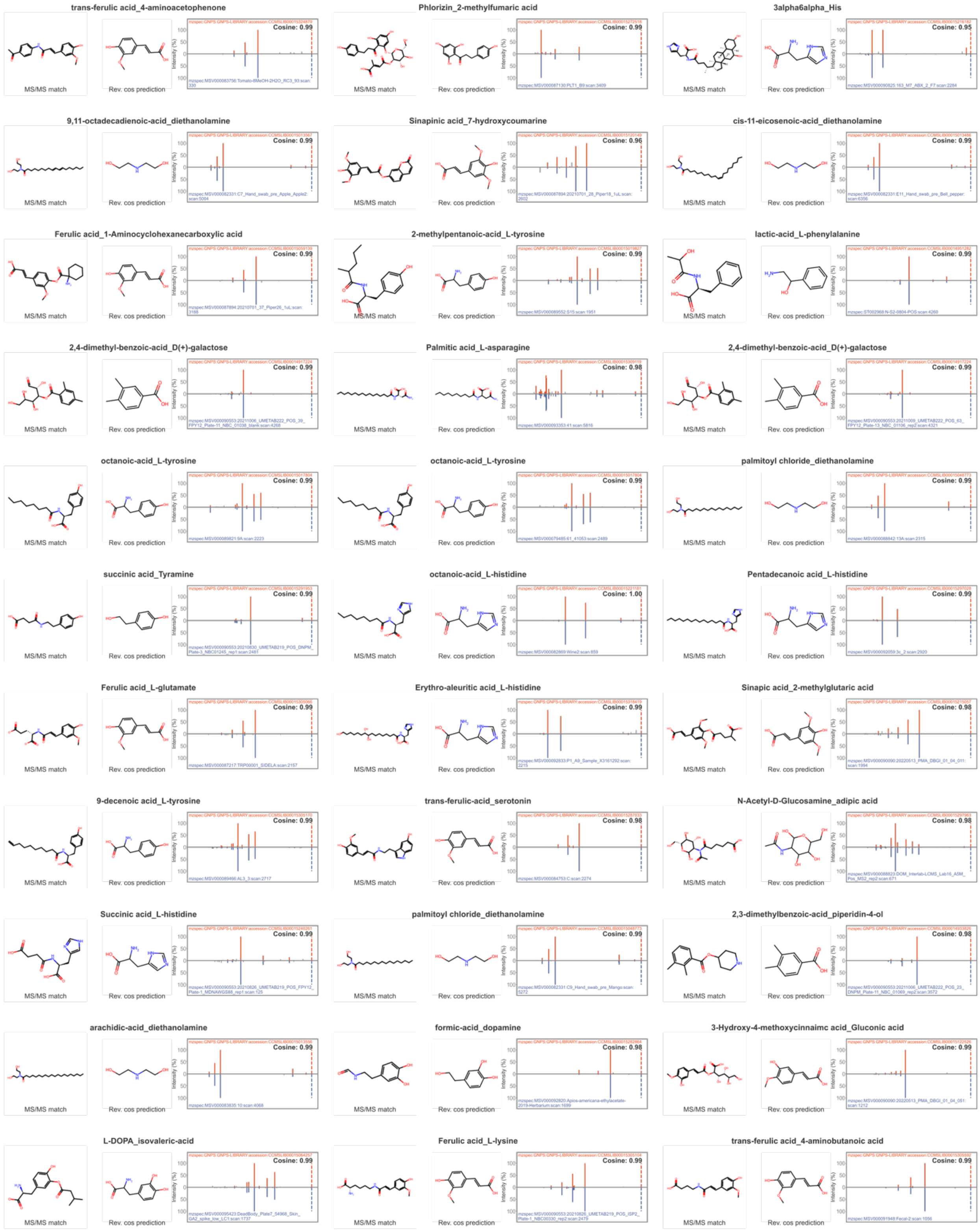

trans-ferulic acid_4-aminoacetophenone
Cosine: 0.99
Phlorizin_2-methylfumaric acid
Cosine: 0.99
3alpha6alpha_His
Cosine: 0.95
9,11-octadecadienoic-acid_diethanolamine
Cosine: 0.99
Sinapinic acid_7-hydroxycoumarine
Cosine: 0.96
cis-11-eicosenoic-acid_diethanolamine
Cosine: 0.99
Ferulic acid_1-Aminocyclohexanecarboxylic acid
Cosine: 0.99
2-methylpentanoic-acid_L-tyrosine
Cosine: 0.99
lactic-acid_L-phenylalanine
Cosine: 0.99
2,4-dimethyl-benzoic-acid_D(+)-galactose
Cosine: 0.99
Palmitic acid_L-asparagine
Cosine: 0.99
2,4-dimethyl-benzoic-acid_D(+)-galactose
Cosine: 0.99
octanoic-acid_L-tyrosine
Cosine: 0.99
octanoic-acid_L-tyrosine
Cosine: 0.99
palmitoyl chloride_diethanolamine
Cosine: 0.99
succinic acid_Tyramine
Cosine: 0.99
octanoic-acid_L-histidine
Cosine: 1.00
Pentadecanoic acid_L-histidine
Cosine: 0.99
Ferulic acid_L-glutamate
Cosine: 0.99
Erythro-aleuritic acid_L-histidine
Cosine: 0.99
Sinapic acid_2-methylglutaric acid
Cosine: 0.98
9-decenoic acid_L-tyrosine
Cosine: 0.99
trans-ferulic-acid_serotonin
Cosine: 0.98
N-Acetyl-D-Glucosamine_adipic acid
Cosine: 0.98
Succinic acid_L-histidine
Cosine: 0.99
palmitoyl chloride_diethanolamine
Cosine: 0.99
2,3-dimethylbenzoic-acid_piperidin-4-ol
Cosine: 0.98
arachidic-acid_diethanolamine
Cosine: 0.99
formic-acid_dopamine
Cosine: 0.98
3-Hydroxy-4-methoxycinnaimc acid_Gluconic acid
Cosine: 0.99
L-DOPA_isovaleric-acid
Cosine: 0.99
Ferulic acid_L-lysine
Cosine: 0.99
trans-ferulic acid_4-aminobutanoic acid
Cosine: 0.99
MS/MS match
Rev. cos prediction
Intensity (%)

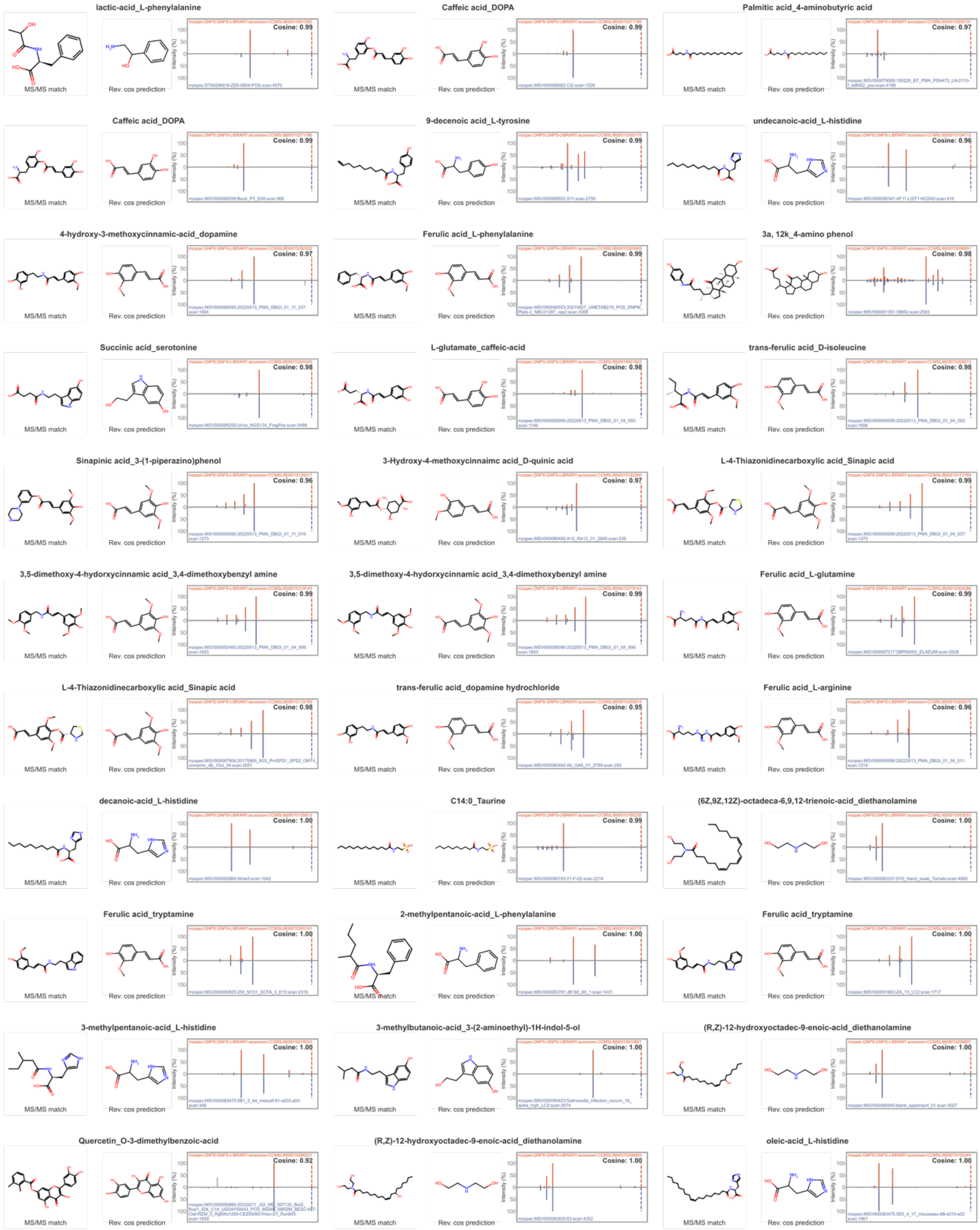
lactic-acid_L-phenylalanine
Cosine: 0.99
Caffeic acid_DOPA
Cosine: 0.99
Palmitic acid_4-aminobutyric acid
Cosine: 0.97
Caffeic acid_DOPA
Cosine: 0.99
9-decenoic acid_L-tyrosine
Cosine: 0.99
undecanoic-acid_L-histidine
Cosine: 0.96
4-hydroxy-3-methoxycinnamic-acid_dopamine
Cosine: 0.97
Ferulic acid_L-phenylalanine
Cosine: 0.98
3a, 12k_4-amino phenol
Cosine: 0.98
Succinic acid_serotonine
Cosine: 0.98
L-glutamate_caffeic-acid
Cosine: 0.98
trans-ferulic acid_D-isoleucine
Cosine: 0.98
Sinapinic acid_3-(1-piperazino)phenol
Cosine: 0.96
3-Hydroxy-4-methoxycinnaimc acid_D-quinic acid
Cosine: 0.97
L-4-Thiazonidinecarboxylic acid_Sinapic acid
Cosine: 0.99
3,5-dimethoxy-4-hydorxycinnamic acid_3,4-dimethoxybenzyl amine
Cosine: 0.99
3,5-dimethoxy-4-hydorxycinnamic acid_3,4-dimethoxybenzyl amine
Cosine: 0.99
Ferulic acid_L-glutamine
Cosine: 0.99
L-4-Thiazonidinecarboxylic acid_Sinapic acid
Cosine: 0.98
trans-ferulic acid_dopamine hydrochloride
Cosine: 0.95
Ferulic acid_L-arginine
Cosine: 0.96
decanoic-acid_L-histidine
Cosine: 1.00
C14:0_Taurine
Cosine: 0.99
(6Z,9Z,12Z)-octadeca-6,9,12-trienoic-acid_diethanolamine
Cosine: 1.00
Ferulic acid_tryptamine
Cosine: 1.00
2-methylpentanoic-acid_L-phenylalanine
Cosine: 1.00
Ferulic acid_tryptamine
Cosine: 1.00
3-methylpentanoic-acid_L-histidine
Cosine: 1.00
3-methylbutanoic-acid_3-(2-aminoethyl)-1H-indol-5-ol
Cosine: 1.00
(R,Z)-12-hydroxyoctadec-9-enoic-acid_diethanolamine
Cosine: 1.00
Quercetin_O-3-dimethylbenzoic-acid
Cosine: 0.92
(R,Z)-12-hydroxyoctadec-9-enoic-acid_diethanolamine
Cosine: 1.00
oleic-acid_L-histidine
Cosine: 1.00
MS/MS match
Rev. cos prediction
Intensity (%)

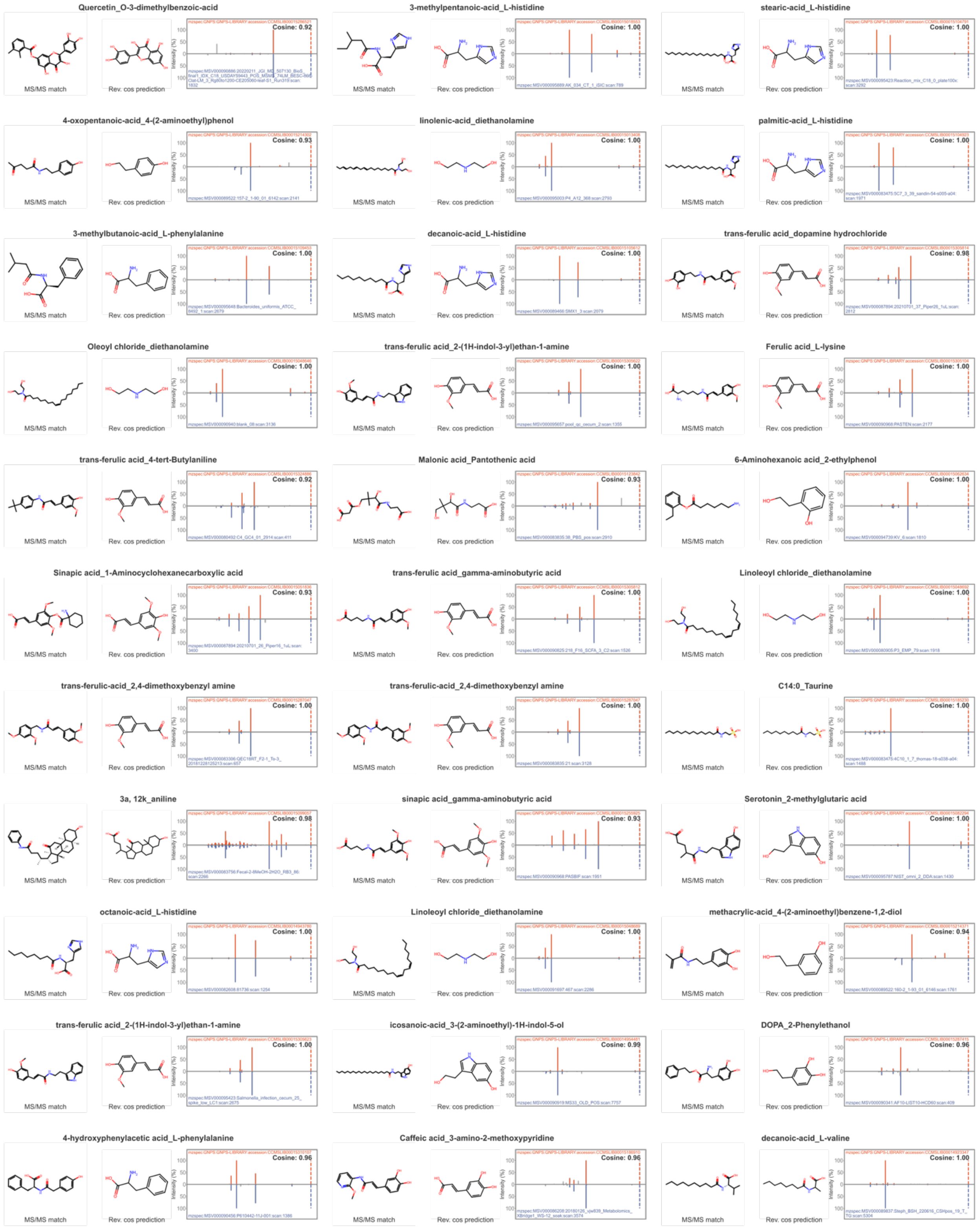
Quercetin_O-3-dimethylbenzoic-acid
Cosine: 0.92
3-methylpentanoic-acid_L-histidine
Cosine: 1.00
stearic-acid_L-histidine
Cosine: 1.00
4-oxopentanoic-acid_4-(2-aminoethyl)phenol
Cosine: 0.93
linolenic-acid_diethanolamine
Cosine: 1.00
palmitic-acid_L-histidine
Cosine: 1.00
3-methylbutanoic-acid_L-phenylalanine
Cosine: 1.00
decanoic-acid_L-histidine
Cosine: 1.00
trans-ferulic acid_dopamine hydrochloride
Cosine: 0.98
Oleoyl chloride_diethanolamine
Cosine: 1.00
trans-ferulic acid_2-(1H-indol-3-yl)ethan-1-amine
Cosine: 1.00
Ferulic acid_L-lysine
Cosine: 1.00
trans-ferulic acid_4-tert-Butylaniline
Cosine: 0.92
Malonic acid_Pantothenic acid
Cosine: 0.93
6-Aminohexanoic acid_2-ethylphenol
Cosine: 1.00
Sinapic acid_1-Aminocyclohexanecarboxylic acid
Cosine: 0.93
trans-ferulic acid_gamma-aminobutyric acid
Cosine: 1.00
Linoleoyl chloride_diethanolamine
Cosine: 1.00
trans-ferulic-acid_2,4-dimethoxybenzyl amine
Cosine: 1.00
trans-ferulic-acid_2,4-dimethoxybenzyl amine
Cosine: 1.00
C14:0_Taurine
Cosine: 1.00
3a, 12k_aniline
Cosine: 0.98
sinapic acid_gamma-aminobutyric acid
Cosine: 0.93
Serotonin_2-methylglutaric acid
Cosine: 1.00
octanoic-acid_L-histidine
Cosine: 1.00
Linoleoyl chloride_diethanolamine
Cosine: 1.00
methacrylic-acid_4-(2-aminoethyl)benzene-1,2-diol
Cosine: 0.94
trans-ferulic acid_2-(1H-indol-3-yl)ethan-1-amine
Cosine: 1.00
icosanoic-acid_3-(2-aminoethyl)-1H-indol-5-ol
Cosine: 0.99
DOPA_2-Phenylethanol
Cosine: 0.96
4-hydroxyphenylacetic acid_L-phenylalanine
Cosine: 0.96
Caffeic acid_3-amino-2-methoxypyridine
Cosine: 0.96
decanoic-acid_L-valine
Cosine: 1.00
MS/MS match
Rev. cos prediction
Intensity (%)

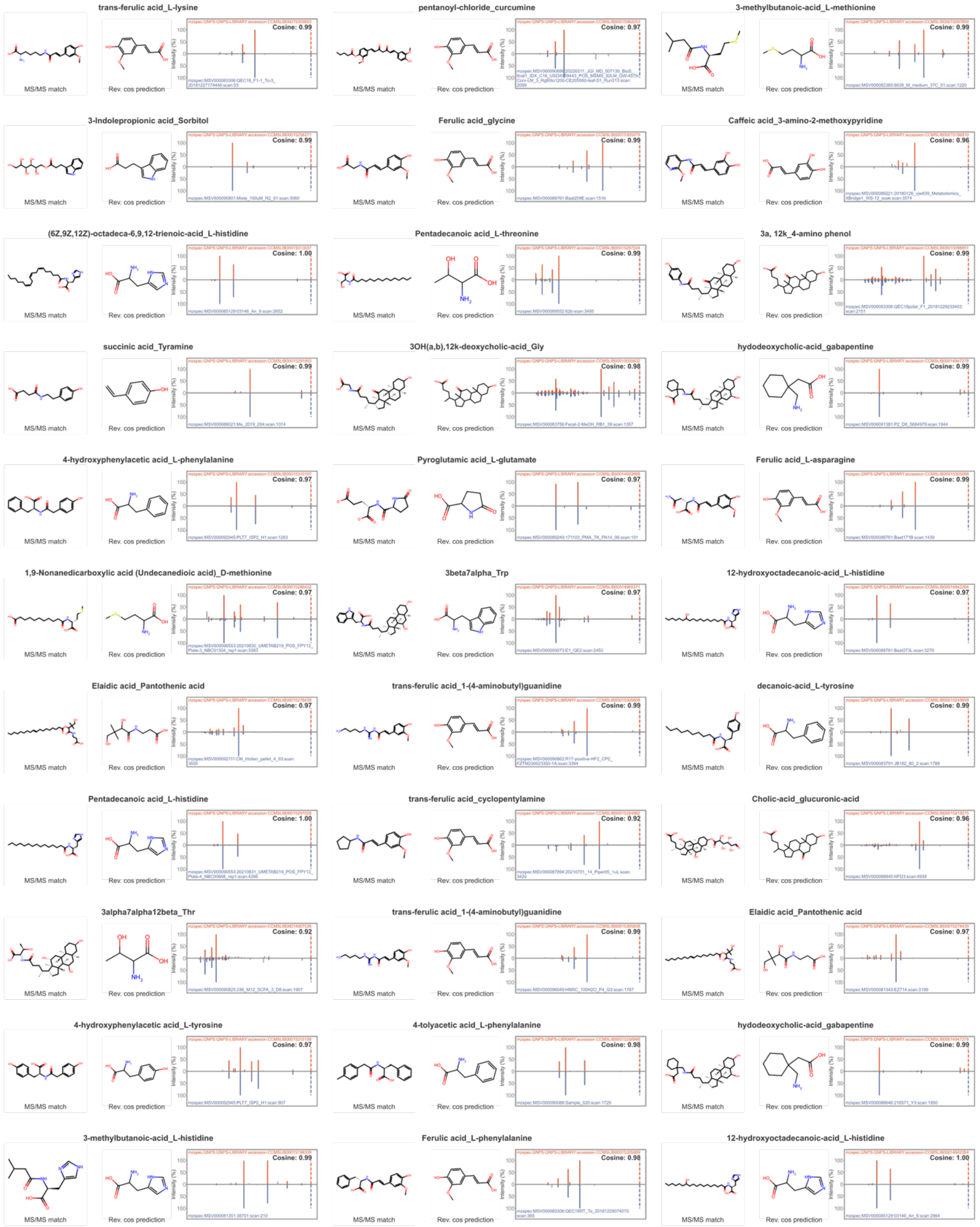
trans-ferulic acid_L-lysine
Cosine: 0.99
MS/MS match
Rev. cos prediction
pentanoyl-chloride_curcumine
Cosine: 0.97
3-methylbutanoic-acid_L-methionine
Cosine: 0.99
3-Indolepropionic acid_Sorbitol
Cosine: 0.99
Ferulic acid_glycine
Cosine: 0.99
Caffeic acid_3-amino-2-methoxypyridine
Cosine: 0.96
(6Z,9Z,12Z)-octadeca-6,9,12-trienoic-acid_L-histidine
Cosine: 1.00
Pentadecanoic acid_L-threonine
Cosine: 0.99
3a, 12k_4-amino phenol
Cosine: 0.99
succinic acid_Tyramine
Cosine: 0.99
3OH(a,b),12k-deoxycholic-acid_Gly
Cosine: 0.98
hydodeoxycholic-acid_gabapentine
Cosine: 0.99
4-hydroxyphenylacetic acid_L-phenylalanine
Cosine: 0.97
Pyroglutamic acid_L-glutamate
Cosine: 0.97
Ferulic acid_L-asparagine
Cosine: 0.99
1,9-Nonanedicarboxylic acid (Undecanedioic acid)_D-methionine
Cosine: 0.97
3beta7alpha_Trp
Cosine: 0.97
12-hydroxyoctadecanoic-acid_L-histidine
Cosine: 0.97
Elaidic acid_Pantothenic acid
Cosine: 0.97
trans-ferulic acid_1-(4-aminobutyl)guanidine
Cosine: 0.99
decanoic-acid_L-tyrosine
Cosine: 0.99
Pentadecanoic acid_L-histidine
Cosine: 1.00
trans-ferulic acid_cyclopentylamine
Cosine: 0.92
Cholic-acid_glucuronic-acid
Cosine: 0.96
3alpha7alpha12beta_Thr
Cosine: 0.92
trans-ferulic acid_1-(4-aminobutyl)guanidine
Cosine: 0.99
Elaidic acid_Pantothenic acid
Cosine: 0.97
4-hydroxyphenylacetic acid_L-tyrosine
Cosine: 0.97
4-tolyacetic acid_L-phenylalanine
Cosine: 0.98
hydodeoxycholic-acid_gabapentine
Cosine: 0.99
3-methylbutanoic-acid_L-histidine
Cosine: 0.99
Ferulic acid_L-phenylalanine
Cosine: 0.98
12-hydroxyoctadecanoic-acid_L-histidine
Cosine: 1.00

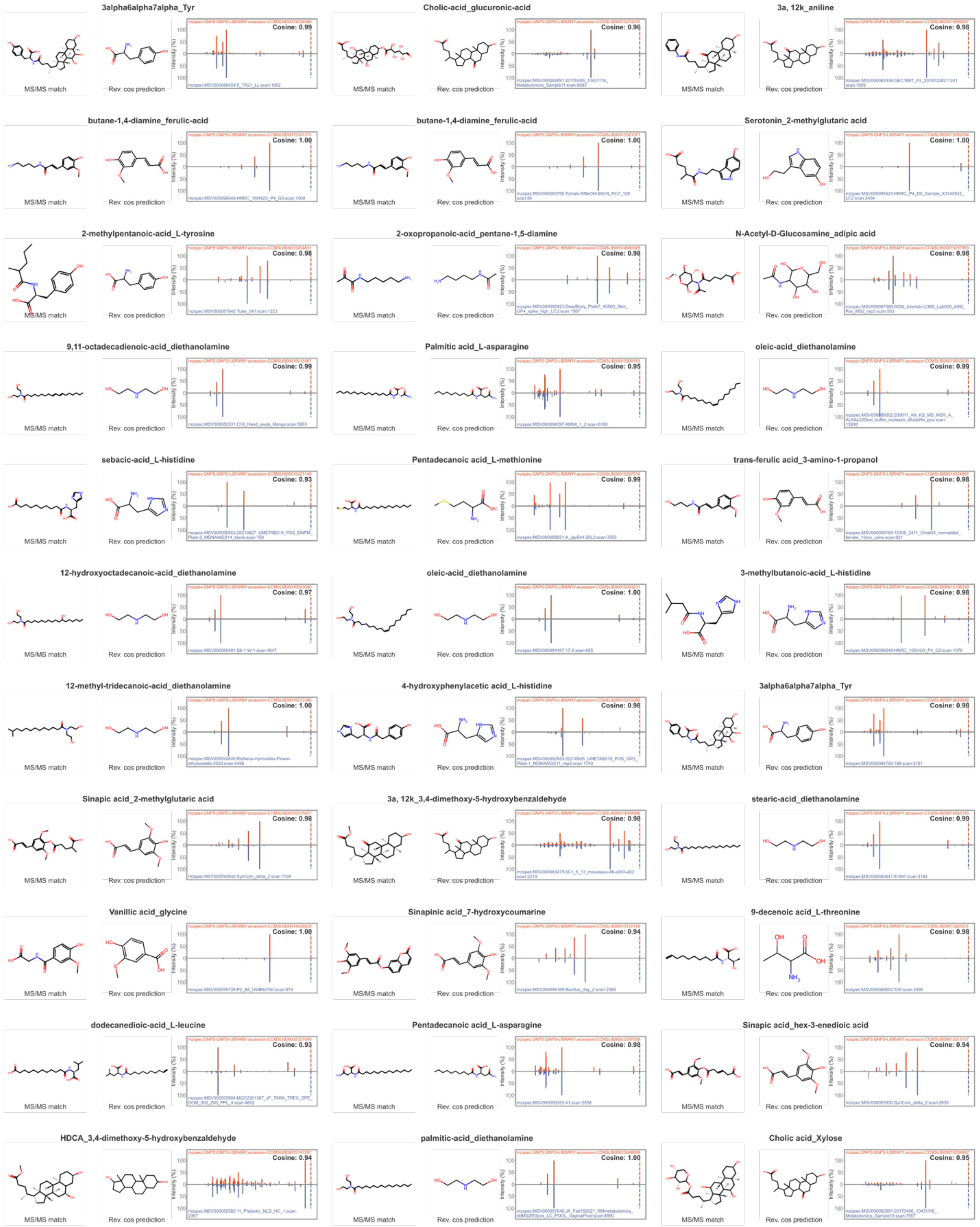
3alpha6alpha7alpha_Tyr
MS/MS match
Rev. cos prediction
Cosine: 0.99
Cholic-acid_glucuronic-acid
MS/MS match
Rev. cos prediction
Cosine: 0.96
3a, 12k_aniline
MS/MS match
Rev. cos prediction
Cosine: 0.98
butane-1,4-diamine_ferulic-acid
MS/MS match
Rev. cos prediction
Cosine: 1.00
butane-1,4-diamine_ferulic-acid
MS/MS match
Rev. cos prediction
Cosine: 1.00
Serotonin_2-methylglutaric acid
MS/MS match
Rev. cos prediction
Cosine: 1.00
2-methylpentanoic-acid_L-tyrosine
MS/MS match
Rev. cos prediction
Cosine: 0.98
2-oxopropanoic-acid_pentane-1,5-diamine
MS/MS match
Rev. cos prediction
Cosine: 0.98
N-Acetyl-D-Glucosamine_adipic acid
MS/MS match
Rev. cos prediction
Cosine: 0.98
9,11-octadecadienoic-acid_diethanolamine
MS/MS match
Rev. cos prediction
Cosine: 0.99
Palmitic acid_L-asparagine
MS/MS match
Rev. cos prediction
Cosine: 0.95
oleic-acid_diethanolamine
MS/MS match
Rev. cos prediction
Cosine: 0.98
sebacic-acid_L-histidine
MS/MS match
Rev. cos prediction
Cosine: 0.93
Pentadecanoic acid_L-methionine
MS/MS match
Rev. cos prediction
Cosine: 0.99
trans-ferulic acid_3-amino-1-propanol
MS/MS match
Rev. cos prediction
Cosine: 0.98
12-hydroxyoctadecanoic-acid_diethanolamine
MS/MS match
Rev. cos prediction
Cosine: 0.97
oleic-acid_diethanolamine
MS/MS match
Rev. cos prediction
Cosine: 1.00
3-methylbutanoic-acid_L-histidine
MS/MS match
Rev. cos prediction
Cosine: 0.98
12-methyl-tridecanoic-acid_diethanolamine
MS/MS match
Rev. cos prediction
Cosine: 1.00
4-hydroxyphenylacetic acid_L-histidine
MS/MS match
Rev. cos prediction
Cosine: 0.98
3alpha6alpha7alpha_Tyr
MS/MS match
Rev. cos prediction
Cosine: 0.98
Sinapic acid_2-methylglutaric acid
MS/MS match
Rev. cos prediction
Cosine: 0.98
3a, 12k_3,4-dimethoxy-5-hydroxybenzaldehyde
MS/MS match
Rev. cos prediction
Cosine: 0.98
stearic-acid_diethanolamine
MS/MS match
Rev. cos prediction
Cosine: 0.99
Vanillic acid_glycine
MS/MS match
Rev. cos prediction
Cosine: 1.00
Sinapinic acid_7-hydroxycoumarine
MS/MS match
Rev. cos prediction
Cosine: 0.94
9-decenoic acid_L-threonine
MS/MS match
Rev. cos prediction
Cosine: 0.98
dodecanedioic-acid_L-leucine
MS/MS match
Rev. cos prediction
Cosine: 0.93
Pentadecanoic acid_L-asparagine
MS/MS match
Rev. cos prediction
Cosine: 0.98
Sinapic acid_hex-3-enedioic acid
MS/MS match
Rev. cos prediction
Cosine: 0.94
HDCA_3,4-dimethoxy-5-hydroxybenzaldehyde
MS/MS match
Rev. cos prediction
Cosine: 0.94
palmitic-acid_diethanolamine
MS/MS match
Rev. cos prediction
Cosine: 1.00
Cholic acid_Xylose
MS/MS match
Rev. cos prediction
Cosine: 0.95

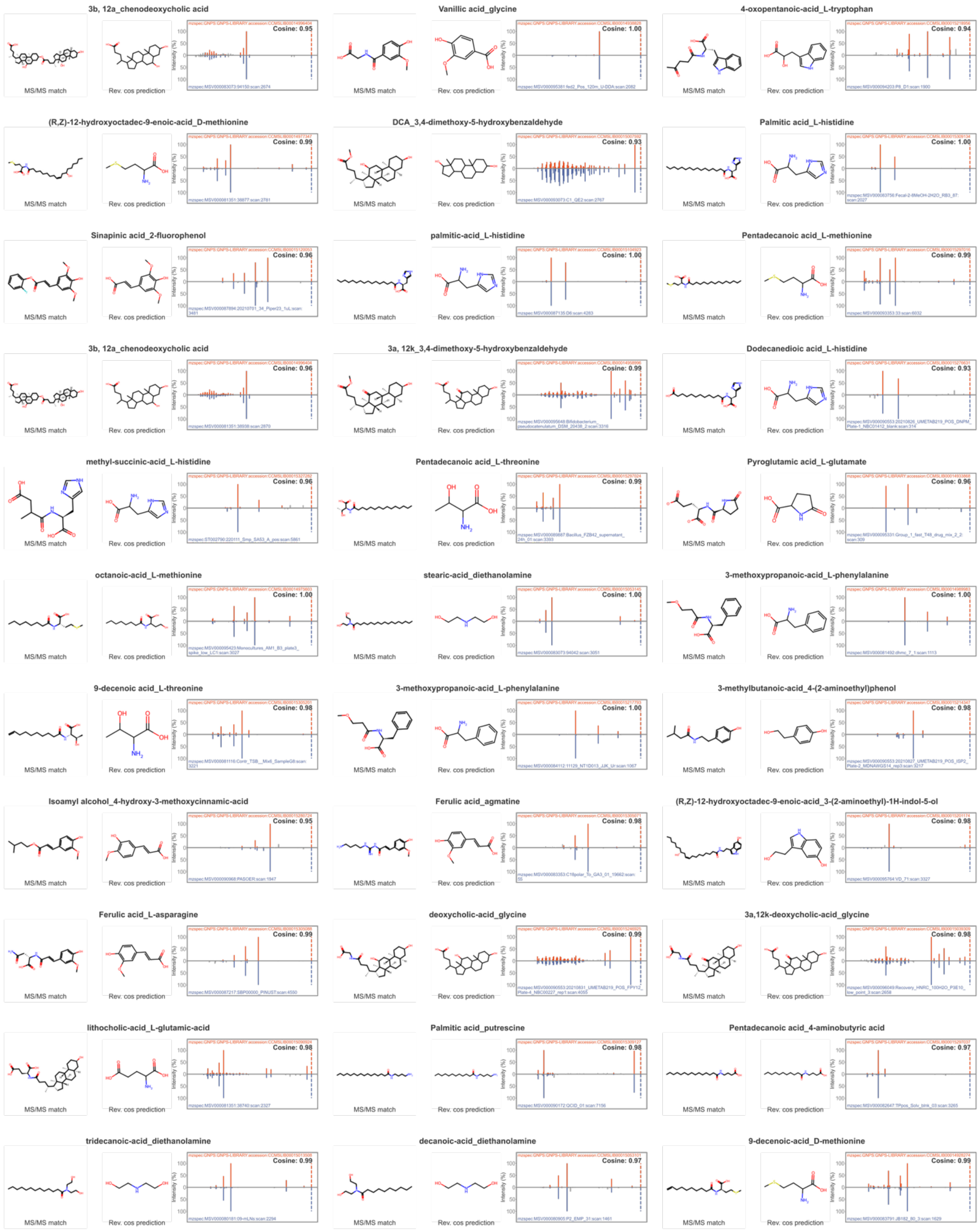
3b, 12a_chenodeoxycholic acid
MS/MS match
Rev. cos prediction
Cosine: 0.95
Vanillic acid_glycine
Cosine: 1.00
4-oxopentanoic-acid_L-tryptophan
Cosine: 0.94
(R,Z)-12-hydroxyoctadec-9-enoic-acid_D-methionine
Cosine: 0.99
DCA_3,4-dimethoxy-5-hydroxybenzaldehyde
Cosine: 0.93
Palmitic acid_L-histidine
Cosine: 1.00
Sinapinic acid_2-fluorophenol
Cosine: 0.96
palmitic-acid_L-histidine
Cosine: 1.00
Pentadecanoic acid_L-methionine
Cosine: 0.99
3b, 12a_chenodeoxycholic acid
Cosine: 0.96
3a, 12k_3,4-dimethoxy-5-hydroxybenzaldehyde
Cosine: 0.99
Dodecanedioic acid_L-histidine
Cosine: 0.93
methyl-succinic-acid_L-histidine
Cosine: 0.96
Pentadecanoic acid_L-threonine
Cosine: 0.99
Pyroglutamic acid_L-glutamate
Cosine: 0.96
octanoic-acid_L-methionine
Cosine: 1.00
stearic-acid_diethanolamine
Cosine: 1.00
3-methoxypropanoic-acid_L-phenylalanine
Cosine: 1.00
9-decenoic acid_L-threonine
Cosine: 0.98
3-methoxypropanoic-acid_L-phenylalanine
Cosine: 1.00
3-methylbutanoic-acid_4-(2-aminoethyl)phenol
Cosine: 0.98
Isoamyl alcohol_4-hydroxy-3-methoxycinnamic-acid
Cosine: 0.95
Ferulic acid_agmatine
Cosine: 0.98
(R,Z)-12-hydroxyoctadec-9-enoic-acid_3-(2-aminoethyl)-1H-indol-5-ol
Cosine: 0.98
Ferulic acid_L-asparagine
Cosine: 0.99
deoxycholic-acid_glycine
Cosine: 0.99
3a,12k-deoxycholic-acid_glycine
Cosine: 0.98
lithocholic-acid_L-glutamic-acid
Cosine: 0.98
Palmitic acid_putrescine
Cosine: 0.98
Pentadecanoic acid_4-aminobutyric acid
Cosine: 0.97
tridecanoic-acid_diethanolamine
Cosine: 0.99
decanoic-acid_diethanolamine
Cosine: 0.97
9-decenoic-acid_D-methionine
Cosine: 0.99

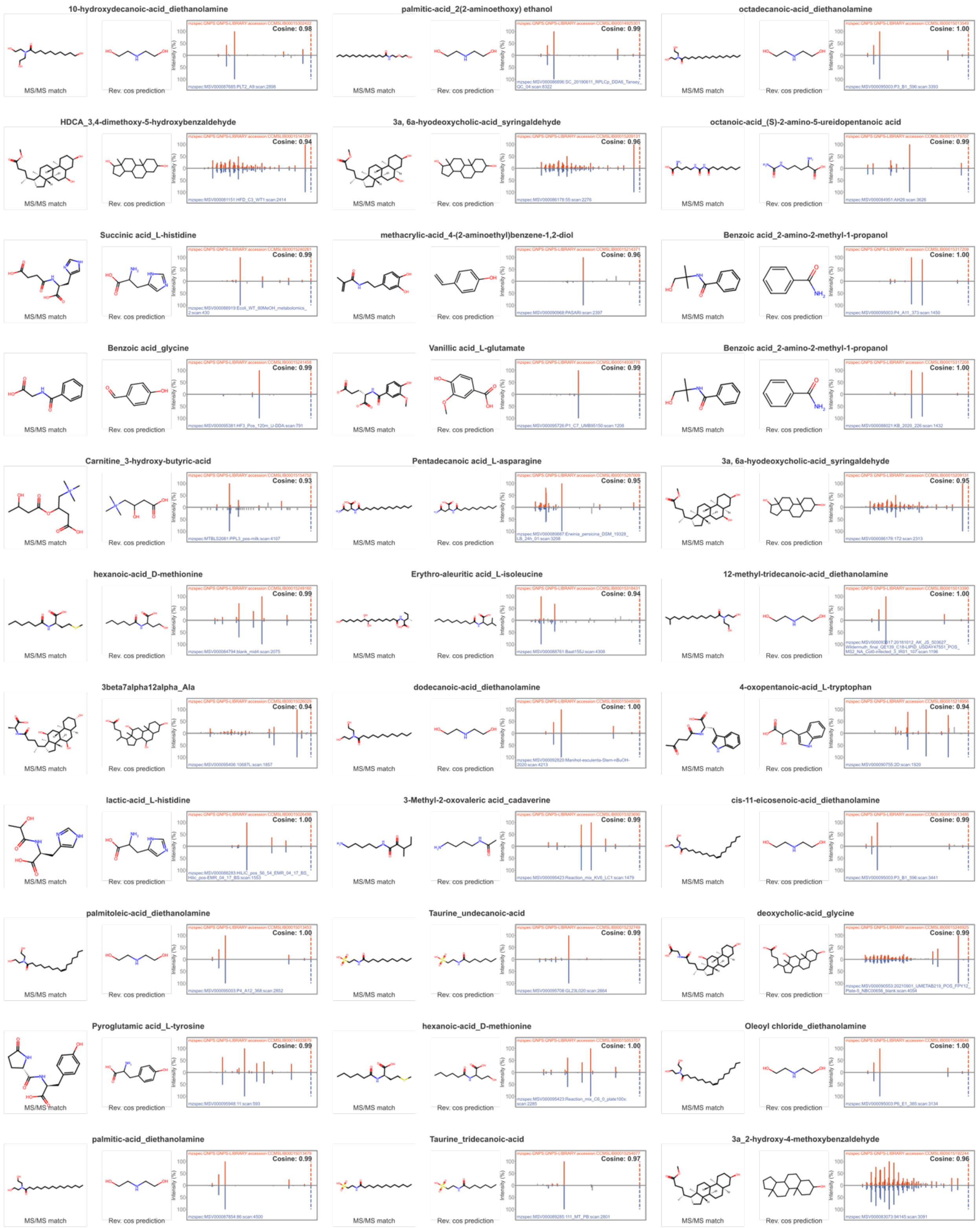
10-hydroxydecanoic-acid_diethanolamine
MS/MS match
Rev. cos prediction
Cosine: 0.98
palmitic-acid_2(2-aminoethoxy) ethanol
MS/MS match
Rev. cos prediction
Cosine: 0.99
octadecanoic-acid_diethanolamine
MS/MS match
Rev. cos prediction
Cosine: 1.00
HDCA_3,4-dimethoxy-5-hydroxybenzaldehyde
MS/MS match
Rev. cos prediction
Cosine: 0.94
3a, 6a-hyodeoxycholic-acid_syringaldehyde
MS/MS match
Rev. cos prediction
Cosine: 0.96
octanoic-acid_(S)-2-amino-5-ureidopentanoic acid
MS/MS match
Rev. cos prediction
Cosine: 0.99
Succinic acid_L-histidine
MS/MS match
Rev. cos prediction
Cosine: 0.99
methacrylic-acid_4-(2-aminoethyl)benzene-1,2-diol
MS/MS match
Rev. cos prediction
Cosine: 0.96
Benzoic acid_2-amino-2-methyl-1-propanol
MS/MS match
Rev. cos prediction
Cosine: 1.00
Benzoic acid_glycine
MS/MS match
Rev. cos prediction
Cosine: 0.99
Vanillic acid_L-glutamate
MS/MS match
Rev. cos prediction
Cosine: 0.99
Benzoic acid_2-amino-2-methyl-1-propanol
MS/MS match
Rev. cos prediction
Cosine: 1.00
Carnitine_3-hydroxy-butyric-acid
MS/MS match
Rev. cos prediction
Cosine: 0.93
Pentadecanoic acid_L-asparagine
MS/MS match
Rev. cos prediction
Cosine: 0.95
3a, 6a-hyodeoxycholic-acid_syringaldehyde
MS/MS match
Rev. cos prediction
Cosine: 0.95
hexanoic-acid_D-methionine
MS/MS match
Rev. cos prediction
Cosine: 0.99
Erythro-aleuritic acid_L-isoleucine
MS/MS match
Rev. cos prediction
Cosine: 0.94
12-methyl-tridecanoic-acid_diethanolamine
MS/MS match
Rev. cos prediction
Cosine: 1.00
3beta7alpha12alpha_Ala
MS/MS match
Rev. cos prediction
Cosine: 0.94
dodecanoic-acid_diethanolamine
MS/MS match
Rev. cos prediction
Cosine: 1.00
4-oxopentanoic-acid_L-tryptophan
MS/MS match
Rev. cos prediction
Cosine: 0.94
lactic-acid_L-histidine
MS/MS match
Rev. cos prediction
Cosine: 1.00
3-Methyl-2-oxovaleric acid_cadaverine
MS/MS match
Rev. cos prediction
Cosine: 0.99
cis-11-eicosenoic-acid_diethanolamine
MS/MS match
Rev. cos prediction
Cosine: 0.99
palmitoleic-acid_diethanolamine
MS/MS match
Rev. cos prediction
Cosine: 1.00
Taurine_undecanoic-acid
MS/MS match
Rev. cos prediction
Cosine: 0.99
deoxycholic-acid_glycine
MS/MS match
Rev. cos prediction
Cosine: 0.99
Pyroglutamic acid_L-tyrosine
MS/MS match
Rev. cos prediction
Cosine: 0.99
hexanoic-acid_D-methionine
MS/MS match
Rev. cos prediction
Cosine: 1.00
Oleoyl chloride_diethanolamine
MS/MS match
Rev. cos prediction
Cosine: 1.00
palmitic-acid_diethanolamine
MS/MS match
Rev. cos prediction
Cosine: 0.99
Taurine_tridecanoic-acid
MS/MS match
Rev. cos prediction
Cosine: 0.97
3a_2-hydroxy-4-methoxybenzaldehyde
MS/MS match
Rev. cos prediction
Cosine: 0.96

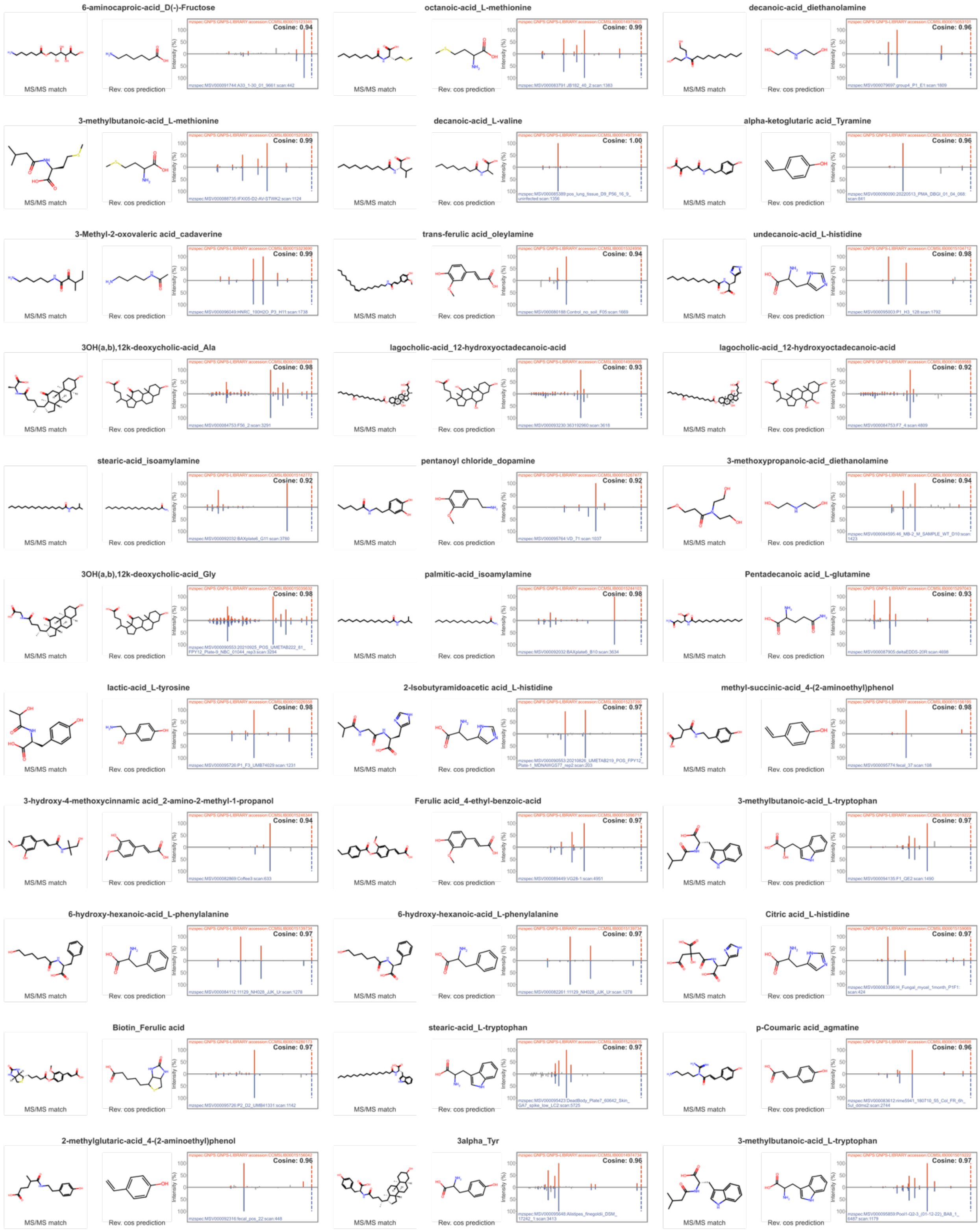
6-aminocaproic-acid_D(-)-Fructose
Cosine: 0.94
octanoic-acid_L-methionine
Cosine: 0.99
decanoic-acid_diethanolamine
Cosine: 0.96
3-methylbutanoic-acid_L-methionine
Cosine: 0.99
decanoic-acid_L-valine
Cosine: 1.00
alpha-ketoglutaric acid_Tyramine
Cosine: 0.96
3-Methyl-2-oxovaleric acid_cadaverine
Cosine: 0.99
trans-ferulic acid_oleylamine
Cosine: 0.94
undecanoic-acid_L-histidine
Cosine: 0.98
3OH(a,b),12k-deoxycholic-acid_Ala
Cosine: 0.98
lagocholic-acid_12-hydroxyoctadecanoic-acid
Cosine: 0.93
lagocholic-acid_12-hydroxyoctadecanoic-acid
Cosine: 0.92
stearic-acid_isoamylamine
Cosine: 0.92
pentanoyl chloride_dopamine
Cosine: 0.92
3-methoxypropanoic-acid_diethanolamine
Cosine: 0.94
3OH(a,b),12k-deoxycholic-acid_Gly
Cosine: 0.98
palmitic-acid_isoamylamine
Cosine: 0.98
Pentadecanoic acid_L-glutamine
Cosine: 0.93
lactic-acid_L-tyrosine
Cosine: 0.98
2-Isobutyramidoacetic acid_L-histidine
Cosine: 0.97
methyl-succinic-acid_4-(2-aminoethyl)phenol
Cosine: 0.98
3-hydroxy-4-methoxycinnamic acid_2-amino-2-methyl-1-propanol
Cosine: 0.94
Ferulic acid_4-ethyl-benzoic-acid
Cosine: 0.97
3-methylbutanoic-acid_L-tryptophan
Cosine: 0.97
6-hydroxy-hexanoic-acid_L-phenylalanine
Cosine: 0.97
6-hydroxy-hexanoic-acid_L-phenylalanine
Cosine: 0.97
Citric acid_L-histidine
Cosine: 0.97
Biotin_Ferulic acid
Cosine: 0.97
stearic-acid_L-tryptophan
Cosine: 0.97
p-Coumaric acid_agmatine
Cosine: 0.96
2-methylglutaric-acid_4-(2-aminoethyl)phenol
Cosine: 0.96
3alpha_Tyr
Cosine: 0.96
3-methylbutanoic-acid_L-tryptophan
Cosine: 0.97
MS/MS match
Rev. cos prediction
Intensity (%)

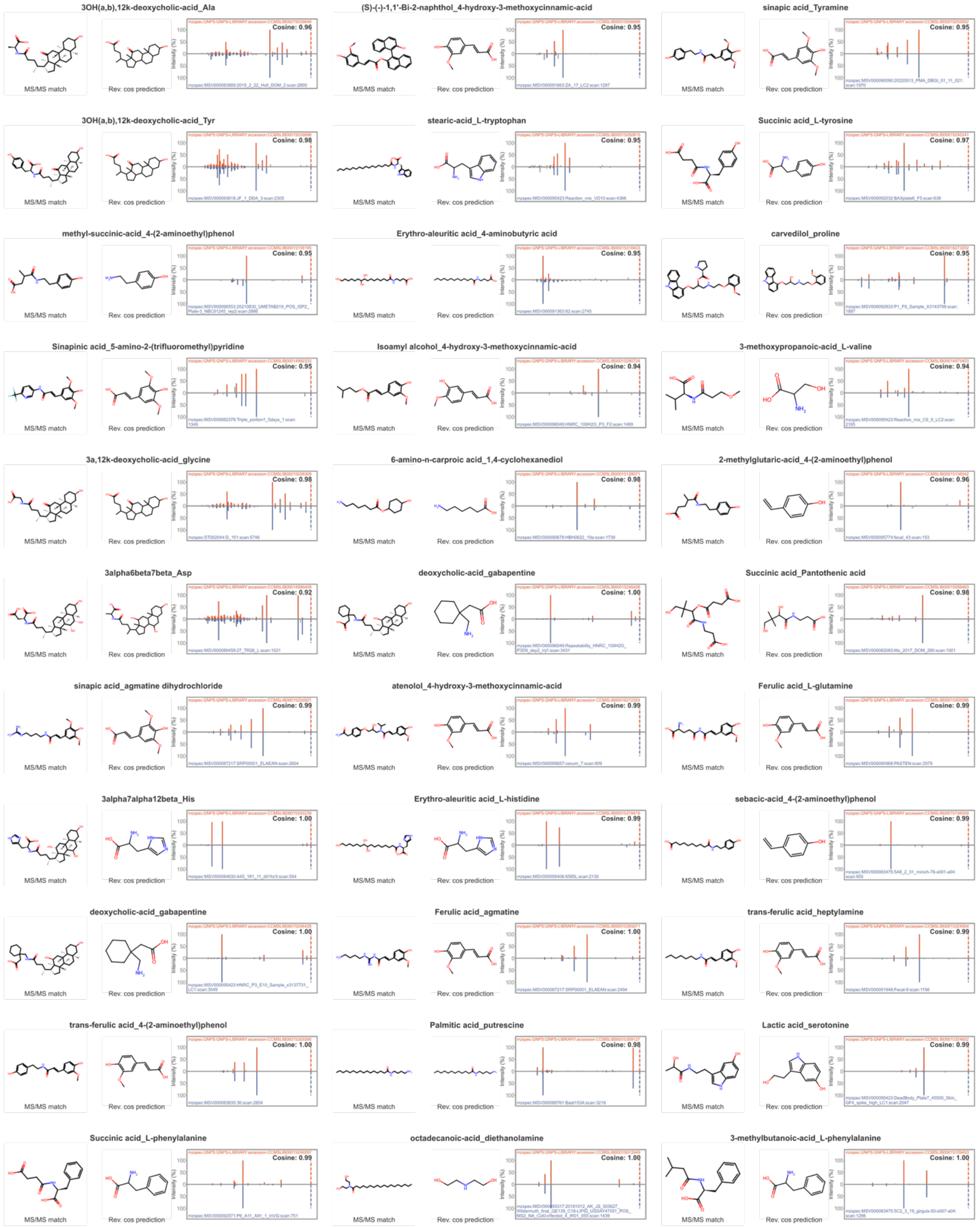
3OH(a,b),12k-deoxycholic-acid_Ala
Cosine: 0.96
(S)-(-)-1,1'-Bi-2-naphthol_4-hydroxy-3-methoxycinnamic-acid
Cosine: 0.95
sinapic acid_Tyramine
Cosine: 0.95
3OH(a,b),12k-deoxycholic-acid_Tyr
Cosine: 0.98
stearic-acid_L-tryptophan
Cosine: 0.95
Succinic acid_L-tyrosine
Cosine: 0.97
methyl-succinic-acid_4-(2-aminoethyl)phenol
Cosine: 0.95
Erythro-aleuritic acid_4-aminobutyric acid
Cosine: 0.95
carvedilol_proline
Cosine: 0.95
Sinapinic acid_5-amino-2-(trifluoromethyl)pyridine
Cosine: 0.95
Isoamyl alcohol_4-hydroxy-3-methoxycinnamic-acid
Cosine: 0.94
3-methoxypropanoic-acid_L-valine
Cosine: 0.94
3a,12k-deoxycholic-acid_glycine
Cosine: 0.98
6-amino-n-carproic acid_1,4-cyclohexanediol
Cosine: 0.98
2-methylglutaric-acid_4-(2-aminoethyl)phenol
Cosine: 0.96
3alpha6beta7beta_Asp
Cosine: 0.92
deoxycholic-acid_gabapentine
Cosine: 1.00
Succinic acid_Pantothenic acid
Cosine: 0.98
sinapic acid_agmatine dihydrochloride
Cosine: 0.99
atenolol_4-hydroxy-3-methoxycinnamic-acid
Cosine: 0.99
Ferulic acid_L-glutamine
Cosine: 0.99
3alpha7alpha12beta_His
Cosine: 1.00
Erythro-aleuritic acid_L-histidine
Cosine: 0.99
sebacic-acid_4-(2-aminoethyl)phenol
Cosine: 0.99
deoxycholic-acid_gabapentine
Cosine: 1.00
Ferulic acid_agmatine
Cosine: 1.00
trans-ferulic acid_heptylamine
Cosine: 0.99
trans-ferulic acid_4-(2-aminoethyl)phenol
Cosine: 1.00
Palmitic acid_putrescine
Cosine: 0.98
Lactic acid_serotonine
Cosine: 0.99
Succinic acid_L-phenylalanine
Cosine: 0.99
octadecanoic-acid_diethanolamine
Cosine: 1.00
3-methylbutanoic-acid_L-phenylalanine
Cosine: 1.00
MS/MS match
Rev. cos prediction

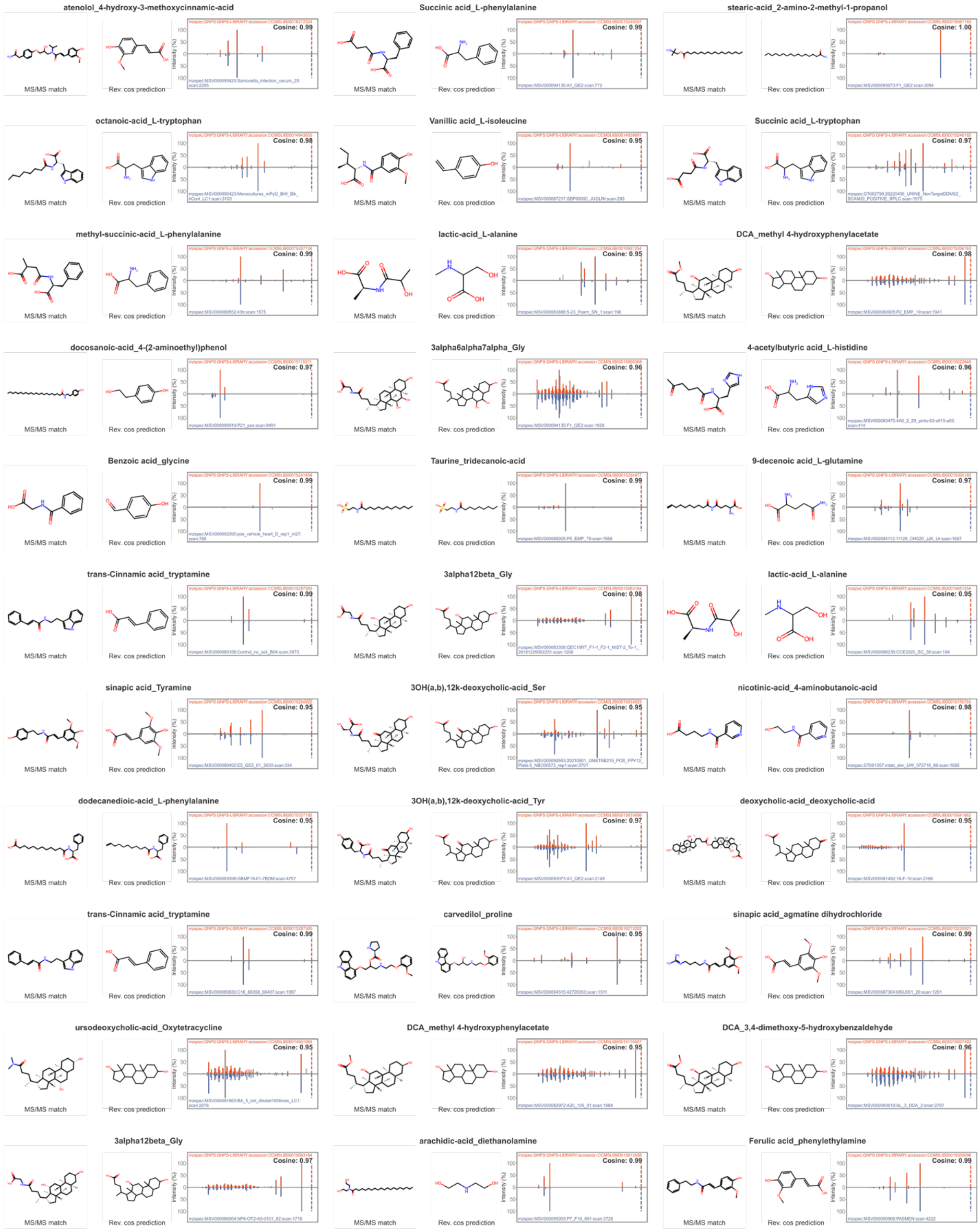
atenolol_4-hydroxy-3-methoxycinnamic-acid
Cosine: 0.99
Succinic acid_L-phenylalanine
Cosine: 0.99
stearic-acid_2-amino-2-methyl-1-propanol
Cosine: 1.00
octanoic-acid_L-tryptophan
Cosine: 0.98
Vanillic acid_L-isoleucine
Cosine: 0.95
Succinic acid_L-tryptophan
Cosine: 0.97
methyl-succinic-acid_L-phenylalanine
Cosine: 0.99
lactic-acid_L-alanine
Cosine: 0.95
DCA_methyl 4-hydroxyphenylacetate
Cosine: 0.98
docosanoic-acid_4-(2-aminoethyl)phenol
Cosine: 0.97
3alpha6alpha7alpha_Gly
Cosine: 0.96
4-acetylbutyric acid_L-histidine
Cosine: 0.96
Benzoic acid_glycine
Cosine: 0.99
Taurine_tridecanoic-acid
Cosine: 0.99
9-decenoic acid_L-glutamine
Cosine: 0.97
trans-Cinnamic acid_tryptamine
Cosine: 0.99
3alpha12beta_Gly
Cosine: 0.98
lactic-acid_L-alanine
Cosine: 0.95
sinapic acid_Tyramine
Cosine: 0.95
3OH(a,b),12k-deoxycholic-acid_Ser
Cosine: 0.95
nicotinic-acid_4-aminobutanoic-acid
Cosine: 0.98
dodecanedioic-acid_L-phenylalanine
Cosine: 0.95
3OH(a,b),12k-deoxycholic-acid_Tyr
Cosine: 0.97
deoxycholic-acid_deoxycholic-acid
Cosine: 0.95
trans-Cinnamic acid_tryptamine
Cosine: 0.99
carvedilol_proline
Cosine: 0.95
sinapic acid_agmatine dihydrochloride
Cosine: 0.99
ursodeoxycholic-acid_Oxytetracycline
Cosine: 0.95
DCA_methyl 4-hydroxyphenylacetate
Cosine: 0.95
DCA_3,4-dimethoxy-5-hydroxybenzaldehyde
Cosine: 0.96
3alpha12beta_Gly
Cosine: 0.97
arachidic-acid_diethanolamine
Cosine: 0.99
Ferulic acid_phenylethylamine
Cosine: 0.99
MS/MS match
Rev. cos prediction
Intensity (%)

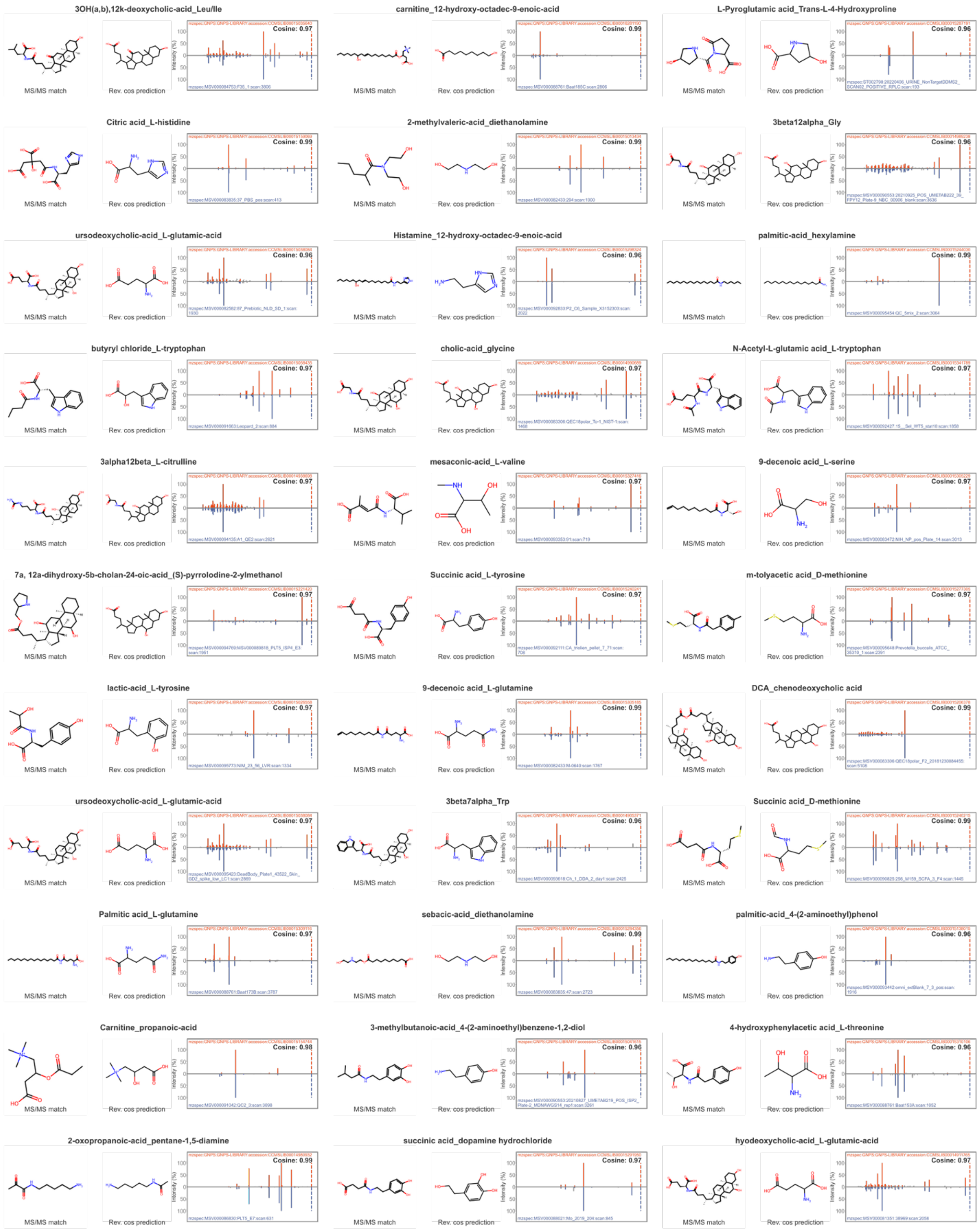
3OH(a,b),12k-deoxycholic-acid_Leu/Ile
MS/MS match
Rev. cos prediction
Cosine: 0.97
carnitine_12-hydroxy-octadec-9-enoic-acid
MS/MS match
Rev. cos prediction
Cosine: 0.99
L-Pyroglutamic acid_Trans-L-4-Hydroxyproline
MS/MS match
Rev. cos prediction
Cosine: 0.96
Citric acid_L-histidine
MS/MS match
Rev. cos prediction
Cosine: 0.99
2-methylvaleric-acid_diethanolamine
MS/MS match
Rev. cos prediction
Cosine: 0.99
3beta12alpha_Gly
MS/MS match
Rev. cos prediction
Cosine: 0.96
ursodeoxycholic-acid_L-glutamic-acid
MS/MS match
Rev. cos prediction
Cosine: 0.96
Histamine_12-hydroxy-octadec-9-enoic-acid
MS/MS match
Rev. cos prediction
Cosine: 0.96
palmitic-acid_hexylamine
MS/MS match
Rev. cos prediction
Cosine: 0.99
butyryl chloride_L-tryptophan
MS/MS match
Rev. cos prediction
Cosine: 0.97
cholic-acid_glycine
MS/MS match
Rev. cos prediction
Cosine: 0.97
N-Acetyl-L-glutamic acid_L-tryptophan
MS/MS match
Rev. cos prediction
Cosine: 0.97
3alpha12beta_L-citrulline
MS/MS match
Rev. cos prediction
Cosine: 0.97
mesaconic-acid_L-valine
MS/MS match
Rev. cos prediction
Cosine: 0.97
9-decenoic acid_L-serine
MS/MS match
Rev. cos prediction
Cosine: 0.97
7a, 12a-dihydroxy-5b-cholan-24-oic-acid_(S)-pyrrolodine-2-ylmethanol
MS/MS match
Rev. cos prediction
Cosine: 0.97
Succinic acid_L-tyrosine
MS/MS match
Rev. cos prediction
Cosine: 0.97
m-tolyacetic acid_D-methionine
MS/MS match
Rev. cos prediction
Cosine: 0.97
lactic-acid_L-tyrosine
MS/MS match
Rev. cos prediction
Cosine: 0.97
9-decenoic acid_L-glutamine
MS/MS match
Rev. cos prediction
Cosine: 0.99
DCA_chenodeoxycholic acid
MS/MS match
Rev. cos prediction
Cosine: 0.99
ursodeoxycholic-acid_L-glutamic-acid
MS/MS match
Rev. cos prediction
Cosine: 0.97
3beta7alpha_Trp
MS/MS match
Rev. cos prediction
Cosine: 0.96
Succinic acid_D-methionine
MS/MS match
Rev. cos prediction
Cosine: 0.99
Palmitic acid_L-glutamine
MS/MS match
Rev. cos prediction
Cosine: 0.97
sebacic-acid_diethanolamine
MS/MS match
Rev. cos prediction
Cosine: 0.99
palmitic-acid_4-(2-aminoethyl)phenol
MS/MS match
Rev. cos prediction
Cosine: 0.96
Carnitine_propanoic-acid
MS/MS match
Rev. cos prediction
Cosine: 0.98
3-methylbutanoic-acid_4-(2-aminoethyl)benzene-1,2-diol
MS/MS match
Rev. cos prediction
Cosine: 0.96
4-hydroxyphenylacetic acid_L-threonine
MS/MS match
Rev. cos prediction
Cosine: 0.96
2-oxopropanoic-acid_pentane-1,5-diamine
MS/MS match
Rev. cos prediction
Cosine: 0.99
succinic acid_dopamine hydrochloride
MS/MS match
Rev. cos prediction
Cosine: 0.99
hyodeoxycholic-acid_L-glutamic-acid
MS/MS match
Rev. cos prediction
Cosine: 0.97

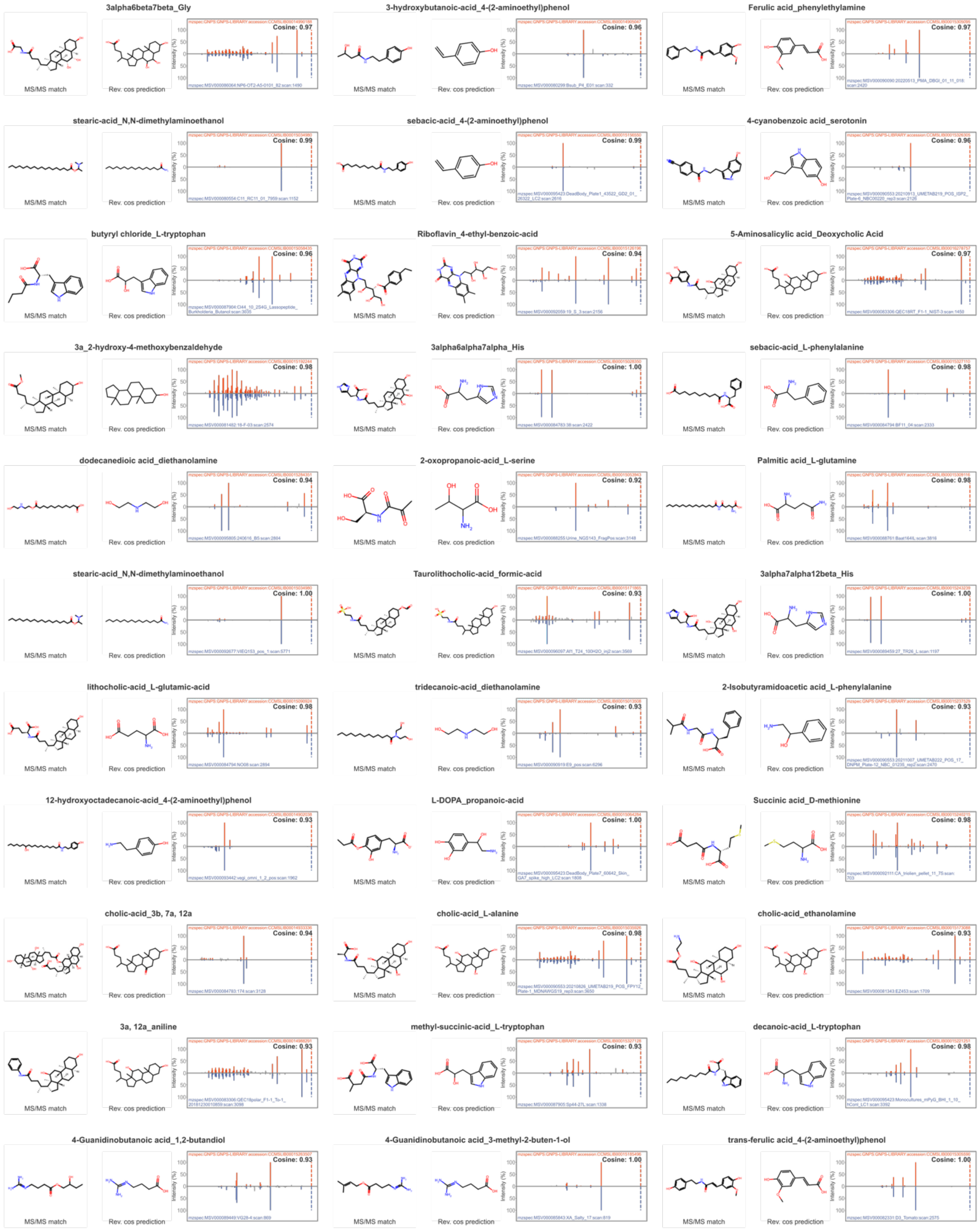
3alpha6beta7beta_Gly
Cosine: 0.97
3-hydroxybutanoic-acid_4-(2-aminoethyl)phenol
Cosine: 0.96
Ferulic acid_phenylethylamine
Cosine: 0.97
stearic-acid_N,N-dimethylaminoethanol
Cosine: 0.99
sebacic-acid_4-(2-aminoethyl)phenol
Cosine: 0.99
4-cyanobenzoic acid_serotonin
Cosine: 0.96
butyryl chloride_L-tryptophan
Cosine: 0.96
Riboflavin_4-ethyl-benzoic-acid
Cosine: 0.94
5-Aminosalicylic acid_Deoxycholic Acid
Cosine: 0.97
3a_2-hydroxy-4-methoxybenzaldehyde
Cosine: 0.98
3alpha6alpha7alpha_His
Cosine: 1.00
sebacic-acid_L-phenylalanine
Cosine: 0.98
dodecanedioic acid_diethanolamine
Cosine: 0.94
2-oxopropanoic-acid_L-serine
Cosine: 0.92
Palmitic acid_L-glutamine
Cosine: 0.98
stearic-acid_N,N-dimethylaminoethanol
Cosine: 1.00
Taurolithocholic-acid_formic-acid
Cosine: 0.93
3alpha7alpha12beta_His
Cosine: 1.00
lithocholic-acid_L-glutamic-acid
Cosine: 0.98
tridecanoic-acid_diethanolamine
Cosine: 0.93
2-Isobutyramidoacetic acid_L-phenylalanine
Cosine: 0.93
12-hydroxyoctadecanoic-acid_4-(2-aminoethyl)phenol
Cosine: 0.93
L-DOPA_propanoic-acid
Cosine: 1.00
Succinic acid_D-methionine
Cosine: 0.98
cholic-acid_3b, 7a, 12a
Cosine: 0.94
cholic-acid_L-alanine
Cosine: 0.98
cholic-acid_ethanolamine
Cosine: 0.93
3a, 12a_aniline
Cosine: 0.93
methyl-succinic-acid_L-tryptophan
Cosine: 0.93
decanoic-acid_L-tryptophan
Cosine: 0.98
4-Guanidinobutanoic acid_1,2-butandiol
Cosine: 0.93
4-Guanidinobutanoic acid_3-methyl-2-buten-1-ol
Cosine: 1.00
trans-ferulic acid_4-(2-aminoethyl)phenol
Cosine: 1.00
MS/MS match
Rev. cos prediction
Intensity (%)

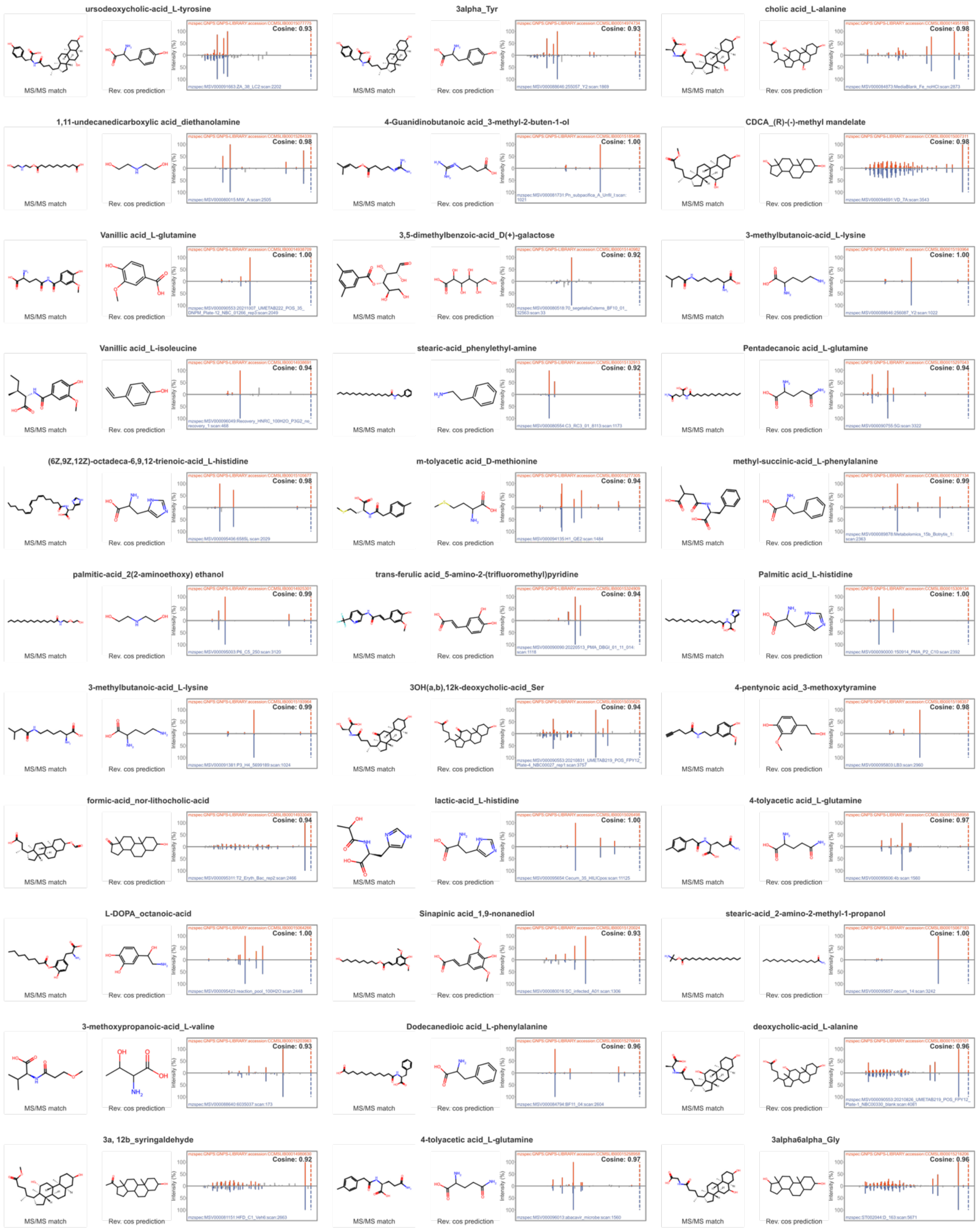
ursodeoxycholic-acid_L-tyrosine
Cosine: 0.93
MS/MS match
Rev. cos prediction
3alpha_Tyr
Cosine: 0.93
MS/MS match
Rev. cos prediction
cholic acid_L-alanine
Cosine: 0.98
MS/MS match
Rev. cos prediction
1,11-undecanedicarboxylic acid_diethanolamine
Cosine: 0.98
MS/MS match
Rev. cos prediction
4-Guanidinobutanoic acid_3-methyl-2-buten-1-ol
Cosine: 1.00
MS/MS match
Rev. cos prediction
CDCA_(R)-(-)-methyl mandelate
Cosine: 0.98
MS/MS match
Rev. cos prediction
Vanillic acid_L-glutamine
Cosine: 1.00
MS/MS match
Rev. cos prediction
3,5-dimethylbenzoic-acid_D(+)-galactose
Cosine: 0.92
MS/MS match
Rev. cos prediction
3-methylbutanoic-acid_L-lysine
Cosine: 1.00
MS/MS match
Rev. cos prediction
Vanillic acid_L-isoleucine
Cosine: 0.94
MS/MS match
Rev. cos prediction
stearic-acid_phenylethyl-amine
Cosine: 0.92
MS/MS match
Rev. cos prediction
Pentadecanoic acid_L-glutamine
Cosine: 0.94
MS/MS match
Rev. cos prediction
(6Z,9Z,12Z)-octadeca-6,9,12-trienoic-acid_L-histidine
Cosine: 0.98
MS/MS match
Rev. cos prediction
m-tolyacetic acid_D-methionine
Cosine: 0.94
MS/MS match
Rev. cos prediction
methyl-succinic-acid_L-phenylalanine
Cosine: 0.99
MS/MS match
Rev. cos prediction
palmitic-acid_2(2-aminoethoxy) ethanol
Cosine: 0.99
MS/MS match
Rev. cos prediction
trans-ferulic acid_5-amino-2-(trifluoromethyl)pyridine
Cosine: 0.94
MS/MS match
Rev. cos prediction
Palmitic acid_L-histidine
Cosine: 1.00
MS/MS match
Rev. cos prediction
3-methylbutanoic-acid_L-lysine
Cosine: 0.99
MS/MS match
Rev. cos prediction
3OH(a,b),12k-deoxycholic-acid_Ser
Cosine: 0.94
MS/MS match
Rev. cos prediction
4-pentynoic acid_3-methoxytyramine
Cosine: 0.98
MS/MS match
Rev. cos prediction
formic-acid_nor-lithocholic-acid
Cosine: 0.94
MS/MS match
Rev. cos prediction
lactic-acid_L-histidine
Cosine: 1.00
MS/MS match
Rev. cos prediction
4-tolyacetic acid_L-glutamine
Cosine: 0.97
MS/MS match
Rev. cos prediction
L-DOPA_octanoic-acid
Cosine: 1.00
MS/MS match
Rev. cos prediction
Sinapinic acid_1,9-nonanediol
Cosine: 0.93
MS/MS match
Rev. cos prediction
stearic-acid_2-amino-2-methyl-1-propanol
Cosine: 1.00
MS/MS match
Rev. cos prediction
3-methoxypropanoic-acid_L-valine
Cosine: 0.93
MS/MS match
Rev. cos prediction
Dodecanedioic acid_L-phenylalanine
Cosine: 0.96
MS/MS match
Rev. cos prediction
deoxycholic-acid_L-alanine
Cosine: 0.96
MS/MS match
Rev. cos prediction
3a, 12b_syringaldehyde
Cosine: 0.92
MS/MS match
Rev. cos prediction
4-tolyacetic acid_L-glutamine
Cosine: 0.97
MS/MS match
Rev. cos prediction
3alpha6alpha_Gly
Cosine: 0.96
MS/MS match
Rev. cos prediction

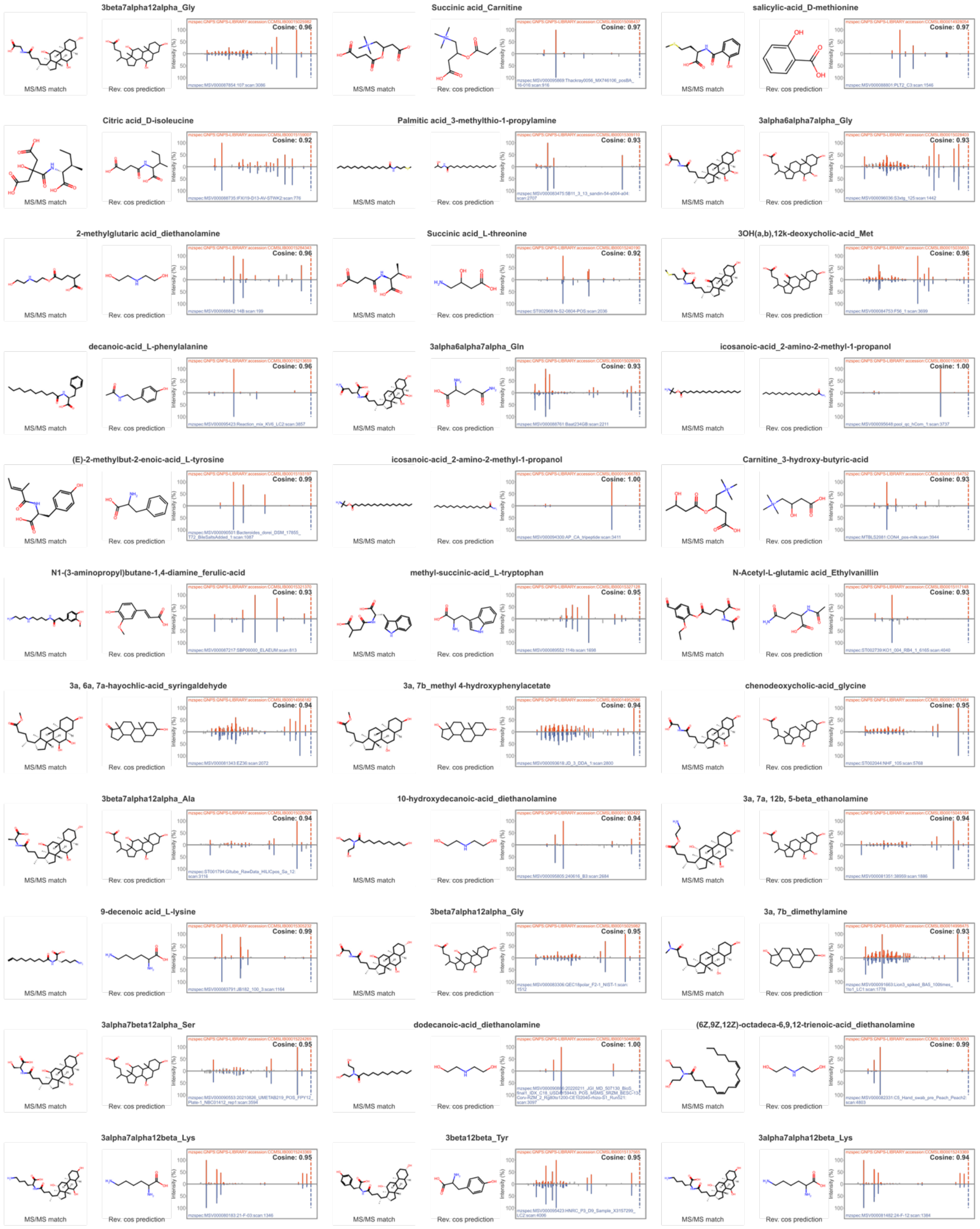
3beta7alpha12alpha_Gly
MS/MS match
Rev. cos prediction
Cosine: 0.96
Succinic acid_Carnitine
MS/MS match
Rev. cos prediction
Cosine: 0.97
salicylic-acid_D-methionine
MS/MS match
Rev. cos prediction
Cosine: 0.97
Citric acid_D-isoleucine
MS/MS match
Rev. cos prediction
Cosine: 0.92
Palmitic acid_3-methylthio-1-propylamine
MS/MS match
Rev. cos prediction
Cosine: 0.93
3alpha6alpha7alpha_Gly
MS/MS match
Rev. cos prediction
Cosine: 0.93
2-methylglutaric acid_diethanolamine
MS/MS match
Rev. cos prediction
Cosine: 0.96
Succinic acid_L-threonine
MS/MS match
Rev. cos prediction
Cosine: 0.92
3OH(a,b),12k-deoxycholic-acid_Met
MS/MS match
Rev. cos prediction
Cosine: 0.96
decanoic-acid_L-phenylalanine
MS/MS match
Rev. cos prediction
Cosine: 0.96
3alpha6alpha7alpha_Gln
MS/MS match
Rev. cos prediction
Cosine: 0.93
icosanoic-acid_2-amino-2-methyl-1-propanol
MS/MS match
Rev. cos prediction
Cosine: 1.00
(E)-2-methylbut-2-enoic-acid_L-tyrosine
MS/MS match
Rev. cos prediction
Cosine: 0.99
icosanoic-acid_2-amino-2-methyl-1-propanol
MS/MS match
Rev. cos prediction
Cosine: 1.00
Carnitine_3-hydroxy-butyric-acid
MS/MS match
Rev. cos prediction
Cosine: 0.93
N1-(3-aminopropyl)butane-1,4-diamine_ferulic-acid
MS/MS match
Rev. cos prediction
Cosine: 0.93
methyl-succinic-acid_L-tryptophan
MS/MS match
Rev. cos prediction
Cosine: 0.93
N-Acetyl-L-glutamic acid_Ethylvanillin
MS/MS match
Rev. cos prediction
Cosine: 0.93
3a, 6a, 7a-hayochlic-acid_syringaldehyde
MS/MS match
Rev. cos prediction
Cosine: 0.94
3a, 7b_methyl 4-hydroxyphenylacetate
MS/MS match
Rev. cos prediction
Cosine: 0.94
chenodeoxycholic-acid_glycine
MS/MS match
Rev. cos prediction
Cosine: 0.95
3beta7alpha12alpha_Ala
MS/MS match
Rev. cos prediction
Cosine: 0.94
10-hydroxydecanoic-acid_diethanolamine
MS/MS match
Rev. cos prediction
Cosine: 0.94
3a, 7a, 12b, 5-beta_ethanolamine
MS/MS match
Rev. cos prediction
Cosine: 0.94
9-decenoic acid_L-lysine
MS/MS match
Rev. cos prediction
Cosine: 0.99
3beta7alpha12alpha_Gly
MS/MS match
Rev. cos prediction
Cosine: 0.99
3a, 7b_dimethylamine
MS/MS match
Rev. cos prediction
Cosine: 0.93
3alpha7beta12alpha_Ser
MS/MS match
Rev. cos prediction
Cosine: 0.95
dodecanoic-acid_diethanolamine
MS/MS match
Rev. cos prediction
Cosine: 1.00
(6Z,9Z,12Z)-octadeca-6,9,12-trienoic-acid_diethanolamine
MS/MS match
Rev. cos prediction
Cosine: 0.99
3alpha7alpha12beta_Lys
MS/MS match
Rev. cos prediction
Cosine: 0.95
3beta12beta_Tyr
MS/MS match
Rev. cos prediction
Cosine: 0.95
3alpha7alpha12beta_Lys
MS/MS match
Rev. cos prediction
Cosine: 0.94

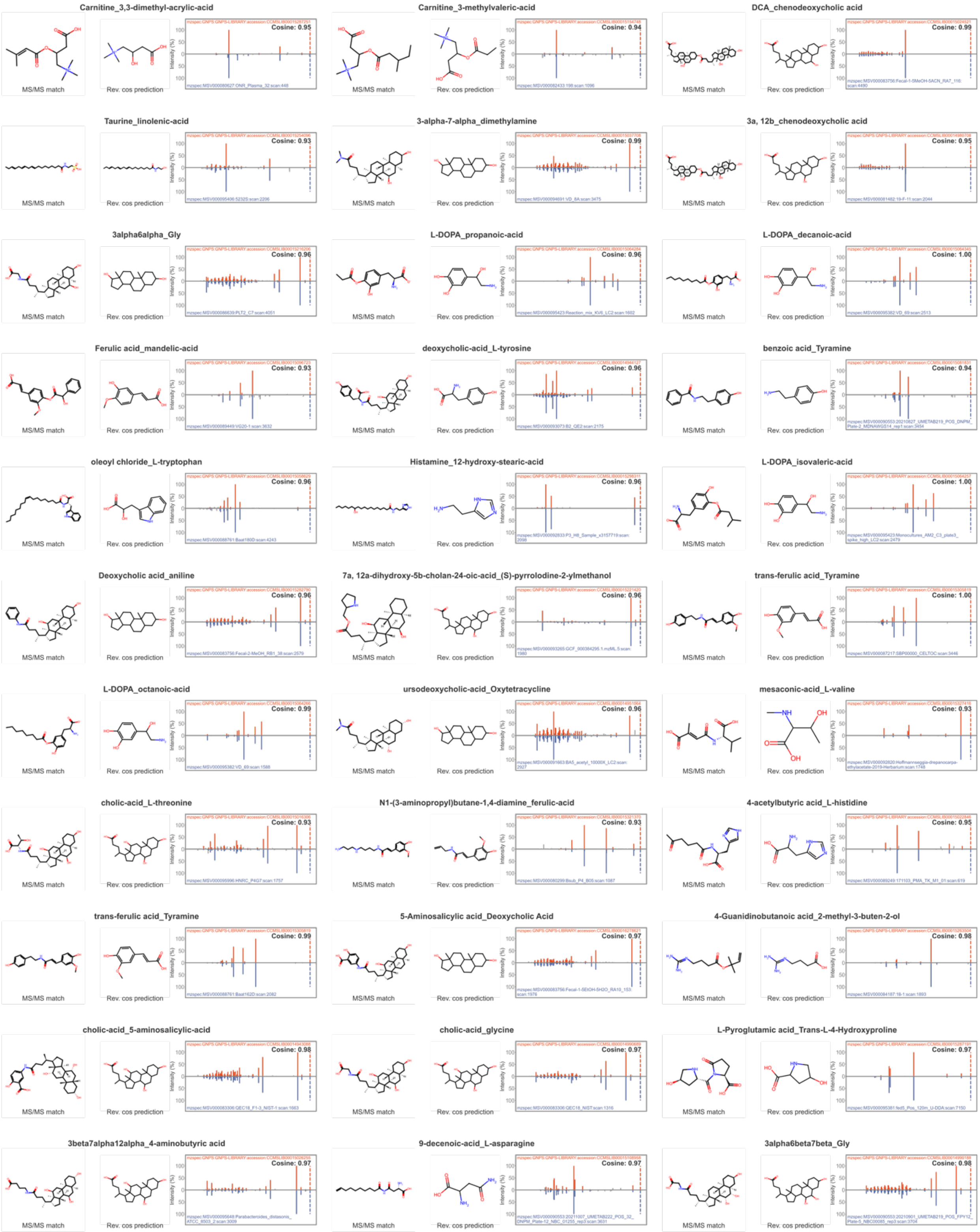

Carnitine_3,3-dimethyl-acrylic-acid
MS/MS match
Rev. cos prediction
Cosine: 0.95
Carnitine_3-methylvaleric-acid
MS/MS match
Rev. cos prediction
Cosine: 0.94
DCA_chenodeoxycholic acid
MS/MS match
Rev. cos prediction
Cosine: 0.99
Taurine_linolenic-acid
MS/MS match
Rev. cos prediction
Cosine: 0.93
3-alpha-7-alpha_dimethylamine
MS/MS match
Rev. cos prediction
Cosine: 0.99
3a, 12b_chenodeoxycholic acid
MS/MS match
Rev. cos prediction
Cosine: 0.95
3alpha6alpha_Gly
MS/MS match
Rev. cos prediction
Cosine: 0.96
L-DOPA_propanoic-acid
MS/MS match
Rev. cos prediction
Cosine: 0.96
L-DOPA_decanoic-acid
MS/MS match
Rev. cos prediction
Cosine: 1.00
Ferulic acid_mandelic-acid
MS/MS match
Rev. cos prediction
Cosine: 0.93
deoxycholic-acid_L-tyrosine
MS/MS match
Rev. cos prediction
Cosine: 0.96
benzoic acid_Tyramine
MS/MS match
Rev. cos prediction
Cosine: 0.94
oleoyl chloride_L-tryptophan
MS/MS match
Rev. cos prediction
Cosine: 0.96
Histamine_12-hydroxy-stearic-acid
MS/MS match
Rev. cos prediction
Cosine: 0.96
L-DOPA_isovaleric-acid
MS/MS match
Rev. cos prediction
Cosine: 1.00
Deoxycholic acid_aniline
MS/MS match
Rev. cos prediction
Cosine: 0.96
7a, 12a-dihydroxy-5b-cholan-24-oic-acid_(S)-pyrrolodine-2-ylmethanol
MS/MS match
Rev. cos prediction
Cosine: 0.96
trans-ferulic acid_Tyramine
MS/MS match
Rev. cos prediction
Cosine: 1.00
L-DOPA_octanoic-acid
MS/MS match
Rev. cos prediction
Cosine: 0.99
ursodeoxycholic-acid_Oxytetracycline
MS/MS match
Rev. cos prediction
Cosine: 0.96
mesaconic-acid_L-valine
MS/MS match
Rev. cos prediction
Cosine: 0.93
cholic-acid_L-threonine
MS/MS match
Rev. cos prediction
Cosine: 0.93
N1-(3-aminopropyl)butane-1,4-diamine_ferulic-acid
MS/MS match
Rev. cos prediction
Cosine: 0.93
4-acetylbutyric acid_L-histidine
MS/MS match
Rev. cos prediction
Cosine: 0.95
trans-ferulic acid_Tyramine
MS/MS match
Rev. cos prediction
Cosine: 0.99
5-Aminosalicylic acid_Deoxycholic Acid
MS/MS match
Rev. cos prediction
Cosine: 0.97
4-Guanidinobutanoic acid_2-methyl-3-buten-2-ol
MS/MS match
Rev. cos prediction
Cosine: 0.98
cholic-acid_5-aminosalicylic-acid
MS/MS match
Rev. cos prediction
Cosine: 0.98
cholic-acid_glycine
MS/MS match
Rev. cos prediction
Cosine: 0.97
L-Pyroglutamic acid_Trans-L-4-Hydroxyproline
MS/MS match
Rev. cos prediction
Cosine: 0.97
3beta7alpha12alpha_4-aminobutyric acid
MS/MS match
Rev. cos prediction
Cosine: 0.97
9-decenoic-acid_L-asparagine
MS/MS match
Rev. cos prediction
Cosine: 0.97
3alpha6beta7beta_Gly
MS/MS match
Rev. cos prediction
Cosine: 0.98

3alpha7alpha12beta_Gln

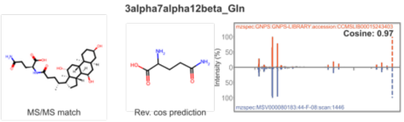


octanoic-acid_L-leucine

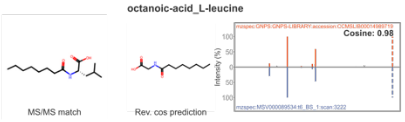


2-Methylbutylamine_eicosanoic-acid

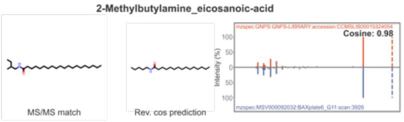


9-decenoic acid_L-asparagine

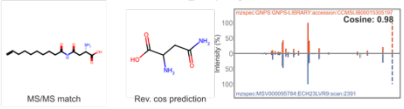


3alpha6alpha7alpha_His

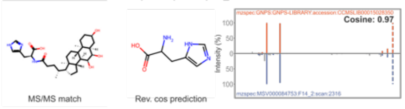


4-Guanidinobutanoic acid_farnesol

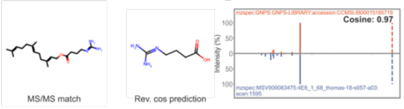


Carnitine_10-undecenoic-acid

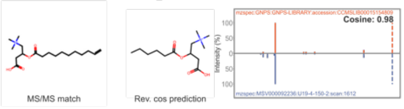


1-(4-aminobutyl)guanidine_caffeic-acid

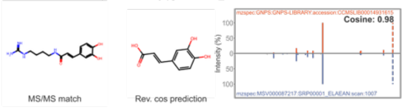


2-methylglutaric acid_diethanolamine

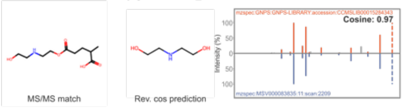


icosanoic-acid_4-(2-aminoethyl)phenol

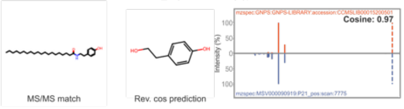


1,3-diphenylguanidine_3-methylvaleric-acid

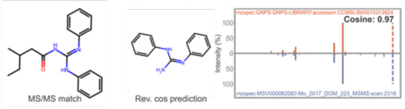


salicylic-acid_D-methionine

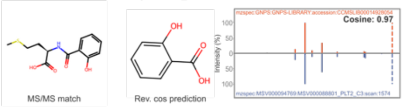


chenodeoxycholic-acid_5-aminosalicylic-acid

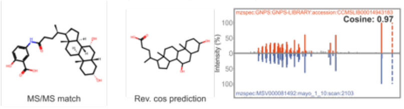


chollic-acid_Tyramine

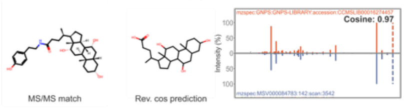


nonadecanoic-acid_N,N-dimethyl-4,4-azo dianiline

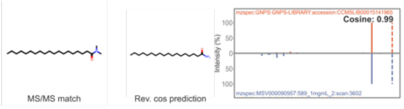


2-(4-isobutylphenyl)propionic-acid_D-methionine

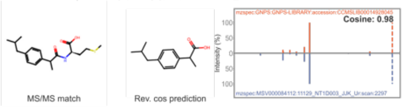


n-Cholyl_Alanine

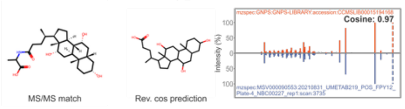


chenodeoxycholic-acid_L-glutamic-acid

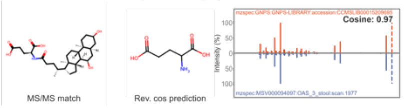


3a, 12b_chenodeoxycholic acid

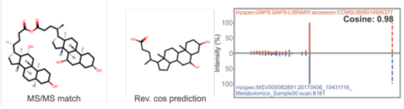


3a, 7k, 12k_(R)-(-)-methyl mandelate

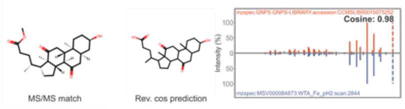


3-alpha-6-alpha-hyodeoxycholic-acid_taurine

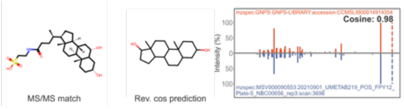


2-(4-isobutylphenyl)propionic-acid_D-methionine

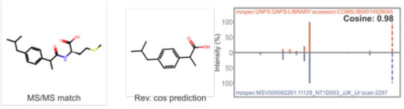


1,11-undecanedicarboxylic acid_diethanolamine

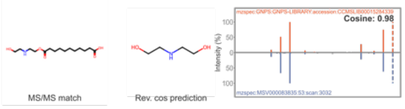


3OH(a,b),12k-deoxycholic-acid_Leu/Ile

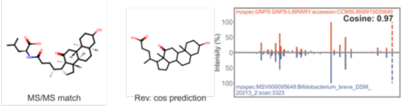


formic-acid_7b-lithocholic-acid

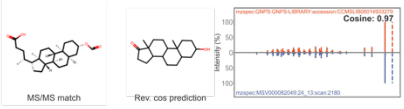


deoxycholic-acid_L-glutamic-acid

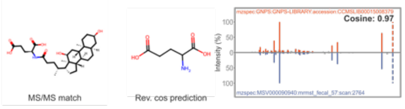


oleoyl chloride_L-tryptophan

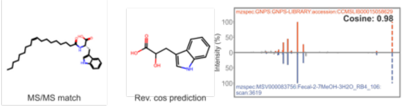


12-hydroxyoctadecanoic-acid_pentane-1,5-diamine

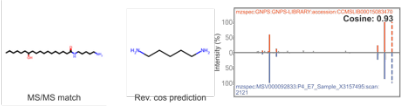


3alpha6beta7beta_4-aminobutyric acid

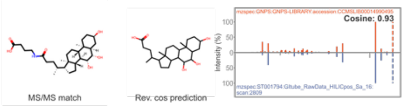


Acetyl-L-Carnitine_1,11-undecanedicarboxylic acid

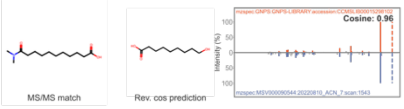


3alpha6alpha_Tyr

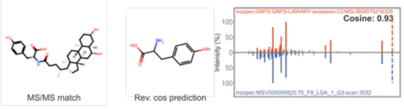


Succinic acid_L-tryptophan

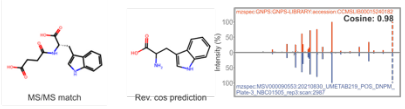


3alpha6beta7beta_Ser

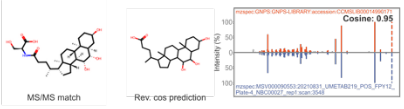

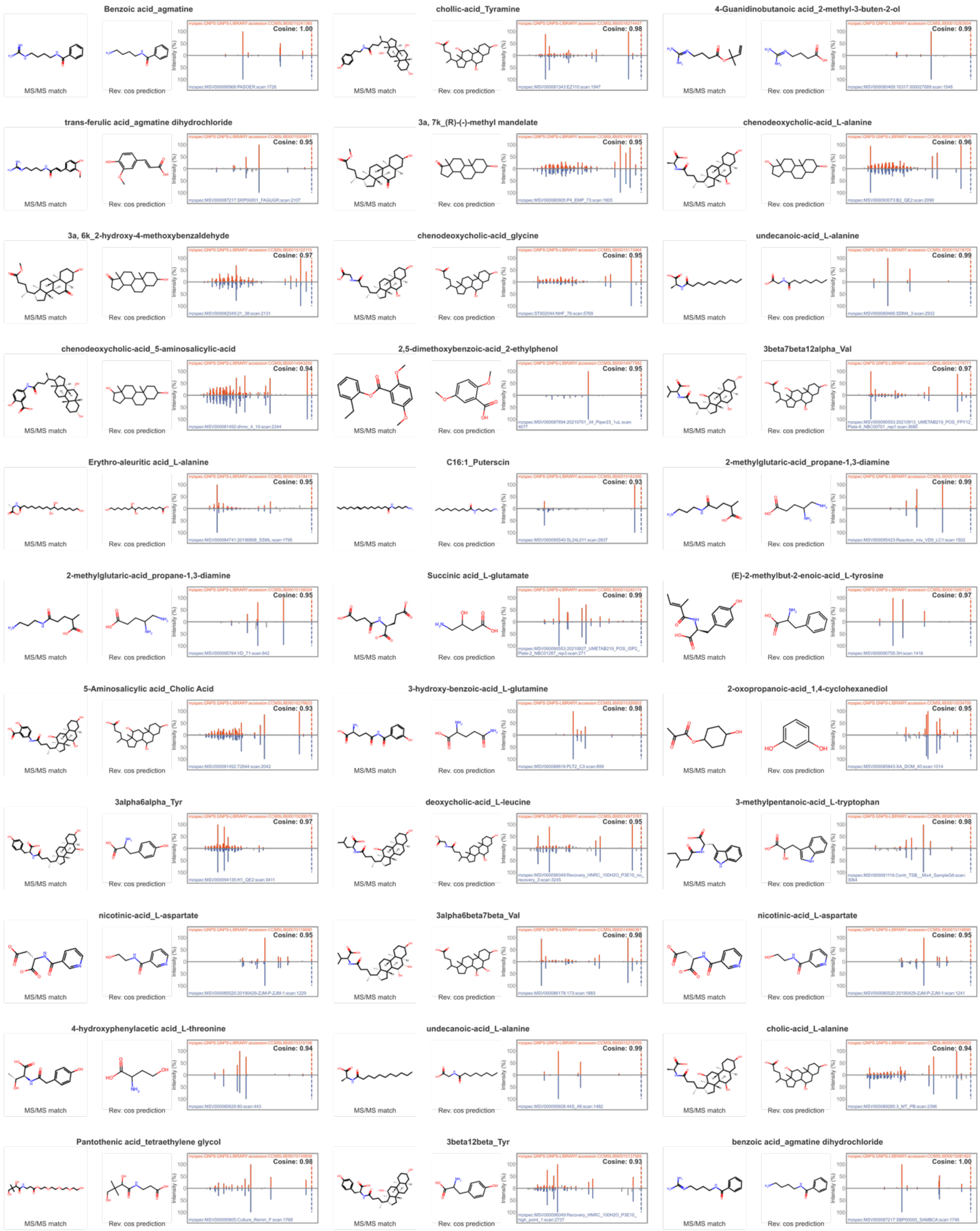
Benzoic acid_agmatine
MS/MS match
Rev. cos prediction
Cosine: 1.00
chollic-acid_Tyramine
MS/MS match
Rev. cos prediction
Cosine: 0.98
4-Guanidinobutanoic acid_2-methyl-3-buten-2-ol
MS/MS match
Rev. cos prediction
Cosine: 0.99
trans-ferulic acid_agmatine dihydrochloride
MS/MS match
Rev. cos prediction
Cosine: 0.95
3a, 7k_(R)-(-)-methyl mandelate
MS/MS match
Rev. cos prediction
Cosine: 0.95
chenodeoxycholic-acid_L-alanine
MS/MS match
Rev. cos prediction
Cosine: 0.96
3a, 6k_2-hydroxy-4-methoxybenzaldehyde
MS/MS match
Rev. cos prediction
Cosine: 0.97
chenodeoxycholic-acid_glycine
MS/MS match
Rev. cos prediction
Cosine: 0.95
undecanoic-acid_L-alanine
MS/MS match
Rev. cos prediction
Cosine: 0.99
chenodeoxycholic-acid_5-aminosalicylic-acid
MS/MS match
Rev. cos prediction
Cosine: 0.94
2,5-dimethoxybenzoic-acid_2-ethylphenol
MS/MS match
Rev. cos prediction
Cosine: 0.94
3beta7beta12alpha_Val
MS/MS match
Rev. cos prediction
Cosine: 0.97
Erythro-aleuritic acid_L-alanine
MS/MS match
Rev. cos prediction
Cosine: 0.95
C16:1_Puterscin
MS/MS match
Rev. cos prediction
Cosine: 0.93
2-methylglutaric-acid_propane-1,3-diamine
MS/MS match
Rev. cos prediction
Cosine: 0.99
2-methylglutaric-acid_propane-1,3-diamine
MS/MS match
Rev. cos prediction
Cosine: 0.95
Succinic acid_L-glutamate
MS/MS match
Rev. cos prediction
Cosine: 0.99
(E)-2-methylbut-2-enoic-acid_L-tyrosine
MS/MS match
Rev. cos prediction
Cosine: 0.97
5-Aminosalicylic acid_Cholic Acid
MS/MS match
Rev. cos prediction
Cosine: 0.93
3-hydroxy-benzoic-acid_L-glutamine
MS/MS match
Rev. cos prediction
Cosine: 0.98
2-oxopropanoic-acid_1,4-cyclohexanediol
MS/MS match
Rev. cos prediction
Cosine: 0.95
3alpha6alpha_Tyr
MS/MS match
Rev. cos prediction
Cosine: 0.97
deoxycholic-acid_L-leucine
MS/MS match
Rev. cos prediction
Cosine: 0.95
3-methylpentanoic-acid_L-tryptophan
MS/MS match
Rev. cos prediction
Cosine: 0.98
nicotinic-acid_L-aspartate
MS/MS match
Rev. cos prediction
Cosine: 0.95
3alpha6beta7beta_Val
MS/MS match
Rev. cos prediction
Cosine: 0.98
nicotinic-acid_L-aspartate
MS/MS match
Rev. cos prediction
Cosine: 0.95
4-hydroxyphenylacetic acid_L-threonine
MS/MS match
Rev. cos prediction
Cosine: 0.94
undecanoic-acid_L-alanine
MS/MS match
Rev. cos prediction
Cosine: 0.99
cholic-acid_L-alanine
MS/MS match
Rev. cos prediction
Cosine: 0.94
Pantothenic acid_tetraethylene glycol
MS/MS match
Rev. cos prediction
Cosine: 0.98
3beta12beta_Tyr
MS/MS match
Rev. cos prediction
Cosine: 0.93
benzoic acid_agmatine dihydrochloride
MS/MS match
Rev. cos prediction
Cosine: 1.00

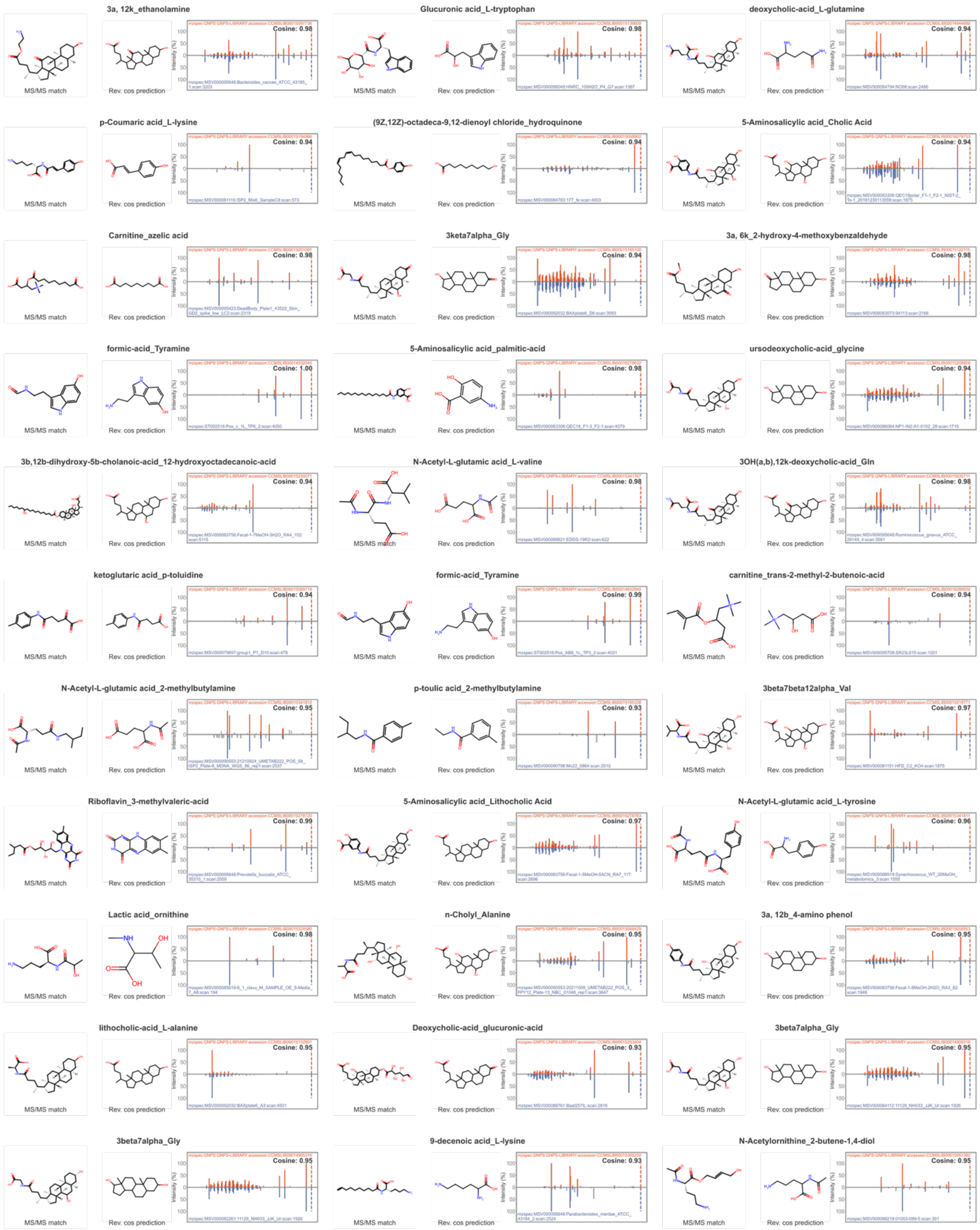
3a, 12k_ethanolamine
Cosine: 0.98
Glucuronic acid_L-tryptophan
Cosine: 0.98
deoxycholic-acid_L-glutamine
Cosine: 0.94
p-Coumaric acid_L-lysine
Cosine: 0.94
(9Z,12Z)-octadeca-9,12-dienoyl chloride_hydroquinone
Cosine: 0.94
5-Aminosalicylic acid_Cholic Acid
Cosine: 0.94
Carnitine_azelic acid
Cosine: 0.98
3keta7alpha_Gly
Cosine: 0.94
3a, 6k_2-hydroxy-4-methoxybenzaldehyde
Cosine: 0.98
formic-acid_Tyramine
Cosine: 1.00
5-Aminosalicylic acid_palmitic-acid
Cosine: 0.98
ursodeoxycholic-acid_glycine
Cosine: 0.94
3b,12b-dihydroxy-5b-cholanoic-acid_12-hydroxyoctadecanoic-acid
Cosine: 0.94
N-Acetyl-L-glutamic acid_L-valine
Cosine: 0.98
3OH(a,b),12k-deoxycholic-acid_Gln
Cosine: 0.98
ketoglutaric acid_p-toluidine
Cosine: 0.94
formic-acid_Tyramine
Cosine: 0.99
carnitine_trans-2-methyl-2-butenoic-acid
Cosine: 0.94
N-Acetyl-L-glutamic acid_2-methylbutylamine
Cosine: 0.95
p-toulic acid_2-methylbutylamine
Cosine: 0.93
3beta7beta12alpha_Val
Cosine: 0.97
Riboflavin_3-methylvaleric-acid
Cosine: 0.99
5-Aminosalicylic acid_Lithocholic Acid
Cosine: 0.97
N-Acetyl-L-glutamic acid_L-tyrosine
Cosine: 0.96
Lactic acid_ornithine
Cosine: 0.98
n-Cholyl_Alanine
Cosine: 0.98
3a, 12b_4-amino phenol
Cosine: 0.95
lithocholic-acid_L-alanine
Cosine: 0.95
Deoxycholic-acid_glucuronic-acid
Cosine: 0.93
3beta7alpha_Gly
Cosine: 0.95
3beta7alpha_Gly
Cosine: 0.95
9-decenoic acid_L-lysine
Cosine: 0.93
N-Acetylornithine_2-butene-1,4-diol
Cosine: 0.95
MS/MS match
Rev. cos prediction

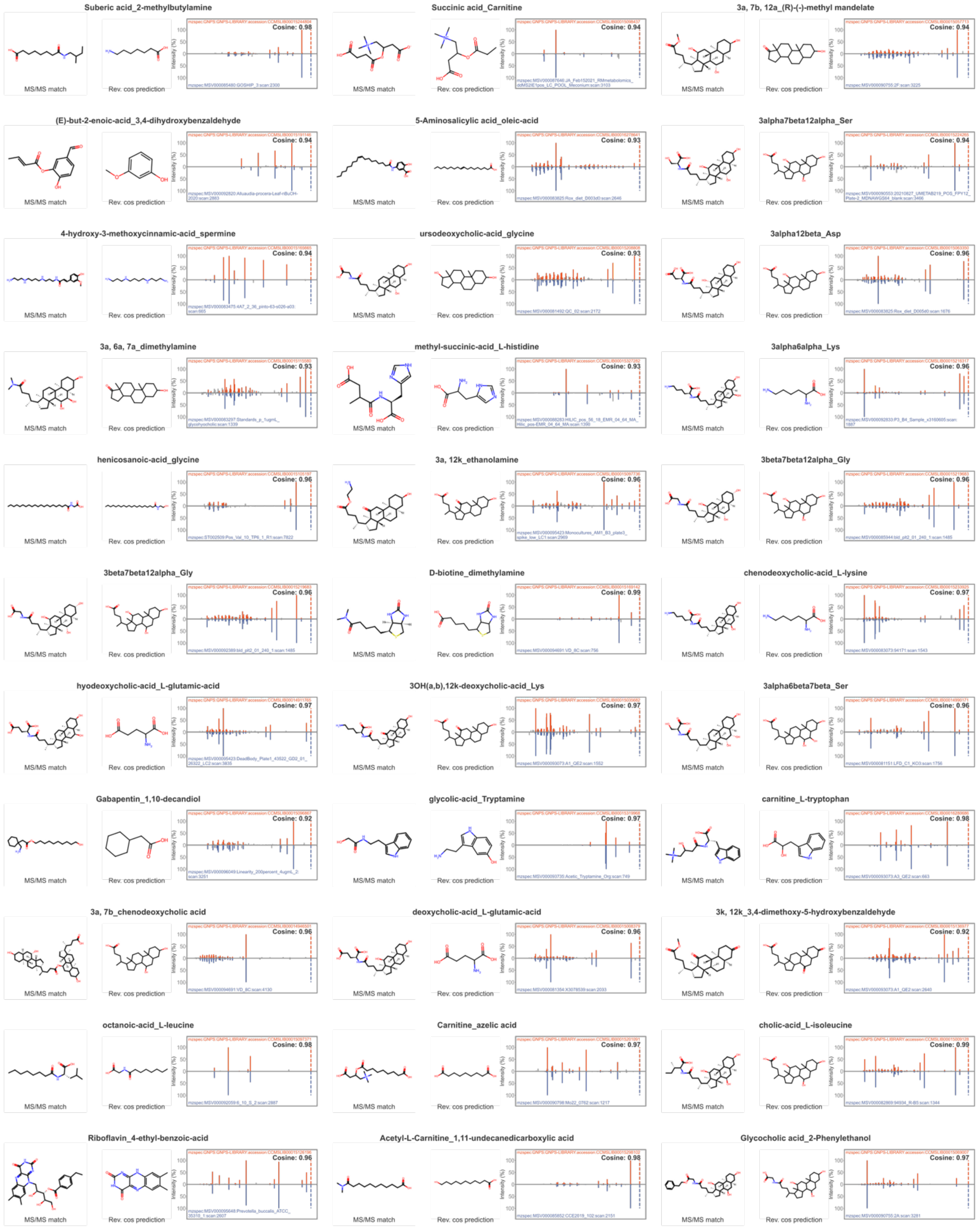

Suberic acid_2-methylbutylamine
Cosine: 0.98
Succinic acid_Carnitine
Cosine: 0.94
3a, 7b, 12a_(R)-(-)-methyl mandelate
Cosine: 0.94
(E)-but-2-enoic-acid_3,4-dihydroxybenzaldehyde
Cosine: 0.94
5-Aminosalicylic acid_oleic-acid
Cosine: 0.93
3alpha7beta12alpha_Ser
Cosine: 0.94
4-hydroxy-3-methoxycinnamic-acid_spermine
Cosine: 0.94
ursodeoxycholic-acid_glycine
Cosine: 0.93
3alpha12beta_Asp
Cosine: 0.96
3a, 6a, 7a_dimethylamine
Cosine: 0.93
methyl-succinic-acid_L-histidine
Cosine: 0.93
3alpha6alpha_Lys
Cosine: 0.96
henicosanoic-acid_glycine
Cosine: 0.96
3a, 12k_ethanolamine
Cosine: 0.96
3beta7beta12alpha_Gly
Cosine: 0.96
3beta7beta12alpha_Gly
Cosine: 0.96
D-biotine_dimethylamine
Cosine: 0.99
chenodeoxycholic-acid_L-lysine
Cosine: 0.97
hyodeoxycholic-acid_L-glutamic-acid
Cosine: 0.97
3OH(a,b),12k-deoxycholic-acid_Lys
Cosine: 0.97
3alpha6beta7beta_Ser
Cosine: 0.96
Gabapentin_1,10-decandiol
Cosine: 0.92
glycolic-acid_Tryptamine
Cosine: 0.97
carnitine_L-tryptophan
Cosine: 0.98
3a, 7b_chenodeoxycholic acid
Cosine: 0.96
deoxycholic-acid_L-glutamic-acid
Cosine: 0.96
3k, 12k_3,4-dimethoxy-5-hydroxybenzaldehyde
Cosine: 0.92
octanoic-acid_L-leucine
Cosine: 0.98
Carnitine_azelic acid
Cosine: 0.97
cholic-acid_L-isoleucine
Cosine: 0.99
Riboflavin_4-ethyl-benzoic-acid
Cosine: 0.96
Acetyl-L-Carnitine_1,11-undecanedicarboxylic acid
Cosine: 0.98
Glycocholic acid_2-Phenylethanol
Cosine: 0.97

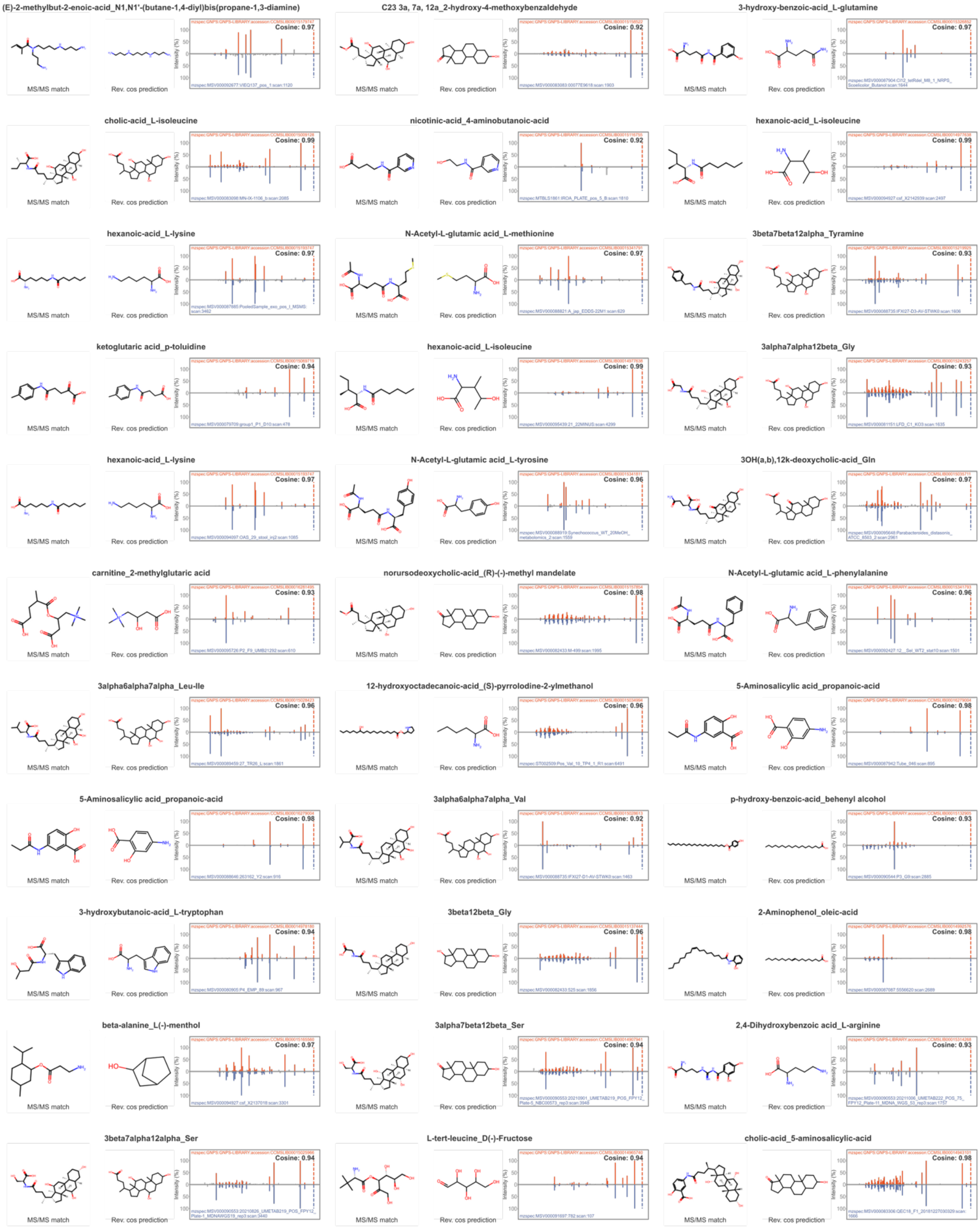

(E)-2-methylbut-2-enoic-acid_N1,N1'-(butane-1,4-diyl)bis(propane-1,3-diamine)
MS/MS match
Rev. cos prediction
Cosine: 0.97
C23 3a, 7a, 12a_2-hydroxy-4-methoxybenzaldehyde
MS/MS match
Rev. cos prediction
Cosine: 0.92
3-hydroxy-benzoic-acid_L-glutamine
MS/MS match
Rev. cos prediction
Cosine: 0.97
cholic-acid_L-isoleucine
MS/MS match
Rev. cos prediction
Cosine: 0.99
nicotinic-acid_4-aminobutanoic-acid
MS/MS match
Rev. cos prediction
Cosine: 0.92
hexanoic-acid_L-isoleucine
MS/MS match
Rev. cos prediction
Cosine: 0.99
hexanoic-acid_L-lysine
MS/MS match
Rev. cos prediction
Cosine: 0.97
N-Acetyl-L-glutamic acid_L-methionine
MS/MS match
Rev. cos prediction
Cosine: 0.97
3beta7beta12alpha_Tyramine
MS/MS match
Rev. cos prediction
Cosine: 0.93
ketoglutaric acid_p-toluidine
MS/MS match
Rev. cos prediction
Cosine: 0.94
hexanoic-acid_L-isoleucine
MS/MS match
Rev. cos prediction
Cosine: 0.99
3alpha7alpha12beta_Gly
MS/MS match
Rev. cos prediction
Cosine: 0.93
hexanoic-acid_L-lysine
MS/MS match
Rev. cos prediction
Cosine: 0.97
N-Acetyl-L-glutamic acid_L-tyrosine
MS/MS match
Rev. cos prediction
Cosine: 0.96
3OH(a,b),12k-deoxycholic-acid_Gln
MS/MS match
Rev. cos prediction
Cosine: 0.97
carnitine_2-methylglutaric acid
MS/MS match
Rev. cos prediction
Cosine: 0.93
norursodeoxycholic-acid_(R)-(-)-methyl mandelate
MS/MS match
Rev. cos prediction
Cosine: 0.98
N-Acetyl-L-glutamic acid_L-phenylalanine
MS/MS match
Rev. cos prediction
Cosine: 0.96
3alpha6alpha7alpha_Leu-Ile
MS/MS match
Rev. cos prediction
Cosine: 0.96
12-hydroxyoctadecanoic-acid_(S)-pyrrolodine-2-ylmethanol
MS/MS match
Rev. cos prediction
Cosine: 0.96
5-Aminosalicylic acid_propanoic-acid
MS/MS match
Rev. cos prediction
Cosine: 0.98
5-Aminosalicylic acid_propanoic-acid
MS/MS match
Rev. cos prediction
Cosine: 0.98
3alpha6alpha7alpha_Val
MS/MS match
Rev. cos prediction
Cosine: 0.92
p-hydroxy-benzoic-acid_behenyl alcohol
MS/MS match
Rev. cos prediction
Cosine: 0.93
3-hydroxybutanoic-acid_L-tryptophan
MS/MS match
Rev. cos prediction
Cosine: 0.94
3beta12beta_Gly
MS/MS match
Rev. cos prediction
Cosine: 0.96
2-Aminophenol_oleic-acid
MS/MS match
Rev. cos prediction
Cosine: 0.98
beta-alanine_L(-)-menthol
MS/MS match
Rev. cos prediction
Cosine: 0.97
3alpha7beta12beta_Ser
MS/MS match
Rev. cos prediction
Cosine: 0.94
2,4-Dihydroxybenzoic acid_L-arginine
MS/MS match
Rev. cos prediction
Cosine: 0.93
3beta7alpha12alpha_Ser
MS/MS match
Rev. cos prediction
Cosine: 0.94
L-tert-leucine_D(-)-Fructose
MS/MS match
Rev. cos prediction
Cosine: 0.94
cholic-acid_5-aminosalicylic-acid
MS/MS match
Rev. cos prediction
Cosine: 0.98

3-Indolepropionic acid_L-methionine

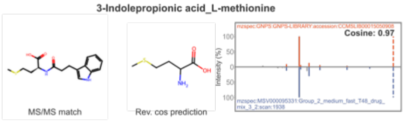


3beta12beta_Ser

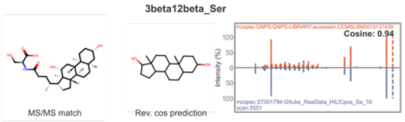


C23 3a, 7a, 12a_2-hydroxy-4-methoxybenzaldehyde

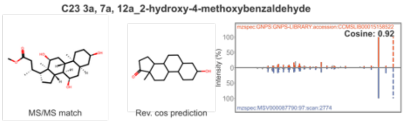


Suberic acid_2-methylbutylamine

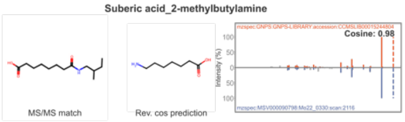


lactic-acid_L-tryptophan

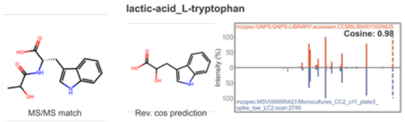


12-hydroxyoctadecanoic-acid_(S)-pyrrolodine-2-ylmethanol

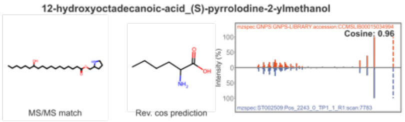


3beta7beta12alpha_Ser

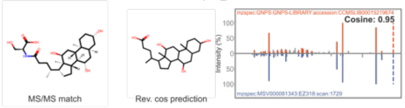


lactic-acid_L-arginine

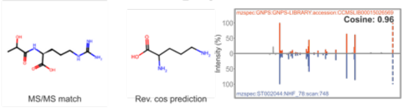


Carnitine_pimelic acid

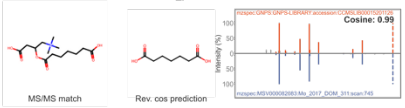


carnitine_trans-2-methyl-2-butenoic-acid

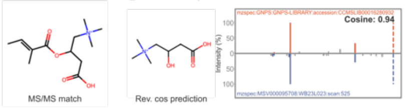


3-methylbutanoic-acid_4-aminobutanoic acid

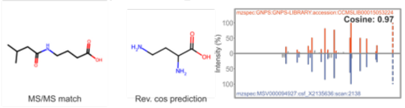


5-Aminosalicylic acid_hexanoic-acid

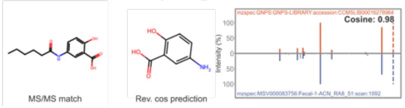


cholic-acid_L-glutamic-acid

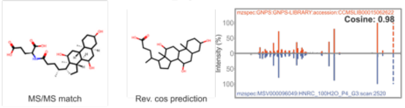


chenodeoxycholic-acid_L-serine

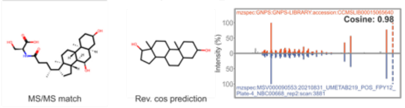


Succinic acid_L-glutamate

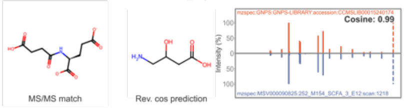


3beta7alpha_4-aminobutyric acid

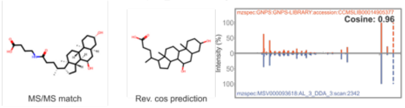


Pyroglutamic acid_L-leucine

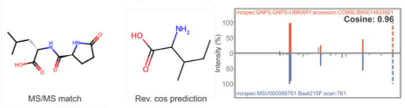


3beta7alpha12alpha_Cadaverine

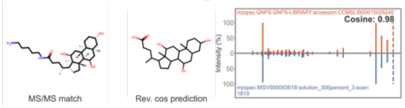


deoxycholic-acid_2-aminophenol

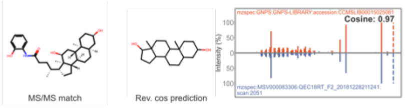


3k,12OH-deoxycholic-acid_Gly

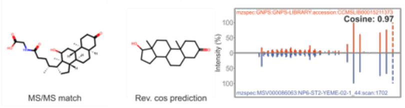


ursodeoxycholic-acid_L-leucine

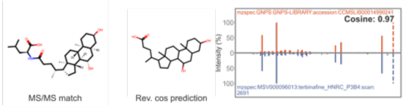


chenodeoxycholic-acid_L-glutamic-acid

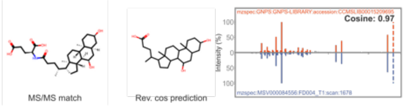


Succnic acid_2-ethylaniline

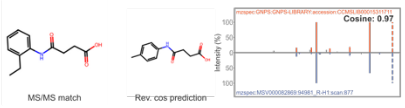


12-methyltridecanoic-acid_2-amino-2-methyl-1-propanol

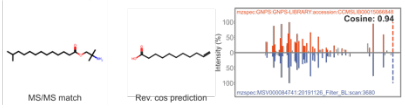


Taurine_dodecanedioic acid

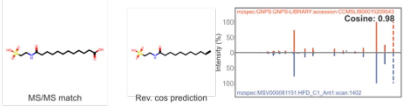


3beta7beta12alpha_Ser

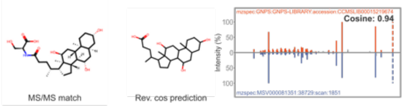


n-Cholyl_Valine

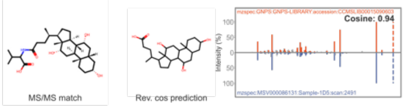


cholic acid_L-tryptophan

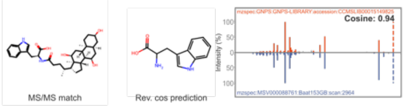


hentriacontanoic-acid_D-serine

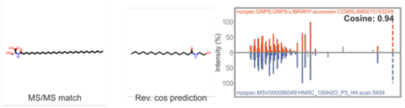


3alpha6beta7beta_Thr

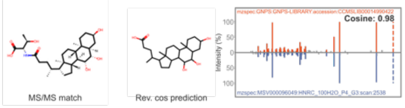


chenodeoxycholic-acid_L-isoleucine

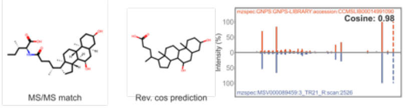


lactic-acid_pentane-1,5-diamine

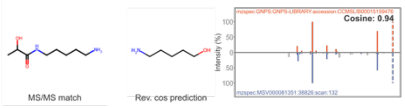


3beta7beta12alpha_4-aminobutyric acid

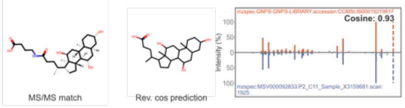

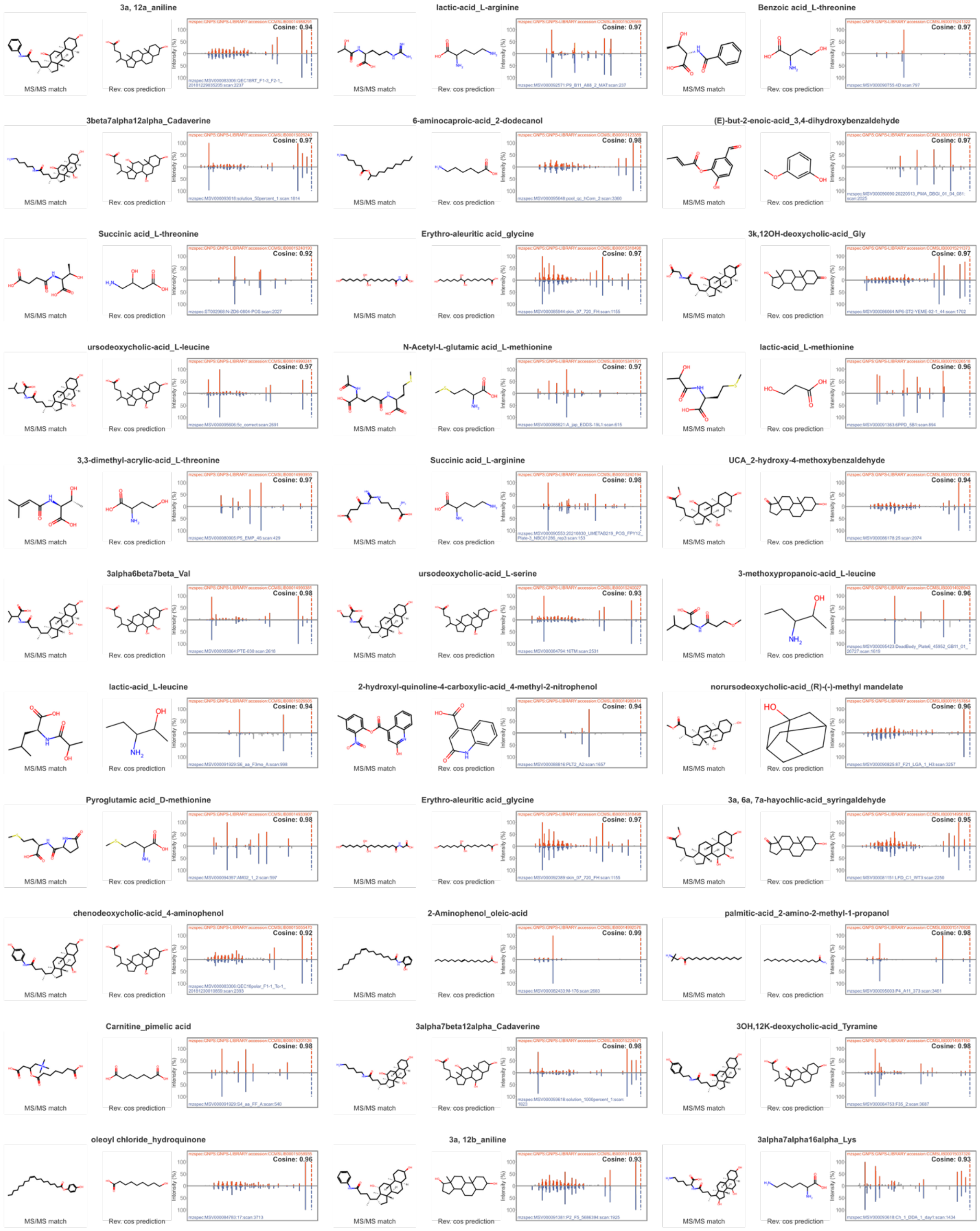
3a, 12a_aniline
Cosine: 0.94
lactic-acid_L-arginine
Cosine: 0.97
Benzoic acid_L-threonine
Cosine: 0.97
3beta7alpha12alpha_Cadaverine
Cosine: 0.97
6-aminocaproic-acid_2-dodecanol
Cosine: 0.98
(E)-but-2-enoic-acid_3,4-dihydroxybenzaldehyde
Cosine: 0.97
Succinic acid_L-threonine
Cosine: 0.92
Erythro-aleuritic acid_glycine
Cosine: 0.97
3k,12OH-deoxycholic-acid_Gly
Cosine: 0.97
ursodeoxycholic-acid_L-leucine
Cosine: 0.97
N-Acetyl-L-glutamic acid_L-methionine
Cosine: 0.97
lactic-acid_L-methionine
Cosine: 0.96
3,3-dimethyl-acrylic-acid_L-threonine
Cosine: 0.97
Succinic acid_L-arginine
Cosine: 0.98
UCA_2-hydroxy-4-methoxybenzaldehyde
Cosine: 0.94
3alpha6beta7beta_Val
Cosine: 0.98
ursodeoxycholic-acid_L-serine
Cosine: 0.93
3-methoxypropanoic-acid_L-leucine
Cosine: 0.96
lactic-acid_L-leucine
Cosine: 0.94
2-hydroxyl-quinoline-4-carboxylic-acid_4-methyl-2-nitrophenol
Cosine: 0.94
norursodeoxycholic-acid_(R)-(-)-methyl mandelate
Cosine: 0.96
Pyroglutamic acid_D-methionine
Cosine: 0.98
Erythro-aleuritic acid_glycine
Cosine: 0.97
3a, 6a, 7a-hayochlic-acid_syringaldehyde
Cosine: 0.95
chenodeoxycholic-acid_4-aminophenol
Cosine: 0.92
2-Aminophenol_oleic-acid
Cosine: 0.92
palmitic-acid_2-amino-2-methyl-1-propanol
Cosine: 0.98
Carnitine_pimelic acid
Cosine: 0.98
3alpha7beta12alpha_Cadaverine
Cosine: 0.98
3OH,12K-deoxycholic-acid_Tyramine
Cosine: 0.98
oleoyl chloride_hydroquinone
Cosine: 0.96
3a, 12b_aniline
Cosine: 0.93
3alpha7alpha16alpha_Lys
Cosine: 0.93
MS/MS match
Rev. cos prediction

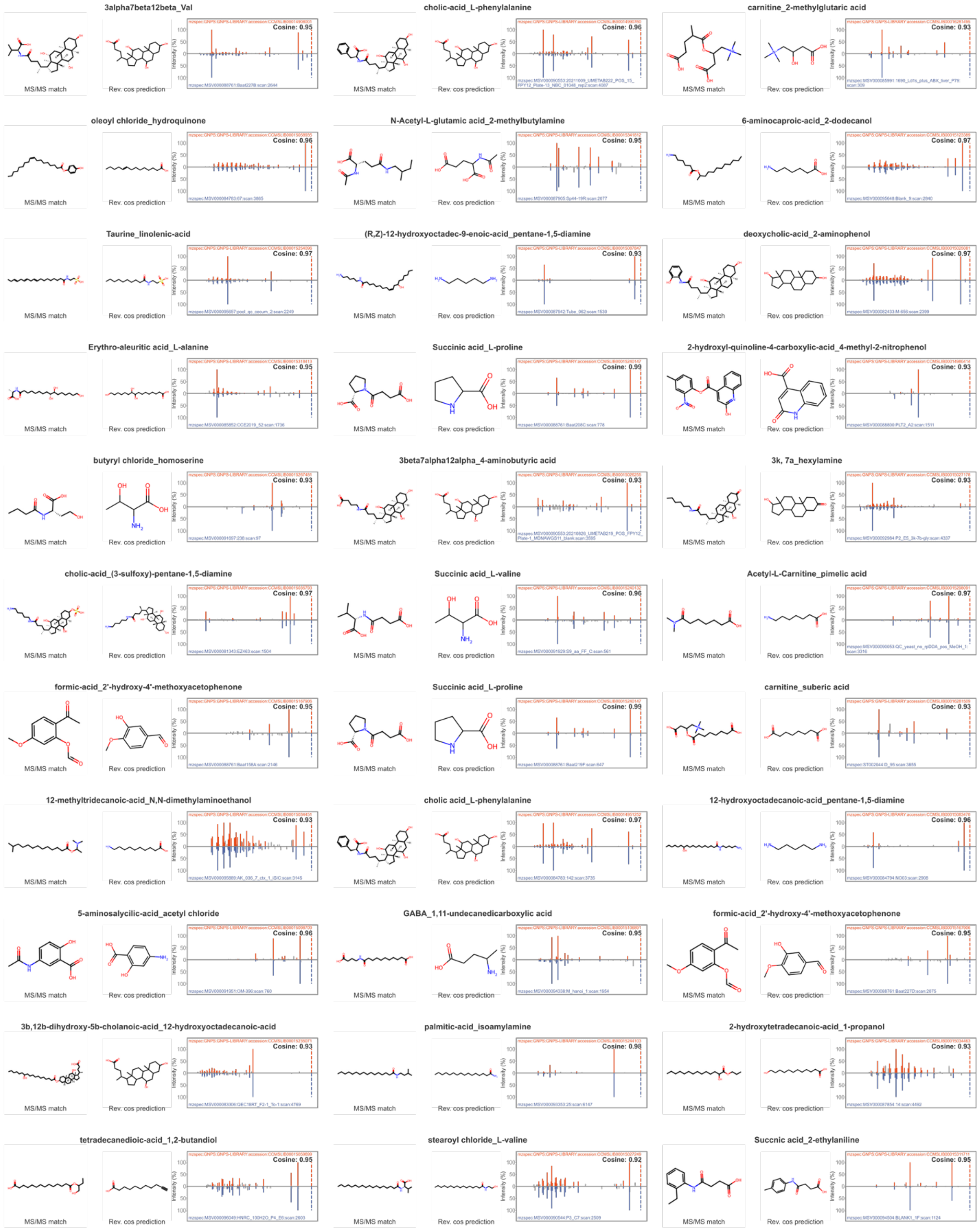
3alpha7beta12beta_Val
Cosine: 0.95
cholic-acid_L-phenylalanine
Cosine: 0.96
carnitine_2-methylglutaric acid
Cosine: 0.93
oleoyl chloride_hydroquinone
Cosine: 0.96
N-Acetyl-L-glutamic acid_2-methylbutylamine
Cosine: 0.95
6-aminocaproic-acid_2-dodecanol
Cosine: 0.97
Taurine_linolenic-acid
Cosine: 0.97
(R,Z)-12-hydroxyoctadec-9-enoic-acid_pentane-1,5-diamine
Cosine: 0.93
deoxycholic-acid_2-aminophenol
Cosine: 0.97
Erythro-aleuritic acid_L-alanine
Cosine: 0.95
Succinic acid_L-proline
Cosine: 0.99
2-hydroxyl-quinoline-4-carboxylic-acid_4-methyl-2-nitrophenol
Cosine: 0.93
butyryl chloride_homoserine
Cosine: 0.93
3beta7alpha12alpha_4-aminobutyric acid
Cosine: 0.93
3k, 7a_hexylamine
Cosine: 0.93
cholic-acid_(3-sulfoxy)-pentane-1,5-diamine
Cosine: 0.97
Succinic acid_L-valine
Cosine: 0.96
Acetyl-L-Carnitine_pimelic acid
Cosine: 0.97
formic-acid_2'-hydroxy-4'-methoxyacetophenone
Cosine: 0.95
Succinic acid_L-proline
Cosine: 0.99
carnitine_suberic acid
Cosine: 0.93
12-methyltridecanoic-acid_N,N-dimethylaminoethanol
Cosine: 0.93
cholic acid_L-phenylalanine
Cosine: 0.97
12-hydroxyoctadecanoic-acid_pentane-1,5-diamine
Cosine: 0.96
5-aminosalycilic-acid_acetyl chloride
Cosine: 0.96
GABA_1,11-undecanedicarboxylic acid
Cosine: 0.97
formic-acid_2'-hydroxy-4'-methoxyacetophenone
Cosine: 0.95
3b,12b-dihydroxy-5b-cholanoic-acid_12-hydroxyoctadecanoic-acid
Cosine: 0.93
palmitic-acid_isoamylamine
Cosine: 0.98
2-hydroxytetradecanoic-acid_1-propanol
Cosine: 0.93
tetradecanedioic-acid_1,2-butandiol
Cosine: 0.95
stearoyl chloride_L-valine
Cosine: 0.92
Succnic acid_2-ethylaniline
Cosine: 0.95
MS/MS match
Rev. cos prediction
Intensity (%)

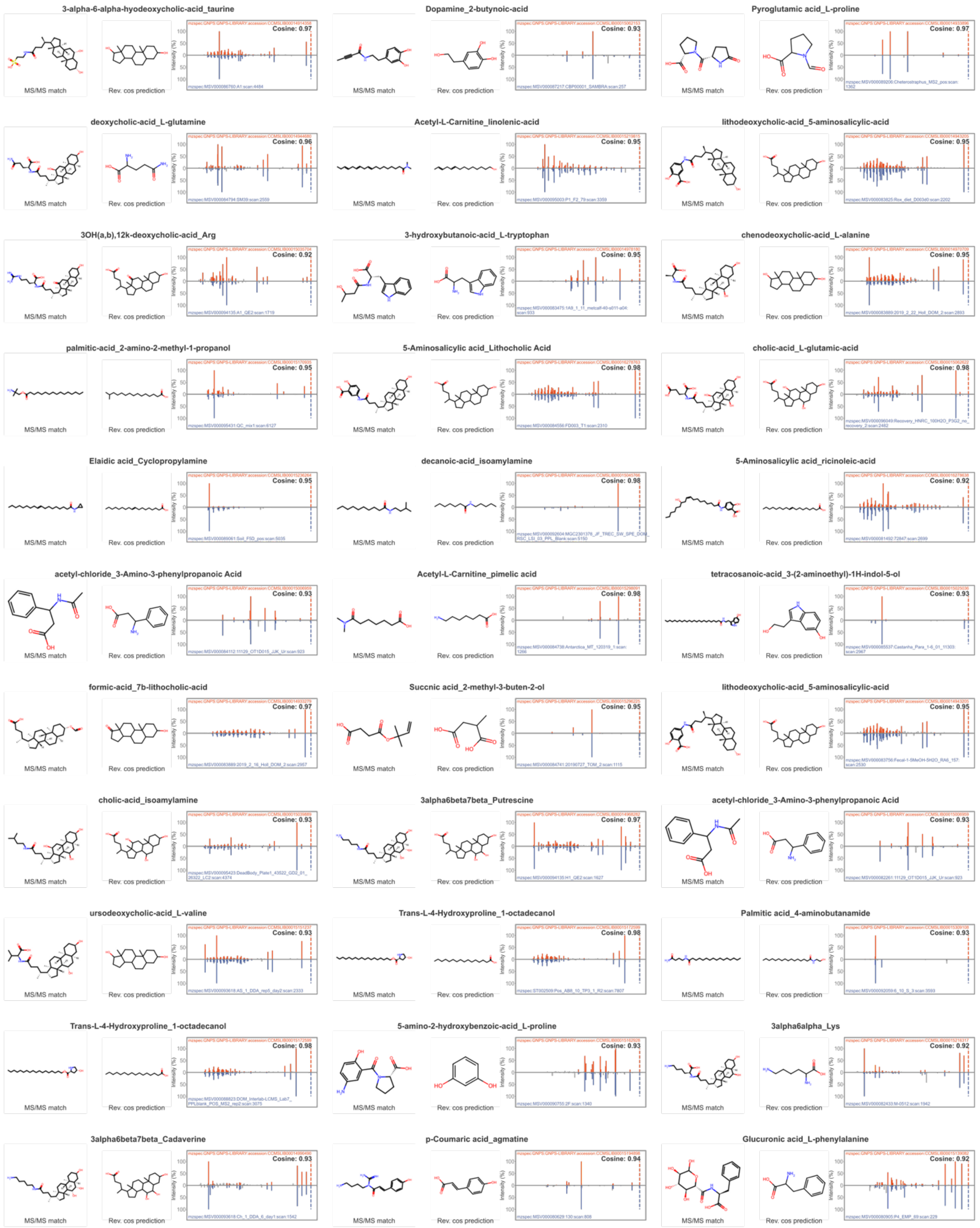
3-alpha-6-alpha-hyodeoxycholic-acid_taurine
Cosine: 0.97
Dopamine_2-butynoic-acid
Cosine: 0.93
Pyroglutamic acid_L-proline
Cosine: 0.97
deoxycholic-acid_L-glutamine
Cosine: 0.96
Acetyl-L-Carnitine_linolenic-acid
Cosine: 0.95
lithodeoxycholic-acid_5-aminosalicylic-acid
Cosine: 0.95
3OH(a,b),12k-deoxycholic-acid_Arg
Cosine: 0.92
3-hydroxybutanoic-acid_L-tryptophan
Cosine: 0.95
chenodeoxycholic-acid_L-alanine
Cosine: 0.95
palmitic-acid_2-amino-2-methyl-1-propanol
Cosine: 0.95
5-Aminosalicylic acid_Lithocholic Acid
Cosine: 0.98
cholic-acid_L-glutamic-acid
Cosine: 0.98
Elaidic acid_Cyclopropylamine
Cosine: 0.95
decanoic-acid_isoamylamine
Cosine: 0.98
5-Aminosalicylic acid_ricinoleic-acid
Cosine: 0.92
acetyl-chloride_3-Amino-3-phenylpropanoic Acid
Cosine: 0.93
Acetyl-L-Carnitine_pimelic acid
Cosine: 0.98
tetracosanoic-acid_3-(2-aminoethyl)-1H-indol-5-ol
Cosine: 0.93
formic-acid_7b-lithocholic-acid
Cosine: 0.97
Succnic acid_2-methyl-3-buten-2-ol
Cosine: 0.95
lithodeoxycholic-acid_5-aminosalicylic-acid
Cosine: 0.95
cholic-acid_isoamylamine
Cosine: 0.93
3alpha6beta7beta_Putrescine
Cosine: 0.97
acetyl-chloride_3-Amino-3-phenylpropanoic Acid
Cosine: 0.93
ursodeoxycholic-acid_L-valine
Cosine: 0.93
Trans-L-4-Hydroxyproline_1-octadecanol
Cosine: 0.98
Palmitic acid_4-aminobutanamide
Cosine: 0.93
Trans-L-4-Hydroxyproline_1-octadecanol
Cosine: 0.98
5-amino-2-hydroxybenzoic-acid_L-proline
Cosine: 0.93
3alpha6alpha_Lys
Cosine: 0.92
3alpha6beta7beta_Cadaverine
Cosine: 0.93
p-Coumaric acid_agmatine
Cosine: 0.94
Glucuronic acid_L-phenylalanine
Cosine: 0.92
MS/MS match
Rev. cos prediction
Intensity (%)

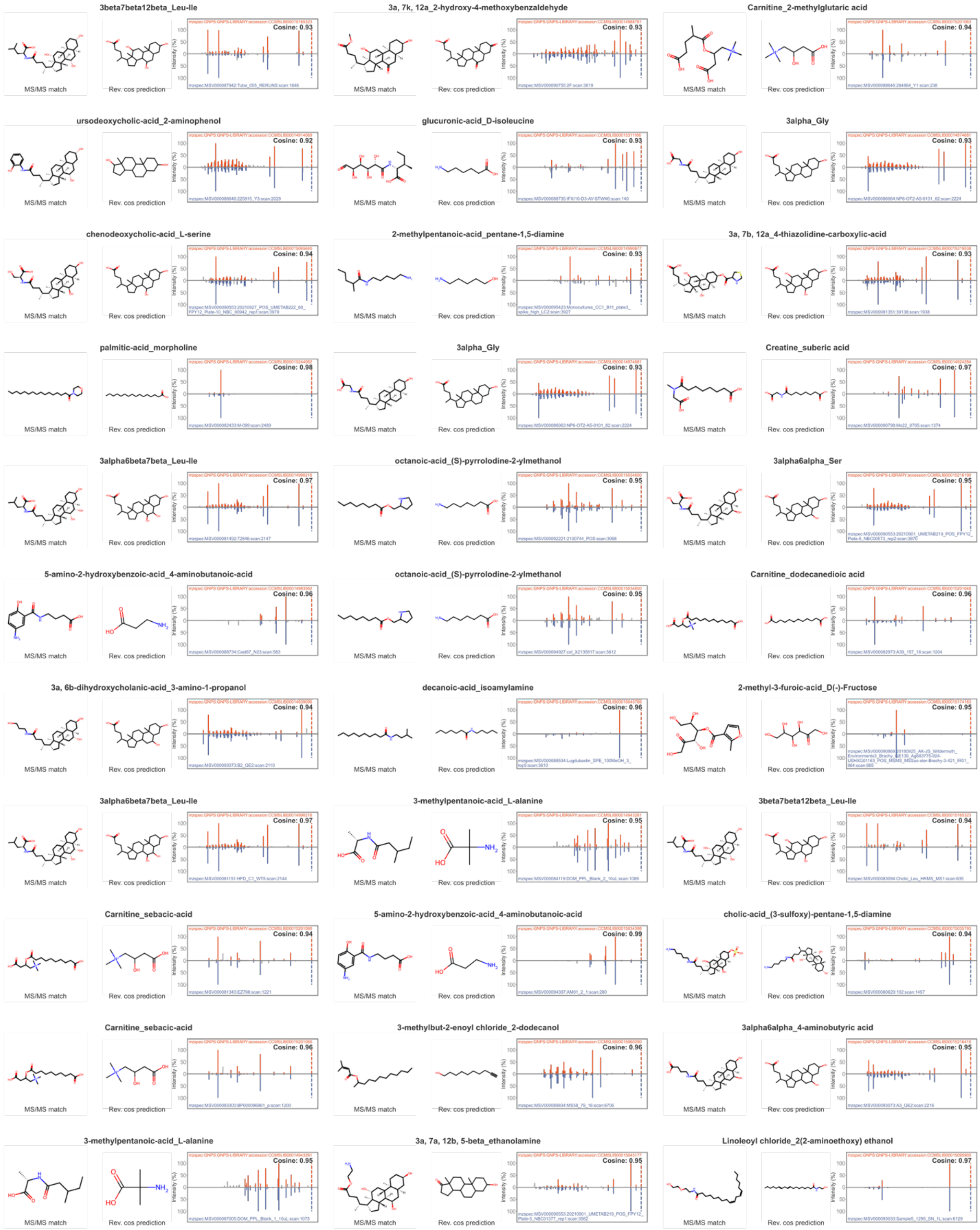
3beta7beta12beta_Leu-Ile
Cosine: 0.93
3a, 7k, 12a_2-hydroxy-4-methoxybenzaldehyde
Cosine: 0.93
Carnitine_2-methylglutaric acid
Cosine: 0.94
ursodeoxycholic-acid_2-aminophenol
Cosine: 0.92
glucuronic-acid_D-isoleucine
Cosine: 0.93
3alpha_Gly
Cosine: 0.93
chenodeoxycholic-acid_L-serine
Cosine: 0.94
2-methylpentanoic-acid_pentane-1,5-diamine
Cosine: 0.93
3a, 7b, 12a_4-thiazolidine-carboxylic-acid
Cosine: 0.93
palmitic-acid_morpholine
Cosine: 0.98
3alpha_Gly
Cosine: 0.93
Creatine_suberic acid
Cosine: 0.97
3alpha6beta7beta_Leu-Ile
Cosine: 0.97
octanoic-acid_(S)-pyrrolodine-2-ylmethanol
Cosine: 0.95
3alpha6alpha_Ser
Cosine: 0.95
5-amino-2-hydroxybenzoic-acid_4-aminobutanoic-acid
Cosine: 0.96
octanoic-acid_(S)-pyrrolodine-2-ylmethanol
Cosine: 0.95
Carnitine_dodecanedioic acid
Cosine: 0.96
3a, 6b-dihydroxycholanic-acid_3-amino-1-propanol
Cosine: 0.94
decanoic-acid_isoamylamine
Cosine: 0.96
2-methyl-3-furoic-acid_D(-)-Fructose
Cosine: 0.95
3alpha6beta7beta_Leu-Ile
Cosine: 0.97
3-methylpentanoic-acid_L-alanine
Cosine: 0.95
3beta7beta12beta_Leu-Ile
Cosine: 0.94
Carnitine_sebacic-acid
Cosine: 0.94
5-amino-2-hydroxybenzoic-acid_4-aminobutanoic-acid
Cosine: 0.99
cholic-acid_(3-sulfoxy)-pentane-1,5-diamine
Cosine: 0.94
Carnitine_sebacic-acid
Cosine: 0.96
3-methylbut-2-enoyl chloride_2-dodecanol
Cosine: 0.96
3alpha6alpha_4-aminobutyric acid
Cosine: 0.95
3-methylpentanoic-acid_L-alanine
Cosine: 0.95
3a, 7a, 12b, 5-beta_ethanolamine
Cosine: 0.95
Linoleoyl chloride_2(2-aminoethoxy) ethanol
Cosine: 0.97
MS/MS match
Rev. cos prediction
Intensity (%)

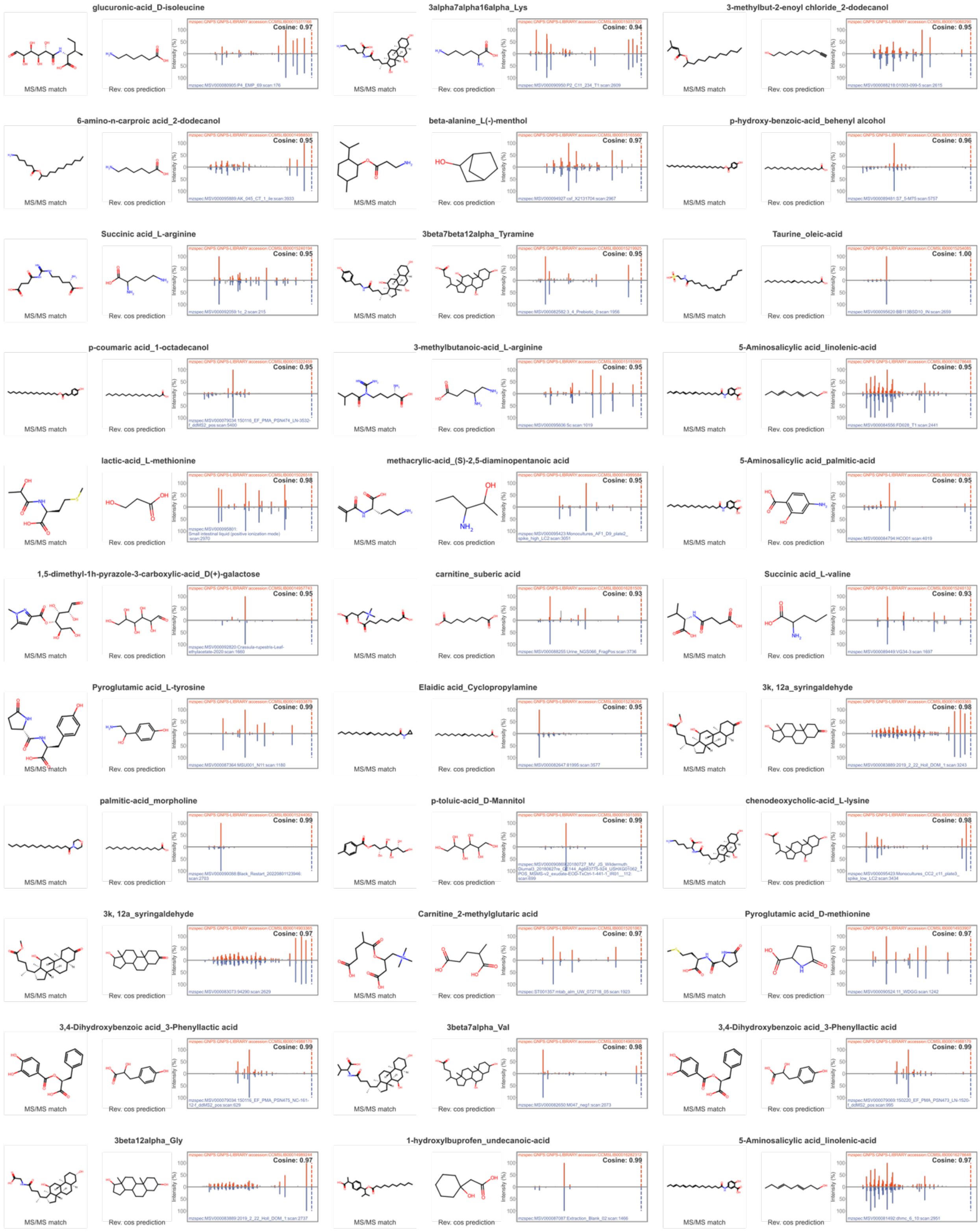
glucuronic-acid_D-isoleucine
Cosine: 0.97
3alpha7alpha16alpha_Lys
Cosine: 0.94
3-methylbut-2-enoyl chloride_2-dodecanol
Cosine: 0.95
6-amino-n-carproic acid_2-dodecanol
Cosine: 0.95
beta-alanine_L(-)-menthol
Cosine: 0.97
p-hydroxy-benzoic-acid_behenyl alcohol
Cosine: 0.96
Succinic acid_L-arginine
Cosine: 0.95
3beta7beta12alpha_Tyramine
Cosine: 0.95
Taurine_oleic-acid
Cosine: 1.00
p-coumaric acid_1-octadecanol
Cosine: 0.95
3-methylbutanoic-acid_L-arginine
Cosine: 0.95
5-Aminosalicylic acid_linolenic-acid
Cosine: 0.95
lactic-acid_L-methionine
Cosine: 0.98
methacrylic-acid_(S)-2,5-diaminopentanoic acid
Cosine: 0.95
5-Aminosalicylic acid_palmitic-acid
Cosine: 0.95
1,5-dimethyl-1h-pyrazole-3-carboxylic-acid_D(+)-galactose
Cosine: 0.95
carnitine_suberic acid
Cosine: 0.93
Succinic acid_L-valine
Cosine: 0.93
Pyroglutamic acid_L-tyrosine
Cosine: 0.99
Elaidic acid_Cyclopropylamine
Cosine: 0.95
3k, 12a_syringaldehyde
Cosine: 0.98
palmitic-acid_morpholine
Cosine: 0.99
p-toluic-acid_D-Mannitol
Cosine: 0.99
chenodeoxycholic-acid_L-lysine
Cosine: 0.98
3k, 12a_syringaldehyde
Cosine: 0.97
Carnitine_2-methylglutaric acid
Cosine: 0.97
Pyroglutamic acid_D-methionine
Cosine: 0.97
3,4-Dihydroxybenzoic acid_3-Phenyllactic acid
Cosine: 0.99
3beta7alpha_Val
Cosine: 0.98
3,4-Dihydroxybenzoic acid_3-Phenyllactic acid
Cosine: 0.99
3beta12alpha_Gly
Cosine: 0.97
1-hydroxyIbuprofen_undecanoic-acid
Cosine: 0.99
5-Aminosalicylic acid_linolenic-acid
Cosine: 0.97
MS/MS match
Rev. cos prediction
Intensity (%)

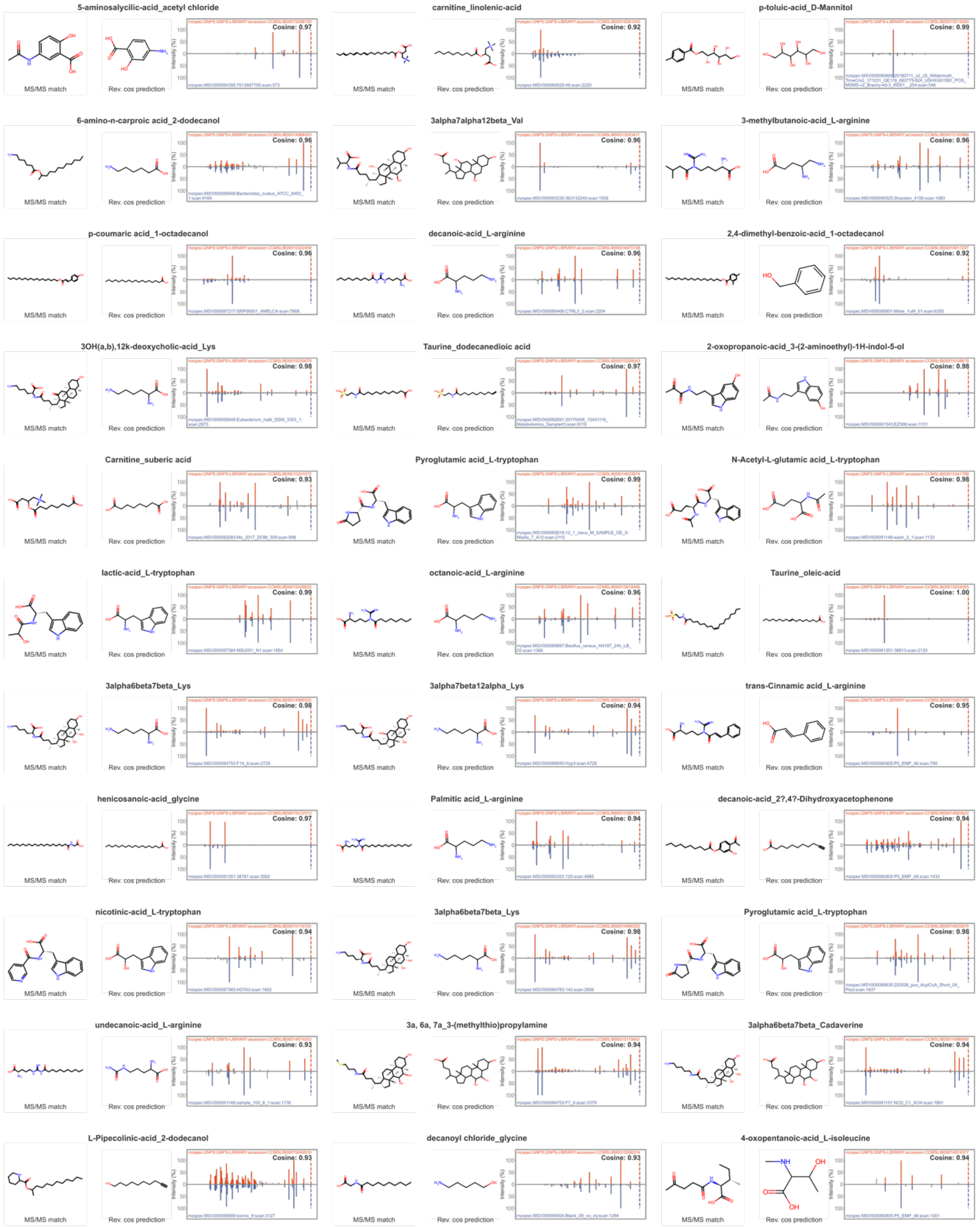

5-aminosalycilic-acid_acetyl chloride
Cosine: 0.97
carnitine_linolenic-acid
Cosine: 0.92
p-toluic-acid_D-Mannitol
Cosine: 0.99
6-amino-n-carproic acid_2-dodecanol
Cosine: 0.96
3alpha7alpha12beta_Val
Cosine: 0.96
3-methylbutanoic-acid_L-arginine
Cosine: 0.96
p-coumaric acid_1-octadecanol
Cosine: 0.96
decanoic-acid_L-arginine
Cosine: 0.96
2,4-dimethyl-benzoic-acid_1-octadecanol
Cosine: 0.92
3OH(a,b),12k-deoxycholic-acid_Lys
Cosine: 0.98
Taurine_dodecanedioic acid
Cosine: 0.97
2-oxopropanoic-acid_3-(2-aminoethyl)-1H-indol-5-ol
Cosine: 0.98
Carnitine_suberic acid
Cosine: 0.93
Pyroglutamic acid_L-tryptophan
Cosine: 0.99
N-Acetyl-L-glutamic acid_L-tryptophan
Cosine: 0.98
lactic-acid_L-tryptophan
Cosine: 0.99
octanoic-acid_L-arginine
Cosine: 0.99
Taurine_oleic-acid
Cosine: 1.00
3alpha6beta7beta_Lys
Cosine: 0.98
3alpha7beta12alpha_Lys
Cosine: 0.94
trans-Cinnamic acid_L-arginine
Cosine: 0.95
henicosanoic-acid_glycine
Cosine: 0.97
Palmitic acid_L-arginine
Cosine: 0.94
decanoic-acid_2?,4?-Dihydroxyacetophenone
Cosine: 0.94
nicotinic-acid_L-tryptophan
Cosine: 0.94
3alpha6beta7beta_Lys
Cosine: 0.98
Pyroglutamic acid_L-tryptophan
Cosine: 0.98
undecanoic-acid_L-arginine
Cosine: 0.93
3a, 6a, 7a_3-(methylthio)propylamine
Cosine: 0.94
3alpha6beta7beta_Cadaverine
Cosine: 0.94
L-Pipecolinic-acid_2-dodecanol
Cosine: 0.93
decanoyl chloride_glycine
Cosine: 0.93
4-oxopentanoic-acid_L-isoleucine
Cosine: 0.94
MS/MS match
Rev. cos prediction

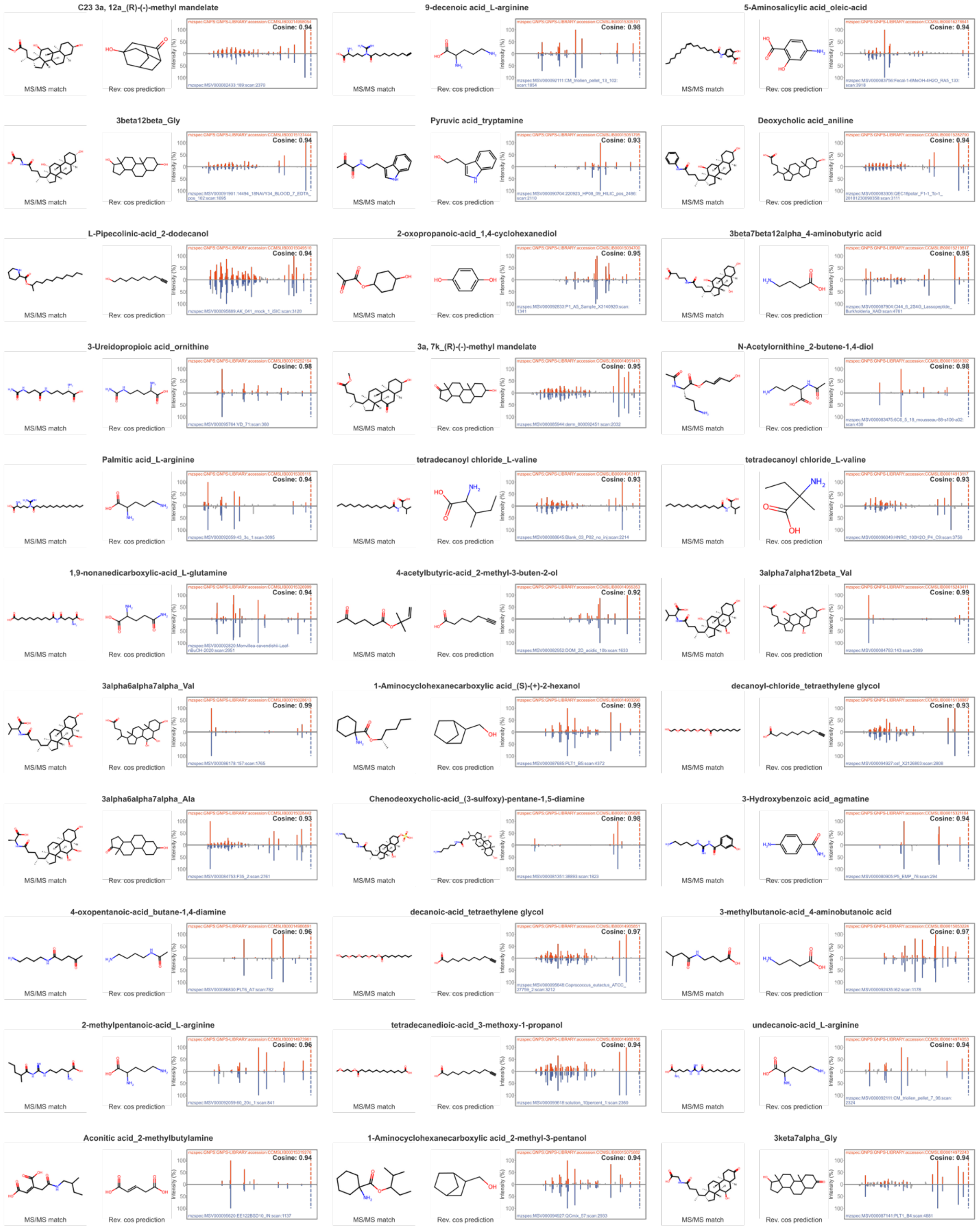
C23 3a, 12a_(R)-(-)-methyl mandelate
Cosine: 0.94
MS/MS match
Rev. cos prediction
9-decenoic acid_L-arginine
Cosine: 0.98
MS/MS match
Rev. cos prediction
5-Aminosalicylic acid_oleic-acid
Cosine: 0.94
MS/MS match
Rev. cos prediction
3beta12beta_Gly
Cosine: 0.94
MS/MS match
Rev. cos prediction
Pyruvic acid_tryptamine
Cosine: 0.93
MS/MS match
Rev. cos prediction
Deoxycholic acid_aniline
Cosine: 0.94
MS/MS match
Rev. cos prediction
L-Pipecolinic-acid_2-dodecanol
Cosine: 0.94
MS/MS match
Rev. cos prediction
2-oxopropanoic-acid_1,4-cyclohexanediol
Cosine: 0.95
MS/MS match
Rev. cos prediction
3beta7beta12alpha_4-aminobutyric acid
Cosine: 0.95
MS/MS match
Rev. cos prediction
3-Ureidopropioic acid_ornithine
Cosine: 0.98
MS/MS match
Rev. cos prediction
3a, 7k_(R)-(-)-methyl mandelate
Cosine: 0.98
MS/MS match
Rev. cos prediction
N-Acetylornithine_2-butene-1,4-diol
Cosine: 0.98
MS/MS match
Rev. cos prediction
Palmitic acid_L-arginine
Cosine: 0.94
MS/MS match
Rev. cos prediction
tetradecanoyl chloride_L-valine
Cosine: 0.93
MS/MS match
Rev. cos prediction
tetradecanoyl chloride_L-valine
Cosine: 0.93
MS/MS match
Rev. cos prediction
1,9-nonanedicarboxylic-acid_L-glutamine
Cosine: 0.94
MS/MS match
Rev. cos prediction
4-acetylbutyric-acid_2-methyl-3-buten-2-ol
Cosine: 0.92
MS/MS match
Rev. cos prediction
3alpha7alpha12beta_Val
Cosine: 0.99
MS/MS match
Rev. cos prediction
3alpha6alpha7alpha_Val
Cosine: 0.99
MS/MS match
Rev. cos prediction
1-Aminocyclohexanecarboxylic acid_(S)-(+)-2-hexanol
Cosine: 0.99
MS/MS match
Rev. cos prediction
decanoyl-chloride_tetraethylene glycol
Cosine: 0.93
MS/MS match
Rev. cos prediction
3alpha6alpha7alpha_Ala
Cosine: 0.93
MS/MS match
Rev. cos prediction
Chenodeoxycholic-acid_(3-sulfoxy)-pentane-1,5-diamine
Cosine: 0.98
MS/MS match
Rev. cos prediction
3-Hydroxybenzoic acid_agmatine
Cosine: 0.94
MS/MS match
Rev. cos prediction
4-oxopentanoic-acid_butane-1,4-diamine
Cosine: 0.96
MS/MS match
Rev. cos prediction
decanoic-acid_tetraethylene glycol
Cosine: 0.97
MS/MS match
Rev. cos prediction
3-methylbutanoic-acid_4-aminobutanoic acid
Cosine: 0.97
MS/MS match
Rev. cos prediction
2-methylpentanoic-acid_L-arginine
Cosine: 0.96
MS/MS match
Rev. cos prediction
tetradecanedioic-acid_3-methoxy-1-propanol
Cosine: 0.94
MS/MS match
Rev. cos prediction
undecanoic-acid_L-arginine
Cosine: 0.94
MS/MS match
Rev. cos prediction
Aconitic acid_2-methylbutylamine
Cosine: 0.94
MS/MS match
Rev. cos prediction
1-Aminocyclohexanecarboxylic acid_2-methyl-3-pentanol
Cosine: 0.94
MS/MS match
Rev. cos prediction
3keta7alpha_Gly
Cosine: 0.94
MS/MS match
Rev. cos prediction

(E)-2-methylbut-2-enoic-acid_L-arginine

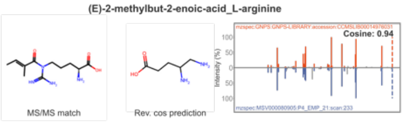


3beta12beta_His

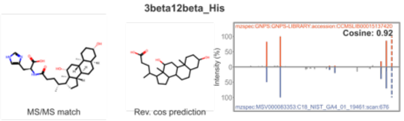


heptanoyl chloride_L-leucine

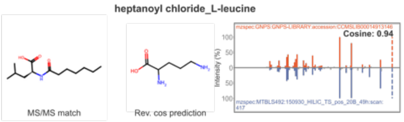


decanoyl chloride_tetraethylene glycol

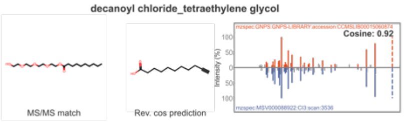


3alpha6alpha_Val

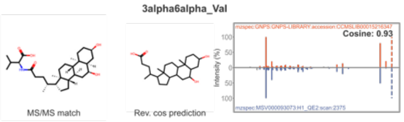


decanoic-acid_L-arginine

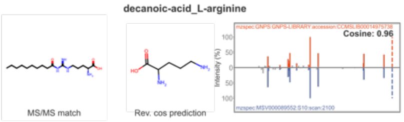


Chenodeoxycholyl_aminophenol

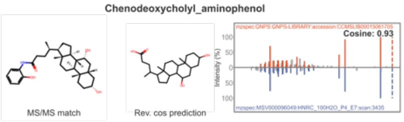


tetradecanedioic-acid_3-methoxy-1-propanol

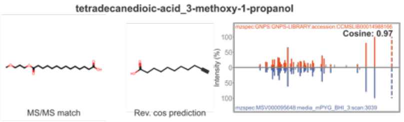


3beta7alpha_Val

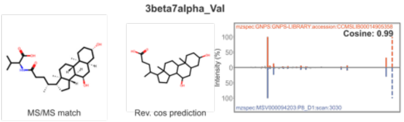


tetradecanedioic-acid_2,3-butanediol

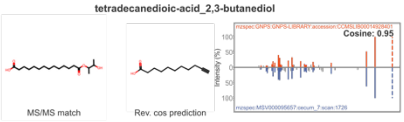


Hyodeoxycholic acid_4-aminobutanoic acid

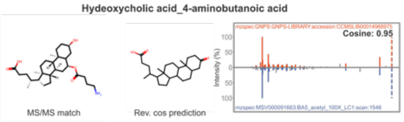


2-hydroxytetradecanoic-acid_1-propanol

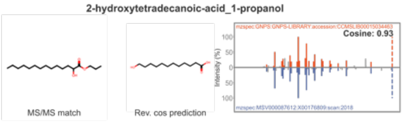


methyl-succinic-acid_L-arginine

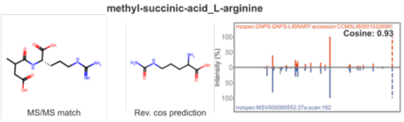


Lactic acid_cadaverine

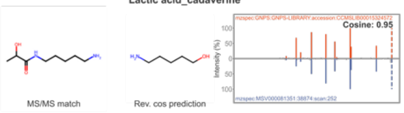


carnitine_linolenic-acid

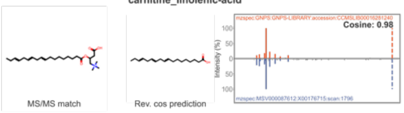


2-methylglutaric acid_phenylethyl-amine

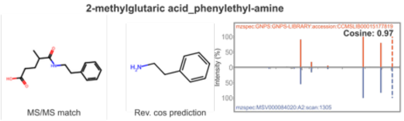


3-methylbutanoic-acid_L-isoleucine

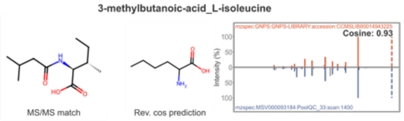


4-oxopentanoic-acid_2-butene-1,4-diol

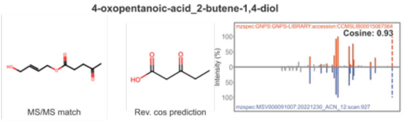


Pyruvic acid_tryptamine

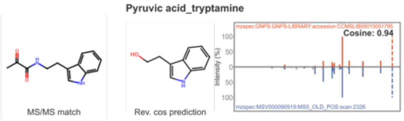


12-methyltridecanoic-acid_N,N-dimethylaminoethanol

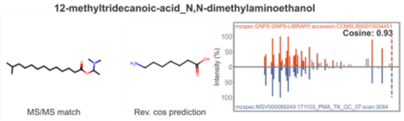


3beta7alpha12alpha_Lys

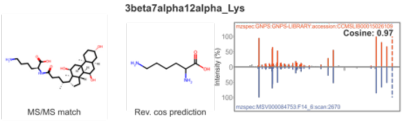


Pyroglutamic acid_L-leucine

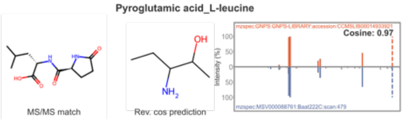


octanoic-acid_L-arginine

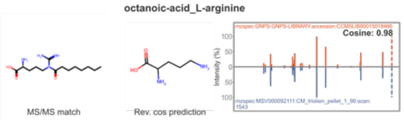


cholic-acid_ethanolamine

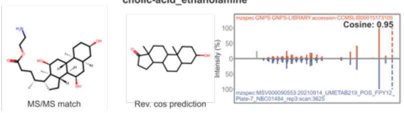


octanoic-acid_L-valine

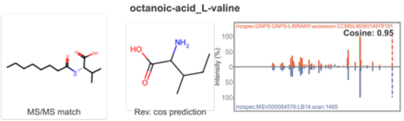


3beta7alpha12alpha_Lys

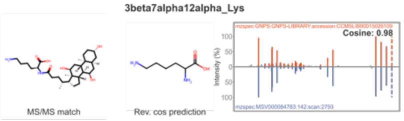


cholic acid_L-phenylalanine

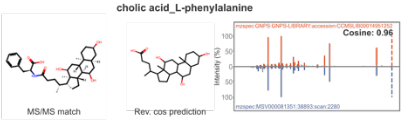


Pentadecanoic acid_2-methylbutylamine

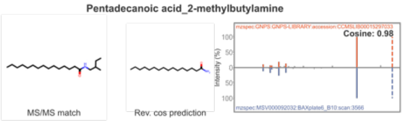


tetradecanedioic-acid_2,3-butanediol

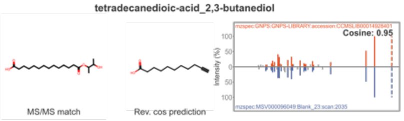


1-hydroxylbuprofen_undecanoic-acid

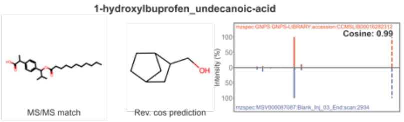


L-tert-leucine_lauryl alcohol

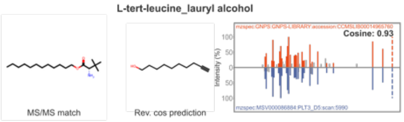


1,5-dimethyl-1h-pyrazole-3-carboxylic-acid_D(+)-galactose

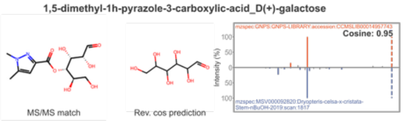


1-Aminocyclohexanecarboxylic acid_(S)-(+)-2-hexanol

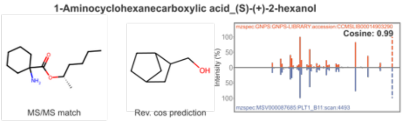

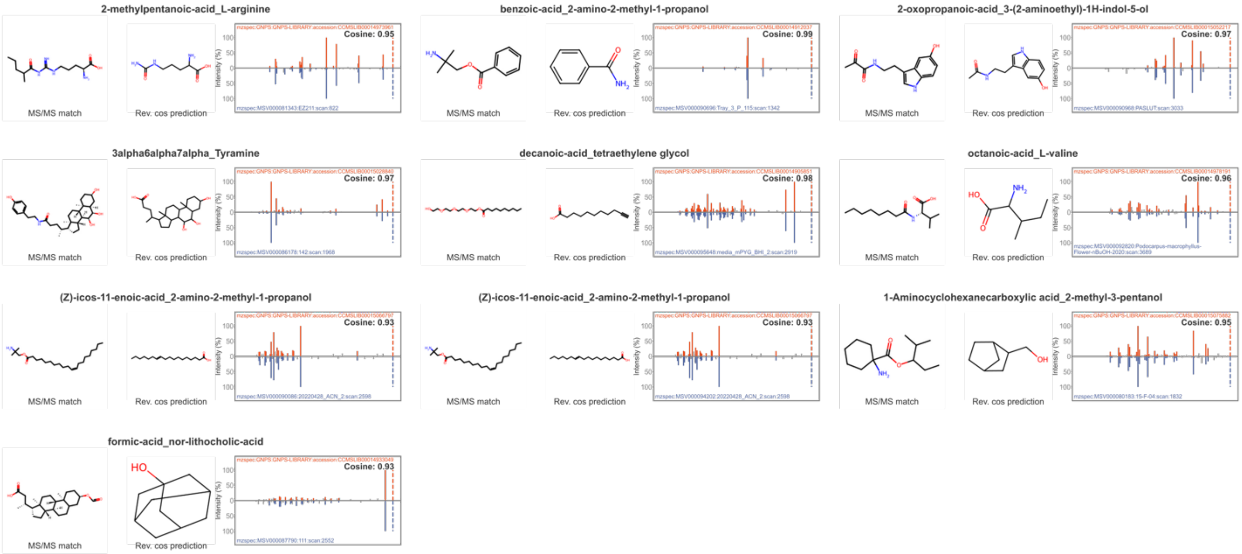
2-methylpentanoic-acid_L-arginine
Cosine: 0.95
benzoic-acid_2-amino-2-methyl-1-propanol
Cosine: 0.99
2-oxopropanoic-acid_3-(2-aminoethyl)-1H-indol-5-ol
Cosine: 0.97
3alpha6alpha7alpha_Tyramine
Cosine: 0.97
decanoic-acid_tetraethylene glycol
Cosine: 0.98
octanoic-acid_L-valine
Cosine: 0.96
(Z)-icos-11-enoic-acid_2-amino-2-methyl-1-propanol
Cosine: 0.93
(Z)-icos-11-enoic-acid_2-amino-2-methyl-1-propanol
Cosine: 0.93
1-Aminocyclohexanecarboxylic acid_2-methyl-3-pentanol
Cosine: 0.95
formic-acid_nor-lithocholic-acid
Cosine: 0.93
MS/MS match
Rev. cos prediction
Intensity (%)
